# Dizocilpine derivatives with neuroprotective effect lacking the psychomimetic side effects

**DOI:** 10.1101/2024.06.17.599304

**Authors:** Jan Konecny, Anna Misiachna, Marketa Chvojkova, Lenka Kleteckova, Marharyta Kolcheva, Martin Novak, Lukas Prchal, Marek Ladislav, Katarina Hemelikova, Jakub Netolicky, Martina Hrabinova, Tereza Kobrlova, Jana Zdarova Karasova, Jaroslav Pejchal, Pavla Jendelova, Yuan-Ping Pang, Karel Vales, Jan Korabecny, Ondrej Soukup, Martin Horak

## Abstract

We aimed to prepare novel dibenzosuberane derivatives that act on N-methyl-D-aspartate (NMDA) receptors with potential neuroprotective effects. Our approach involved modifying the tropane moiety of MK-801, a potent open-channel blocker known for its psychomimetic side effects, by introducing a seven-membered ring with substituted base moieties specifically to alleviate these undesirable effects. Our *in silico* analyses showed that these derivatives should have high gastrointestinal absorption and cross the blood-brain barrier (BBB). Our pharmacokinetic studies in rats supported this conclusion and confirmed the ability of leading compounds **3l** and **6f** to penetrate the BBB. Electrophysiological experiments showed that all compounds exhibited different inhibitory activity towards the two major NMDA receptor subtypes, GluN1/GluN2A and GluN1/GluN2B. Of the selected compounds intentionally differing in the inhibitory efficacy, **6f** showed high relative inhibition (∼90% for GluN1/GluN2A), while **3l** showed moderate inhibition (∼50%). An *in vivo* toxicity study determined that compounds **3l** and **6f** were safe at 10 mg/kg doses with no adverse effects. Behavioral studies demonstrated that these compounds did not induce hyperlocomotion or impair prepulse inhibition of startle response in rats. Neuroprotective assays using a model of NMDA-induced hippocampal neurodegeneration showed that compound **3l** at a concentration of 30 μM significantly reduced hippocampal damage in rats. These results suggest that these novel dibenzosuberane derivatives are promising candidates for developing NMDA receptor-targeted therapies with minimal psychotomimetic side effects.

## 1. Introduction

*N*-methyl-D-aspartate (NMDA) receptors are essential ionotropic glutamate receptors facilitating excitatory synaptic transmission within the mammalian central nervous system (CNS). They are crucial for the pathology of several CNS disorders, including epilepsy, Alzheimer’s disease (AD), and autism [1]. In the adult forebrain, the prevalent forms of NMDA receptors are the diheteromeric GluN1/GluN2A and GluN1/GluN2B, along with the triheteromeric GluN1/GluN2A/GluN2B receptors [1, 2]. The GluN subunits share a common structure: an extracellular N-terminal domain, a ligand-binding domain formed by S1 and S2 segments, a transmembrane domain (TMD), and an intracellular C-terminal domain. The TMD includes three transmembrane-spanning helices (M1, M3, and M4) and a membrane loop (M2), with the M2 loops and M3 helices together, forming the ion channel and the binding site for Mg^2+^, which blocks the receptor in its resting state [1, 3, 4].

Numerous modulators of NMDA receptors operate through competitive, allosteric, or open-channel blocking mechanisms. Notably, the pharmacological modulation of GluN2B-containing NMDA receptors is of particular interest due to their potential effectiveness in treating neurodegenerative diseases such as Alzheimeŕs disease (AD) and Parkinson’s disease as well as epilepsy, pain, and depression [5–10]. However, the selective antagonists acting at GluN2B-containing NMDA receptors exhibited side effects in pre-clinical studies or have not yet achieved clinical success [11–14]. Among the well-recognized NMDA receptor drugs are memantine, ketamine, and dizocilpine (MK-801). While all three are open-channel blockers for GluN1/GluN2 receptors, memantine is approved for AD treatment [15]; ketamine serves as an anesthetic and antidepressant [16, 17]; and MK-801 acts as a model compound in schizophrenia symptom studies [18, 19]. The varied pharmacological impacts of these drugs likely stem from their distinct molecular mechanisms of action. Although memantine and ketamine bind relatively weakly to the ion channel pore, MK-801 is known for its high affinity and near-’irreversible’ binding within the GluN1/GluN2 ion channel pore, resulting in potent effects but also psychomimetic actions, which limit its clinical use [20–25].

This study aims to develop derivatives of MK-801, which would exert lower affinity towards NMDA receptors while maintaining its beneficial efficacy. We have designed a series of 27 novel dibenzosuberane derivatives that demonstrated varying inhibitory activity toward the major NMDA receptor subtypes. Applying an *in vitro* selection process consisting of mainly electrophysiology, cytotoxicity, and *in vivo* pharmacokinetics, the selected compounds were analyzed for *in vivo* side effects and neuroprotection assessment using NMDA-induced hippocampal lesion.

## 2. Results and Discussion

### 3.1 Design

The dibenzosuberane scaffold represents the core of the newly designed derivatives in this study. The compounds were structurally inspired by a previously published series of 9*H*-fluoren-9-amines [26]. In this family, we combined scaffolds of the two well-known NMDA receptor ligands, tacrine, and carbazole, to obtain derivatives with micromolar potency towards the NMDA receptor [27, 28]. The main area of interest in the current series was to improve the affinity and potency of novel derivatives towards NMDA receptors. We took advantage of the structural features known from discovering phencyclidine (PCP) derivatives. Introducing one or more methylene groups between two cyclic moieties resulted in ligand stiffening, which enhanced the affinity for NMDA receptors [29]. Consequently, we incorporated a seven-membered ring into our novel compounds, replacing the traditional five-membered ring. These molecules are structurally similar to MK-801 [30]. However, given its high affinity and slow clearance, MK-801 is associated with severe medicinal risks such as Olney’s lesions and behavioral disturbance, thereby limiting its use in human medicine [20, 21]. Therefore, to mitigate these risks, we altered the tropane moiety of MK-801, changed the ligand topology, and introduced substituted basic moieties (primary or secondary amines) at position 5 on the seven-membered ring (Fig. 1).

**Fig. 1.**
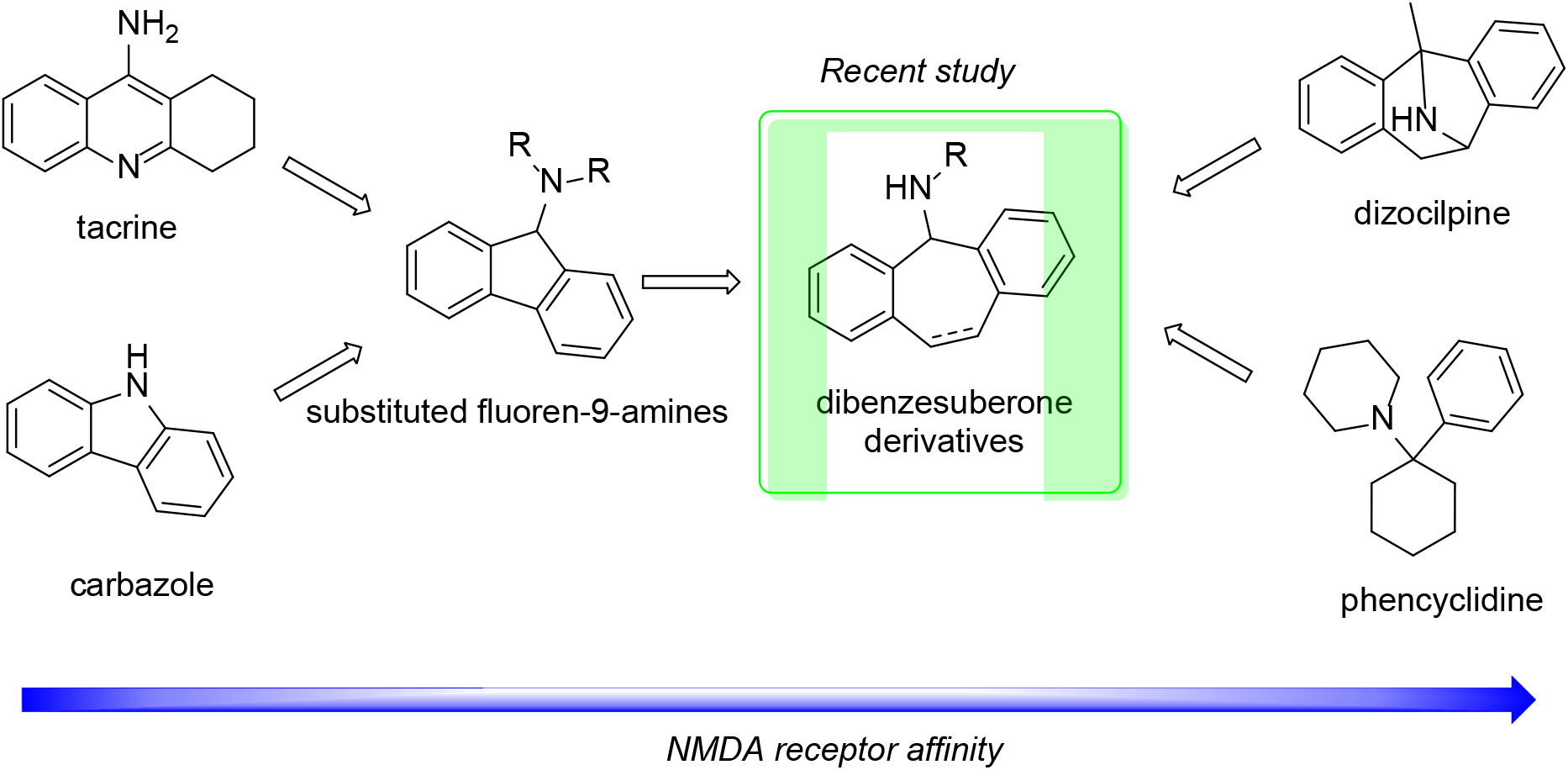
Design of novel dibenzosuberone derivatives possessing activity towards NMDA receptor.

### 3.2. In silico *prediction of CNS and oral availability*

*In silico* absorption, distribution, metabolism, and excretion (ADME) testing provides a robust predictive framework for assessing peroral and CNS availabilities, crucial for targeting neurodegenerative diseases such as AD, where the CNS is the primary focus [31–33]. Additionally, peroral administration is preferred for dementia patients due to its convenience [34, 35]. Thus, we initially screened the designed compounds *in silico* to predict their peroral and CNS availabilities and assessed potential pharmacokinetic profiles, drug-likeness, and ADME characteristics using the web-based tool SwissADME [36–38]. A prototype open-channel blocker of the NMDA receptor, memantine, was used as a reference compound with known CNS status and pharmacokinetic profile (Table 1; also in Tables S1 and S2, Supplementary Information) [39]. We also employed the blood-brain barrier (BBB) permeation score algorithm for CNS prediction, presuming CNS availability with the highest reported predictive score [40]. The results from the BBB score fit between 5-6 (except derivative 6k), indicating the high possibility for the compounds to permeate through the BBB [40]. Table S1 (Supplementary Information) presents the physiological parameters used for BBB score calculation. In general, most of the compounds also fulfill the drug-likeness in various pharmacological models defined by pharmaceutical companies like Vebeŕs (GSK) [41], Lipinskís (Pfizer) [42], Egańs (Pharmacopeia) [43], Ghosés (Amgen) [44], and in some cases Mueggés (Bayer) [45] (results are presented in Table S2, Supplementary Information). In summary, the proposed derivatives showed a high probability of crossing the BBB with good predictive bioavailability and promising pharmacokinetic profile.

**Table 1.** Summary of *in silico* ADME and drug-likeness of derivatives 3a-o and 6a-l with memantine using various prediction models.

| Compound | GIA<sup>a</sup> | BBB<sup>b</sup> | BBB score<sup>c</sup> | Lipinski<sup>d</sup> | Bio. Score<sup>e</sup> |
| --- | --- | --- | --- | --- | --- |
| 3a | High | Yes | 5.63 | Yes | 0.55 |
| 3b | High | Yes | 5.63 | Yes | 0.55 |
| 3c | High | Yes | 5.63 | Yes | 0.55 |
| 3d | High | Yes | 5.62 | Yes | 0.55 |
| 3e | High | Yes | 5.59 | No | 0.55 |
| 3f | High | Yes | 5.59 | No | 0.55 |
| 3g | High | Yes | 5.53 | No | 0.55 |
| 3h | High | Yes | 5.10 | No | 0.55 |
| 3i | High | Yes | 5.52 | Yes | 0.55 |
| 3j | High | Yes | 5.68 | Yes | 0.55 |
| 3k | High | Yes | 5.11 | No | 0.55 |
| 3l | High | Yes | 5.48 | Yes | 0.55 |
| 3m | High | Yes | 5.61 | Yes | 0.55 |
| 3n | High | Yes | 5.14 | No | 0.55 |
| 3o | High | Yes | 5.58 | Yes | 0.55 |
| 6a | High | Yes | 5.33 | Yes | 0.55 |
| 6b | High | Yes | 5.32 | Yes | 0.55 |
| 6c | High | Yes | 5.31 | Yes | 0.55 |
| 6d | High | Yes | 5.30 | Yes | 0.55 |
| 6e | High | Yes | 5.29 | Yes | 0.55 |
| 6f | High | Yes | 5.28 | Yes | 0.55 |
| 6g | High | Yes | 5.23 | No | 0.55 |
| 6h | High | Yes | 5.37 | No | 0.55 |
| 6i | High | Yes | 5.17 | Yes | 0.55 |
| 6j | High | Yes | 5.37 | Yes | 0.55 |
| 6k | High | Yes | 4.82 | Yes | 0.55 |
| 6l | High | Yes | 5.17 | Yes | 0.55 |
| Memantine | High | Yes | 4.61 | Yes | 0.55 |

<sup>a</sup> GIA = Gastrointestinal absorption; <sup>b</sup> BBB = Blood-brain barrier permeation; <sup>c</sup> ref. [40]; <sup>d</sup> ref. [42]; <sup>e</sup> Bio. Score = Bioavailability score [46].

### 3.3. Chemistry

*N*-Substituted 10,11-dihydro-5*H*-dibenzo[*a,d*][7]annulen-5-amines were prepared in one-step synthesis between the commercially available 5-chloro-10,11-dihydro-5*H*-dibenzo[*a,d*][7]annulene (1) with the various primary or secondary amines (2a-o) in dry acetonitrile (Scheme 1). This reaction afforded *N*-substituted-10,11-dihydro-5*H*-dibenzo[*a,d*][7]annulen-5-amines (3a-o; Scheme 1) in mild-to-moderate yields (17 – 66%). The compounds were converted to respective hydrochloride salts.

**Scheme 1.**
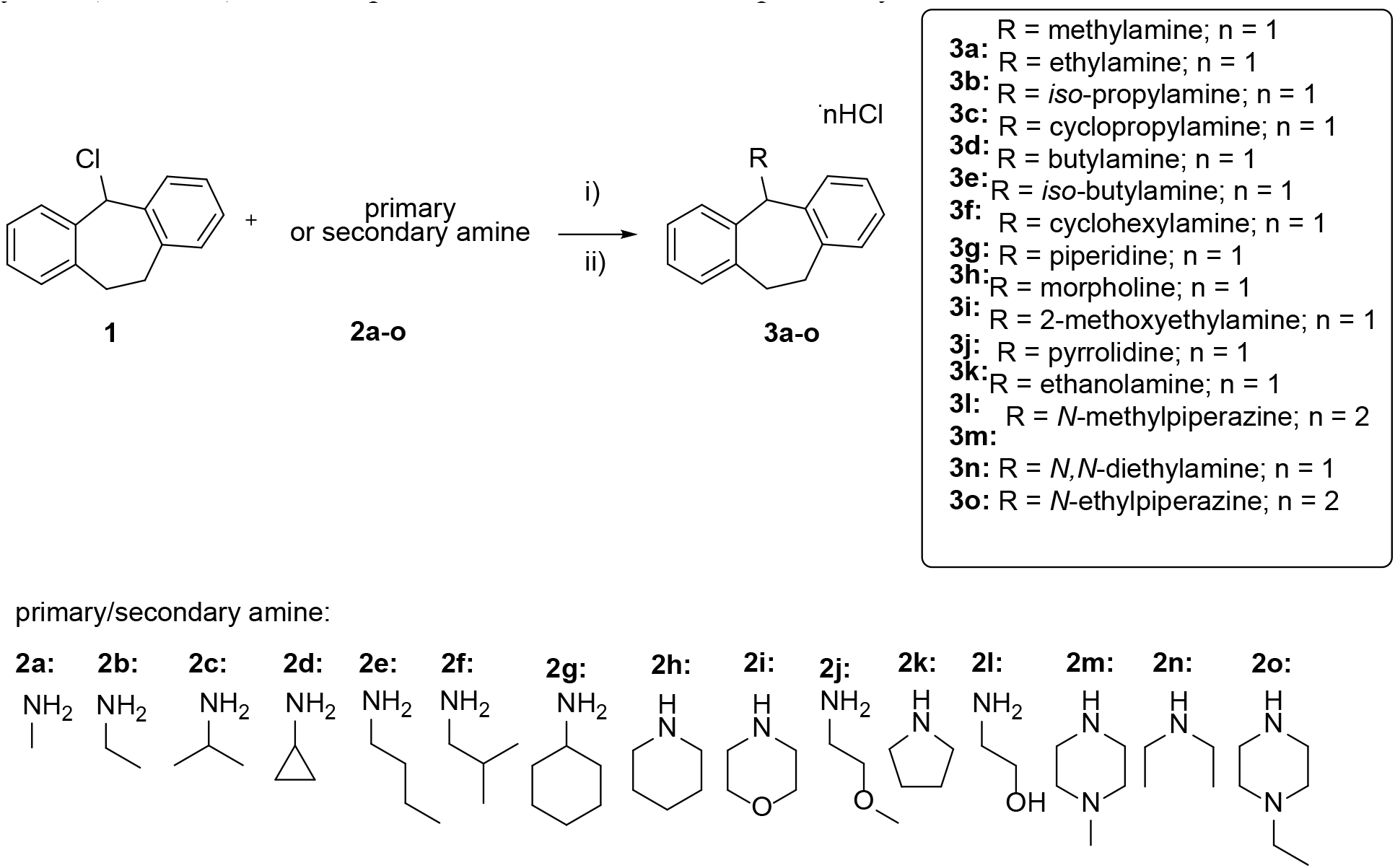
Synthetic procedure for preparing *N*-substituted-10,11-dihydro-5*H*-dibenzo[*a,d*][7]annulen-5-amines 3a-o. Reaction conditions: (i) MeCN, 40 °C, 3 h and (ii) HCl (37% aq.sol.), MeOH, RT, 1 h. *N*-Substituted-5*H*-dibenzo[*a,d*][7]annulen-5-amines (6a-l; Scheme 2) were prepared from the commercially available 5*H*-dibenzo[*a,d*][7]annulen-5-ol (4). Starting material 4 was dissolved in dry DCM and treated with thionyl chloride to provide 5-chloro-5*H*-dibenzo[*a,d*][7]annulene (5), which was used directly for the next reaction (Scheme 2). Final derivatives 6a-l were prepared as hydrochloride salts by the above procedure with moderate-to-good overall yields (41 – 74%). The compounds 3a-o and 6a-l were characterized by ^1^H, ^13^C NMR spectra, melting points, and HRMS analysis. LC analysis confirmed purity for all derivatives >95% (Supplementary Information).

**Scheme 2.**
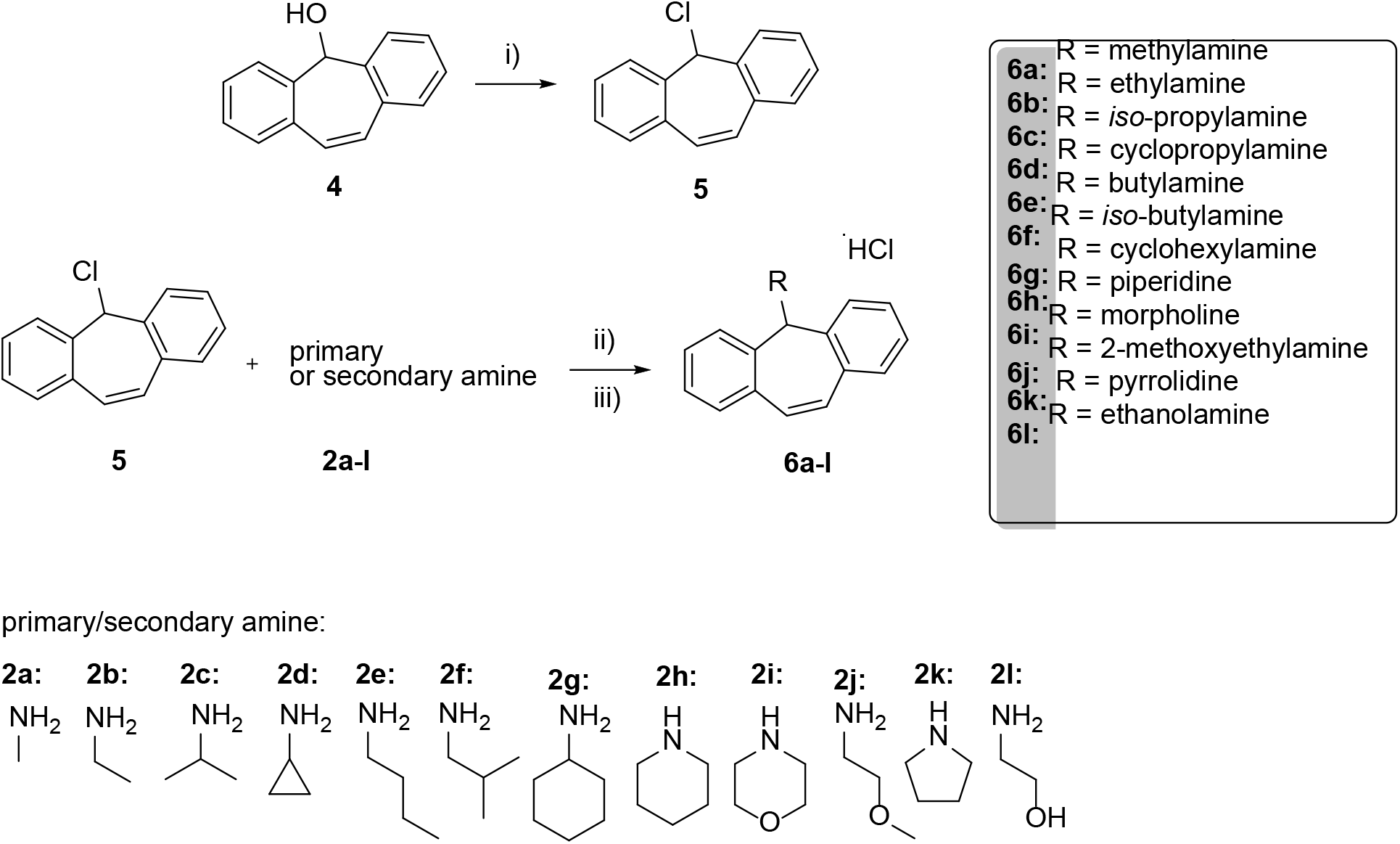
Synthetic procedure for preparing *N*-substituted-5*H*-dibenzo[*a,d*][7]annulen-5-amines 6a-l. Reaction conditions: (i) SOCl_2_, DCM, RT, 12 h; (ii) MeCN, 40 °C, 3 h and (iii) HCl (37% aq.sol.), MeOH, RT, 1 h.

### 3.4. Evaluation of antagonist activity towards NMDA receptor and cytotoxicity

First, we determined the relative inhibition of all novel compounds on the two major diheteromeric NMDA receptor subtypes in the adult cortex – GluN1/GluN2A and GluN1/GluN2B [1] – using a clinically relevant concentration (3 µM). We kept the measured cells at a membrane potential of -60 mV, and the values of relative inhibition by the compounds were calculated from the L-glutamate-induced responses of human versions of the GluN1/GluN2A (hGluN1/hGluN2A) and GluN1/GluN2B (hGluN1/hGluN2B) receptors obtained under steady-state conditions. These experiments showed that all the studied compounds inhibit the responses of both NMDAR subtypes (see Table 2). Specifically, in the case of hGluN1/hGluN2A receptors, we obtained relative inhibition values ranging from ∼5% for compound **3h** to ∼93% for compound **6f**, while for hGluN1/hGluN2B receptors, we measured values ranging from ∼3% for compound **3i** to ∼92% for compound **6g** at a fixed dose of 3 µM of the tested compound. These results showed that our series of synthesized compounds contain potent inhibitors of both NMDAR subtypes. From a structure-activity point of view, derivatives **e**, **f**, and **g** in both the **3** and **6** series representing butylamine, iso-butylamine, and cyclohexylamine substituents, respectively, showed the most pronounced effect on both NMDAR subtypes.

**Table 2.** Inhibitory effect of synthesized compounds at human GluN1/GluN2A and GluN1/GluN2B receptors expressed in the HEK293 cells and their cytotoxicity measured using CHO-K1 cells.

| Compound | hGluN1/hGluN2A<br>$\text{RI}^a$ (%) ± SEM | hGluN1/hGluN2B<br>$\text{RI}^a$ (%) ± SEM | n (hGluN1/hGluN2A<br>/hGluN1/hGluN2B) | Cytotoxicity<br>CHO-K1<br>$\text{IC}_{50} \pm \text{SEM}$ ($\mu\text{M}$) |
| --- | --- | --- | --- | --- |
| **3a** | 82.34 ± 3.25 | 70.97 ± 1.75 | 5/5 | 409.65 ± 4.85 |
| **3b** | 34.05 ± 4.60 | 51.84 ± 3.39 | 4/5 | 364.07 ± 25.58 |
| **3c** | 52.23 ± 5.23 | 76.92 ± 3.57 | 4/7 | 263.20 ± 11.30 |
| **3d** | 40.11 ± 3.42 | 53.57 ± 3.16 | 10/6 | 177.00 ± 6.60 |
| **3e** | 76.43 ± 4.03 | 85.79 ± 2.19 | 6/7 | 127.57 ± 7.55 |
| **3f** | 76.79 ± 2.99 | 91.62 ± 0.99 | 6/5 | 92.38 ± 6.07 |
| **3g** | 79.42 ± 2.06 | 85.95 ± 3.13 | 5/5 | 85.00 ± 6.94 |
| **3h** | 5.40 ± 1.39 | 5.20 ± 1.56 | 5/5 | 73.78 ± 1.83 |
| **3i** | 6.78 ± 1.67 | 3.43 ± 1.33 | 5/5 | 95.25 ± 1.91 |
| **3j** | 34.90 ± 4.80 | 66.63 ± 3.90 | 5/4 | 384.90 ± 3.40 |
| **3k** | 49.80 ± 3.27 | 62.14 ± 3.44 | 5/4 | 97.68 ± 11.78 |
| **3l** | 47.96 ± 5.18 | 49.67 ± 4.47 | 4/4 | 719.95 ± 31.35 |
| **3m** | 9.70 ± 2.41 | 9.49 ± 2.17 | 5/5 | 121.50 ± 7.30 |
| **3n** | 68.43 ± 4.84 | 75.00 ± 3.14 | 4/6 | 86.41 ± 3.18 |
| **3o** | 7.64 ± 0.84 | 5.22 ± 1.49 | 4/6 | 146.80 ± 5.05 |
| **6a** | 54.32 ± 3.11 | 68.93 ± 3.57 | 4/7 | 299.35 ± 41.25 |
| **6b** | 39.93 ± 4.78 | 59.85 ± 3.15 | 11/6 | 497.90 ± 3.90 |
| **6c** | 75.62 ± 2.36 | 76.05 ± 2.22 | 6/6 | 250.85 ± 0.45 |
| **6d** | 45.00 ± 3.55 | 57.18 ± 3.27 | 6/4 | 260.50 ± 3.80 |
| **6e** | 81.12 ± 2.62 | 79.19 ± 2.84 | 6/5 | 100.75 ± 1.96 |
| **6f** | 93.17 ± 1.11 | 91.77 ± 0.71 | 6/5 | 101.75 ± 9.27 |
| **6g** | 92.12 ± 1.48 | 92.37 ± 1.82 | 6/5 | 81.29 ± 9.35 |
| **6h** | 29.85 ± 2.73 | 58.40 ± 4.37 | 6/5 | 380.56 ± 16.35 |
| **6i** | 31.90 ± 4.16 | 57.65 ± 3.47 | 7/6 | 164.70 ± 13.50 |
| **6j** | 35.63 ± 1.09 | 60.00 ± 4.39 | 4/7 | 390.45 ± 19.05 |
| **6k** | 48.56 ± 2.99 | 85.53 ± 0.21 | 6/4 | >900 |
| **6l** | 38.17 ± 3.80 | 48.78 ± 4.13 | 7/4 | 332.70 ± 7.60 |

<sup>a</sup>Relative inhibition (RI) was calculated as the ratio of steady-state currents evoked by 1 mM L-glutamate in the presence and absence of a 3 μM of the tested compound (multiplied by 100) at a membrane potential of -60 mV. Further details are described in the text. Data are presented as mean ± SEM; n = number of measured HEK293 cells. For cytotoxicity data, n=3.

The open-channel blockers of the NMDA receptors, such as ketamine and memantine, have been shown to have positive clinical effects in the treatment of depression and AD, but other open-channel blockers, such as MK-801, have significant negative side effects [1, 20, 21, 23]. It can be assumed that the positive and negative effects of a given inhibitor of the NMDA receptor *in vivo* are caused by their affinity and the kinetics of inhibition, in addition to their availability in the CNS. Therefore, for further electrophysiological characterization and *in vivo* testing, we selected the **6f** compound, which showed ˃90% (high) relative inhibition on both studied NMDA receptor subtypes, and the **3l** compound, which showed ∼50% (medium) relative inhibition values under the same conditions. In further experiments, we generated concentration-response curves for **6f** (0.03-10 µM) and **3l** (0.3-300 µM) on both hGluN1/hGluN2A and hGluN1/hGluN2B receptors at negative (-60 mV) and positive (40 mV) membrane potentials (Fig. 2A-D). Our analysis showed that the **6f** compound exhibited IC_50_ values ranging from ∼0.2 to ∼0.4 µM at negative membrane potential, whereas IC_50_ values were approximately two orders of magnitude higher at positive membrane potential. Consistent with our data shown in Table 2, we observed that the **3l** compound inhibited both NMDA receptor subtypes at negative membrane potential with IC_50_ values of ∼4 µM, while at positive membrane potential, IC_50_ values were >200 µM (results summarized in Table 3). These results showed that both compounds are voltage-dependent inhibitors with different potencies at the studied NMDA receptors.

**Fig. 2.**
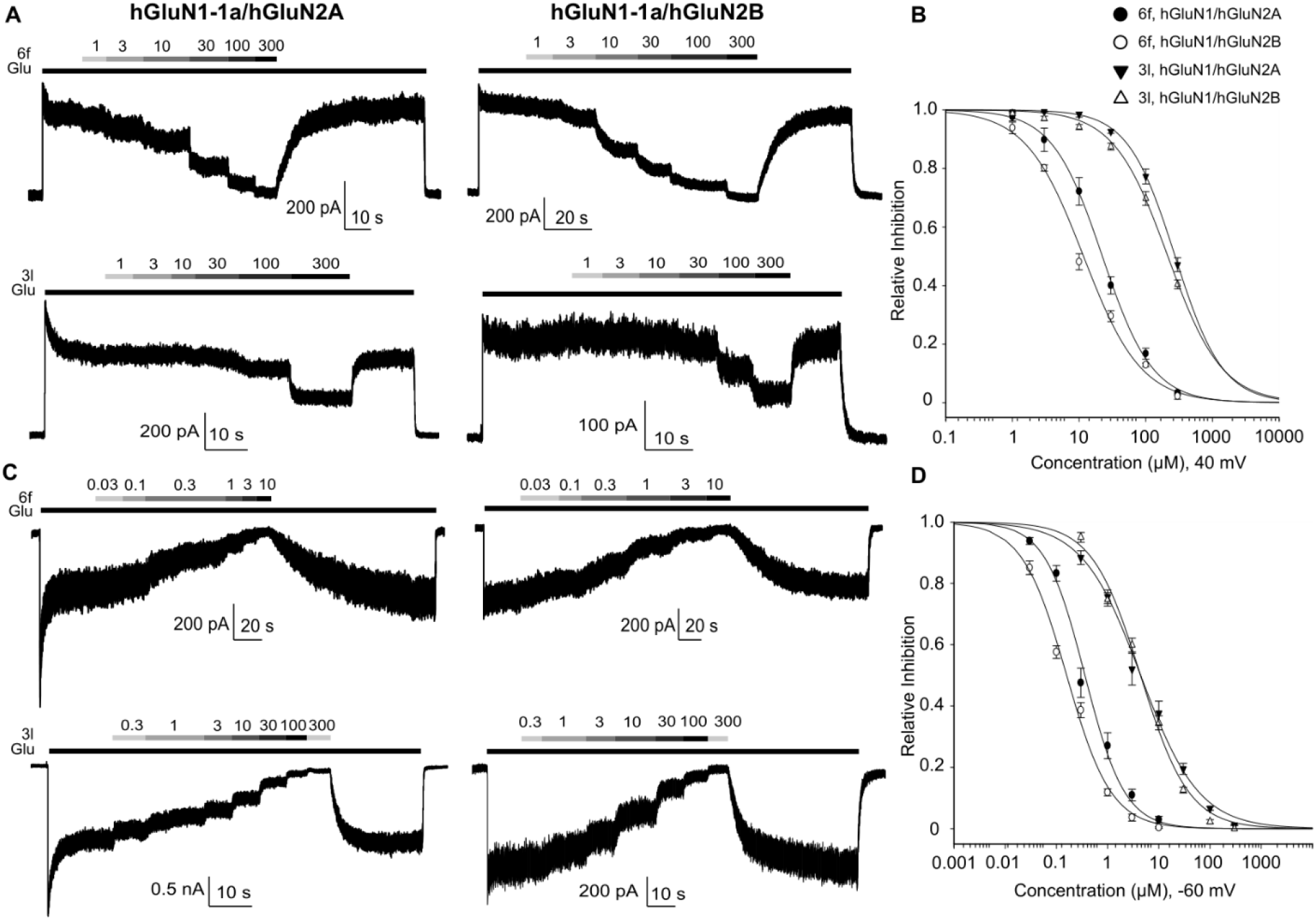
Compound **6f** is a more potent voltage-dependent inhibitor of hGluN1/hGluN2A and hGluN1/hGluN2B receptors than compound **3l**. (A, C) Representative whole-cell voltage-clamp recordings of HEK293 cells expressing hGluN1/hGluN2A or hGluN1/hGluN2B receptors showing the inhibitory effects of compound **6f** (0.03 - 300 µM) and compound **3l** (0.3 - 300 µM) at membrane potentials of 40 mV (A) and -60 mV (C). Glu = 1 mM L-glutamate. (B, D) Normalized inhibitory concentration-response curves for compound **6f** and compound **3l** for hGluN1/hGluN2A and hGluN1/hGluN2B receptors at membrane potentials of 40 mV (B) and -60 mV (D). Concentration-response curves were fitted using *Equation 1*; the resulting IC_50_ values, Hill coefficients, and the numbers of cells measured for each membrane potential are shown in Table 3.

**Table 3.** Concentration-dependent analysis of the inhibitory effect of 6f and 3l at human GluN1/GluN2A and GluN1/GluN2B receptors measured at different membrane potentials.

| Compound | Receptors | IC<sub>50</sub> (-60 mV)<br>(μM) | h (-60 mV) | IC<sub>50</sub> (40 mV)<br>(μM) | h (40 mV) | n (-60/40<br>mV) |
| --- | --- | --- | --- | --- | --- | --- |
| **6f** | hGluN1/hGluN2A | 0.36 ± 0.06 | 1.12 ± 0.11 | 22.37 ± 3.43 | 1.16 ± 0.07 | 4/7 |
| **6f** | hGluN1/hGluN2B | 0.16 ± 0.01 | 1.00 ± 0.05 | 11.67 ± 1.23 | 1.01 ± 0.05 | 7/10 |
| **3l** | hGluN1/hGluN2A | 4.42 ± 0.90 | 0.76 ± 0.02 | 276.46 ± 20.65 | 1.23 ± 0.07 | 4/5 |
| **3l** | hGluN1/hGluN2B | 4.40 ± 0.42 | 0.92 ± 0.03 | 214.75 ± 12.60 | 1.07 ± 0.08 | 9/9 |

In the next phase, we aimed to examine the kinetics of inhibition induced by **6f** and **3l** at both NMDA receptor subtypes. Similar to our previous study [47], we determined the time constants of onset (τ_on_) and offset (τ_off_) of inhibition and induced by different concentrations of **6f** (1, 10, 100 µM) and **3l** (10, 100, 300 µM), measured in the steady-state conditions of the current responses induced by 1 mM L-glutamate and 100 µM glycine at a holding potential of -60 mV (Fig. 3A-B). Using one- or two-parameter exponential fits, we discovered that the compound **6f** exhibited significantly slower τ_off_ values compared to those of the **3l** compound at both NMDA receptor subtypes, although no significant differences were observed in τ_on_ values. (Fig. 3C-D). This indicates that the two compounds might exert distinct pharmacological effects *in vivo*. A relevant comparison can be made with dizocilpine (MK-801), known for its potent psychomimetic properties, which are attributed to its high affinity and nearly irreversible blockade of NMDA receptors [18, 24].

**Fig. 3.**
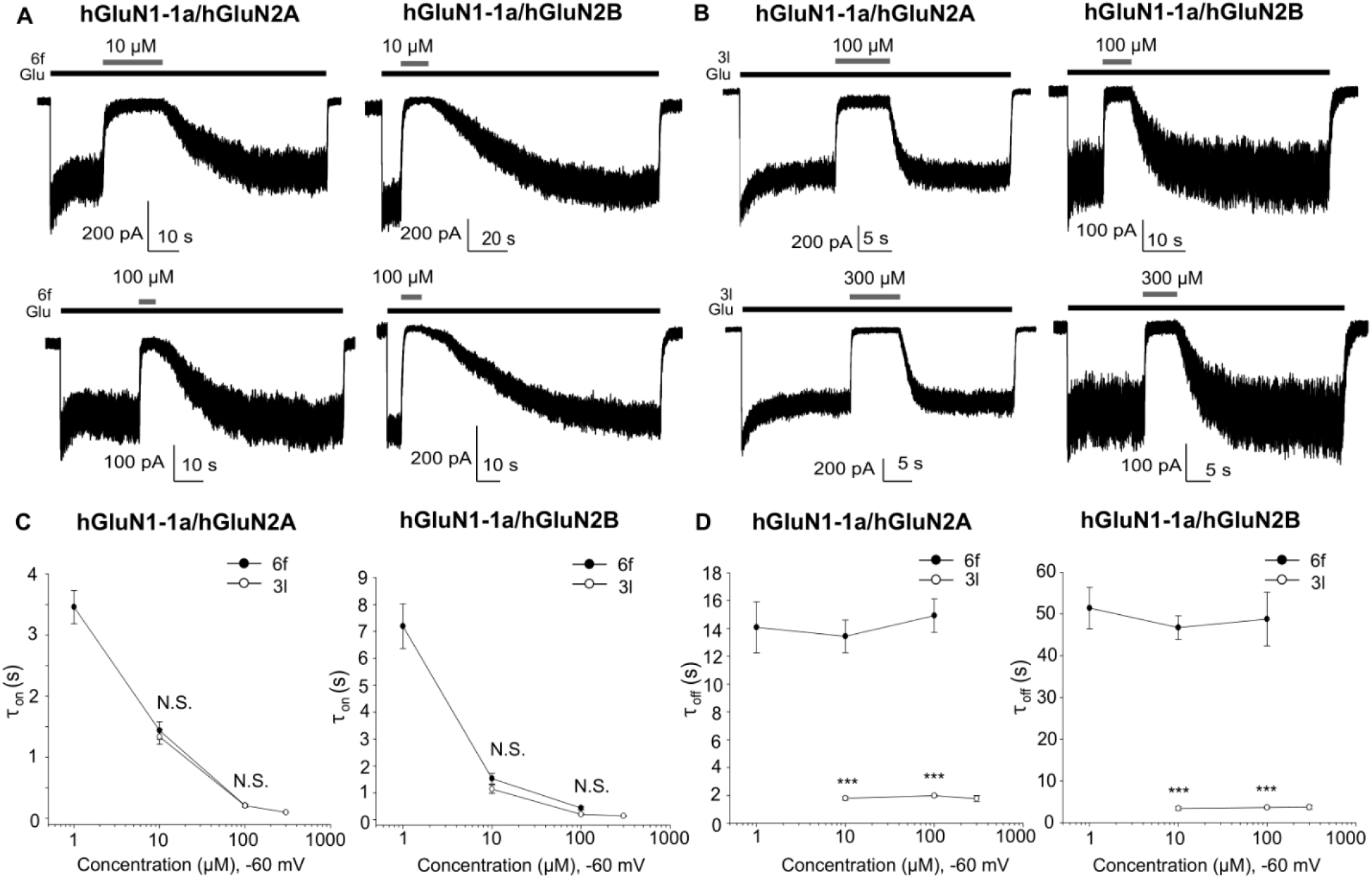
Derivative **6f** exhibits a markedly slower inhibition offset than **3l** at hGluN1/hGluN2A and hGluN1/hGluN2B receptors. (A, B) Representative whole-cell voltage-clamp recordings of HEK293 cells expressing hGluN1/hGluN2A or hGluN1/hGluN2B receptors showing inhibition onset and offset of 10 and 100 µM **6f** (A) and 100 and 300 µM **3l** (B) at a membrane potential of -60 mV. Glu = 1 mM L-glutamate. (C, D) Summary of calculated onset (τ_on_; G) and offset (τ_off_; H) time constants for **6f** and **3l** inhibition of hGluN1/hGluN2A or hGluN1/hGluN2B receptors obtained using *Equation 2* (***p < 0.001, t-test, n ≥ 5).

Next, we analyzed the safety profile of the compounds on the eukaryotic cell line CHO-K1 using 3-[4,5-dimethylthiazol-2-yl]-2,5-diphenyltetrazolium bromide (MTT) (Table 2). IC_50_ values in the tens to thousands of µM generally relate to the intermediate cytotoxicity of the tested compounds. Compounds **3l** and **6k** were identified as the least cytotoxic compounds with 720 and >900 µM values, respectively. Compound **3l** carries a hydrophilic ethanolamine moiety, which could ensure its low cell permeability and low cytotoxicity [48]. On the other hand, compound **6k** bears more lipoplilic pyrrolidine moiety, lowering its solubility. Indeed, we could not determine the IC_50_ value due to its limited solubility in an aqueous solution, but the highest tested concentration (900 µM) was not sufficient to reduce cell viability to 50% and other artifacts like plastic wall sticking should be considered.

Finally, we have also evaluated the ability to inhibit human cholinesterases (AChE and BuChE). Applied Ellmańs method revealed only weak inhibitory activity in cyclohexylamine (**3g** and **6g**) (IC_50_=14.4 ± 1.3 µM, IC_50_=8.5 ± 0.4 µM respectivelly), and piperidine (**6h**) (IC_50_=16.5 ± 0.9 µM) derivatives on BuChE was found. The other compound did not show any inhibitory activity on cholinesterases in tested concentrations (up to 100 µM).

Based on our *in vitro* electrophysiological data, we selected compounds **3g**, **6g**, and **6f,** which showed high (∼90%) relative inhibition at both NMDA receptor subtypes studied, for the next phase of *in vivo* investigation. We further selected compound **3l**, which exhibited moderate (∼50%) inhibition at both NMDA receptor subtypes studied, due to the possibility that strong inhibitor potency to the NMDA receptor may lead to psychomimetic side effects *in vivo*. Compounds **3l** and **6f** with an aliphatic side chain also showed the lowest cytotoxic effect in the respective class of moderate to high potency compounds, which was not the case for compounds **3g** and **6g**, highly potent inhibitors of the NMDA receptors bearing a lipophilic cyclohexylamine moiety.

### 3.5. In vivo pharmacokinetic and toxicity study

The four selected compounds were first evaluated for their ability to cross the BBB, a key property of drugs targeting CNS disorders. A rat model was used for the pilot pharmacokinetic evaluation, and the compounds were evaluated at two time points, namely at 15 and 60 minutes after intraperitoneally (*i.p.*) administration of the 5 mg/kg dose (Fig. 4). The data obtained confirmed the ability of all the tested compounds to cross the BBB, although we observed slight differences between individual compounds. Specifically, the best brain penetration was observed for **3l**, while the other 3 compounds showed the same pattern and rapid elimination from the brain. Due to the higher cytotoxicity and lower solubility (data not shown) of compounds **3g** and **6g**, we decided to proceed further with compounds **3l** and **6f** as representatives of compounds with moderate (**3l**) or high (**6f**) potency for NMDA receptors, in the following behavioral and neuroprotective assays *in vivo*. However, before these studies, we assessed the no-observed-adverse-effect dose level (NOAEL). Both compounds **3l** and **6f** were safe at 10 mg/kg, whereas a decrease in respiration and activity loss was observed at 20 mg/kg. Therefore, the NOAEL was set as 10 mg/kg.

**Fig. 4.**
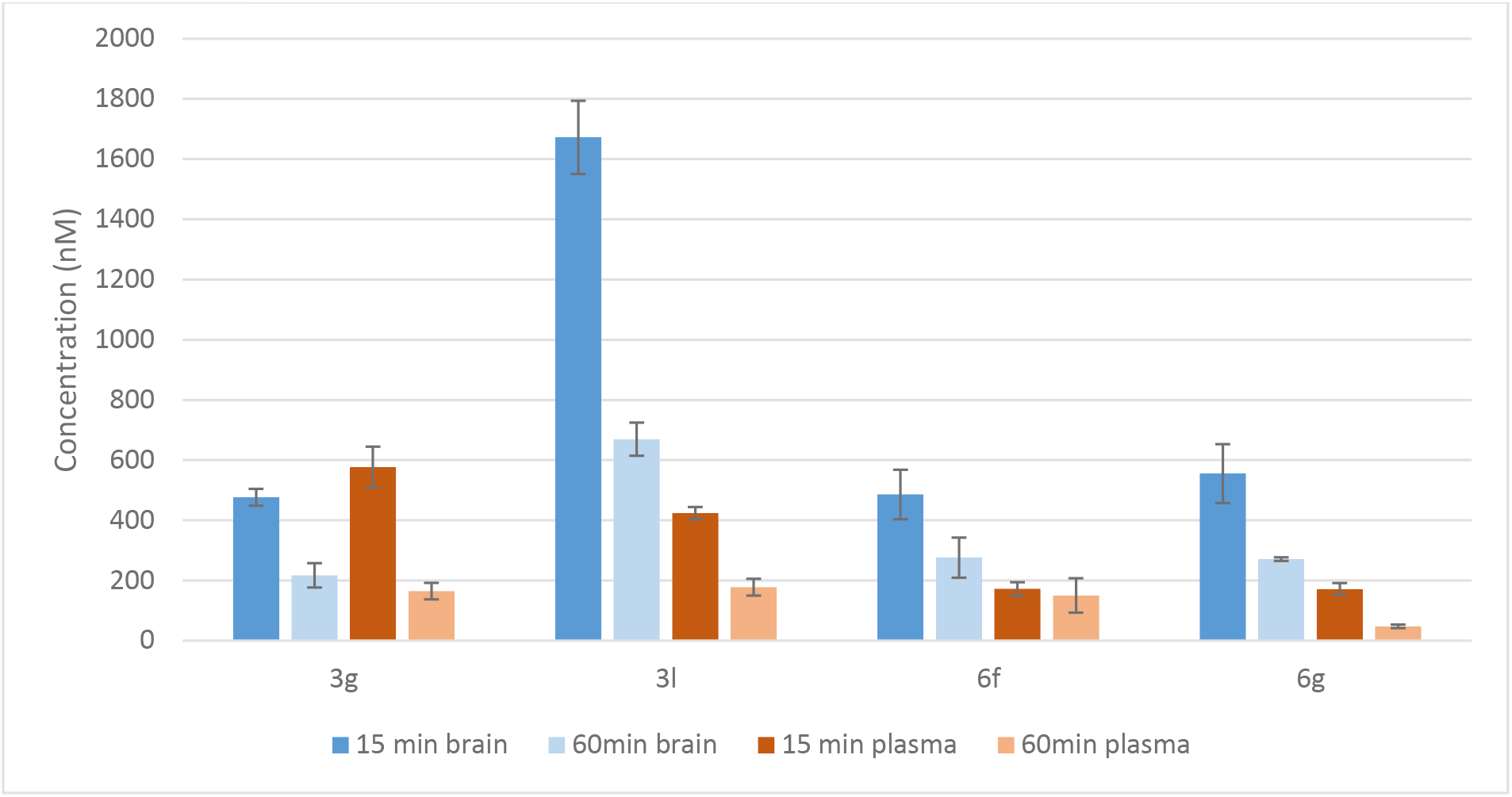
The concentration of the selected compounds in plasma and brain at the 15 and 60 minute intervals after 5 mg/kg *i.p.* in rats. Results are shown as mean ± SEM (n=3).

### 3.6. Evaluation of behavioral side effects of **3l** and **6f**

As noted above, an important limitation of the clinical use of NMDA receptor antagonists is their potential induction of severe psychotomimetic side effects, which in laboratory rodents manifest as hyperlocomotion, impaired prepulse inhibition of the startle response, and other behavioral changes [49–51]. Considering our *in vitro* data and ability to cross the BBB and NOAEL dose of 10 mg/kg, we next evaluated the effects of the compounds **3l** and **6f** at two doses, 1 and 10 mg/kg (administered *i.p.*) on locomotion in the open field test and on prepulse inhibition of startle response in rats. The prototypical high-affinity “irreversible” open-channel blocker of the NMDA receptor, MK-801, was used as a comparator drug. First, analysis of the distance that laboratory rats moved in the open-field test for 10 min showed that 0.2 mg/kg MK-801 induced significant hyperlocomotion; in contrast, the tested compounds **3l** and **6f** did not affect the locomotor activity of the rats, at either the 1 or 10 mg/kg dose used (Fig. 5A). Similarly, in the test of the prepulse inhibition of the acoustic startle response, we observed that 0.3 mg/kg MK-801 induced a deficit in prepulse inhibition, whereas compounds **3l** and **6f** at 1 or 10 mg/kg did not impair prepulse inhibition (Fig. 5B). In conclusion, we did not detect undesirable psychotomimetic effects of **3l** and **6f** at significantly higher concentrations compared with MK-801 results, which showed marked psychotomimetic effects in agreement with the literature [49–51].

**Fig. 5.**
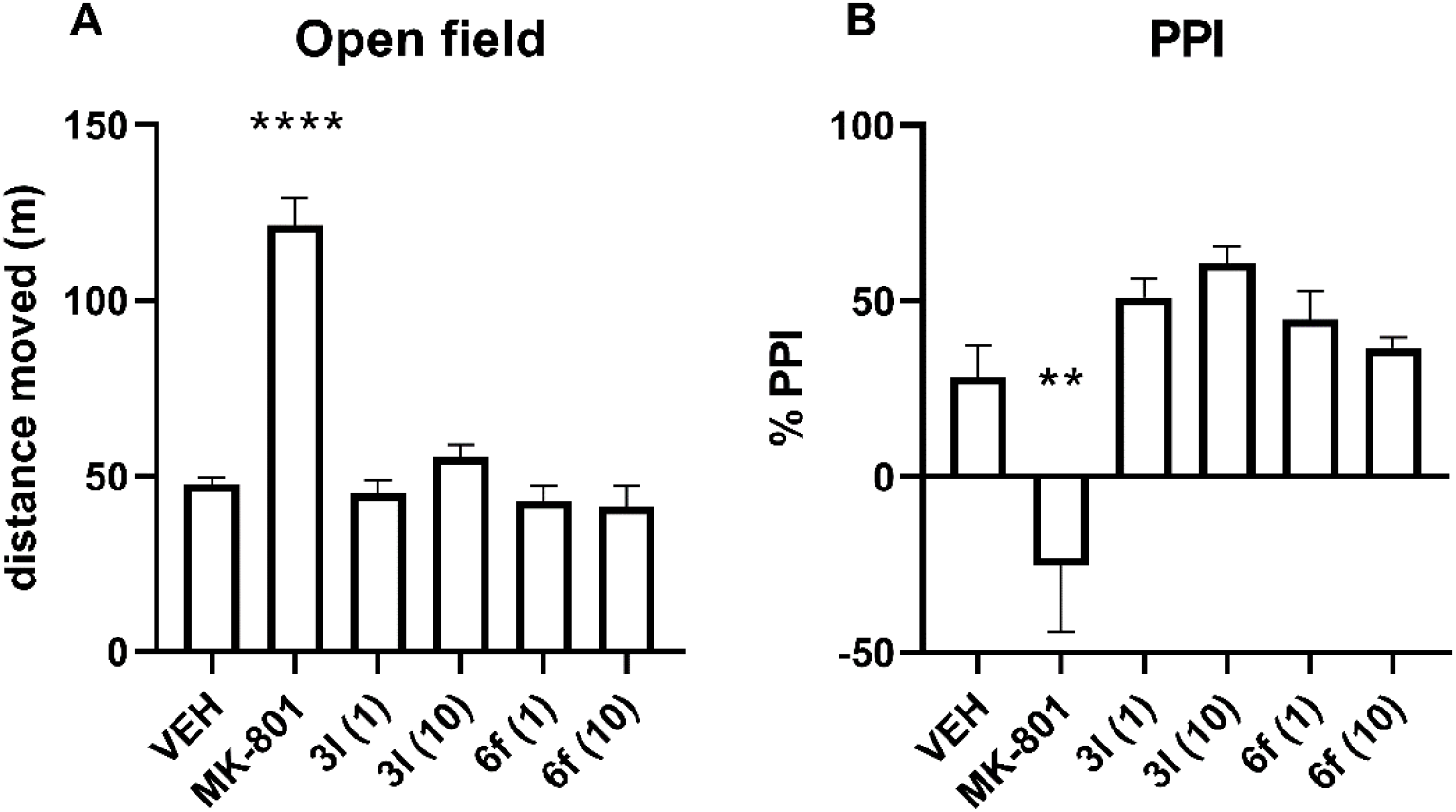
The effects of **3l**, **6f** (1 and 10 mg/kg), and MK-801 on the locomotor activity of rats in the open field (A) and on the prepulse inhibition of startle response (PPI; B). The compounds **3l** and **6f** did not induce hyperlocomotion or deficit of the prepulse inhibition, indicating a low risk of psychotomimetic side effects. Conversely, MK-801 (0.2 mg/kg for open field and 0.3 mg/kg for PPI) disrupted both these parameters. * vs. vehicle (VEH), ** p < 0.01, **** p < 0.0001; ANOVA and Dunnett’s multiple comparisons test, F (5, 32) = 43.32, p < 0.0001 for open field test, F (5, 31) = 9.208, p < 0.0001 for prepulse inhibition test.

### 3.7. In vivo neuroprotective effect

Finally, we investigated whether the selected drugs exhibit neuroprotective effects in a model of hippocampal lesion induced by the application of 25 mM NMDA to the rat dorsal hippocampus, leading to excessive activation of NMDA receptors and subsequent extensive and long-lasting damage of CA1, CA3, gyrus dentatus and hilus in the affected hippocampus and surrounding neural tissue [52]. Our previous studies have demonstrated that this model is a suitable screening method to assess the potential neuroprotective effect of novel compounds acting at the NMDA receptors [53–55]. The total hippocampal damage score analysis revealed increased scores in NMDA-treated rats compared to control animals. Rats co-administered with NMDA and 30 µM **3l** showed significantly lower damage scores, whereas no significant neuroprotective effect was demonstrated in the case of **6f** (Fig. 6A-B).

**Fig. 6.**
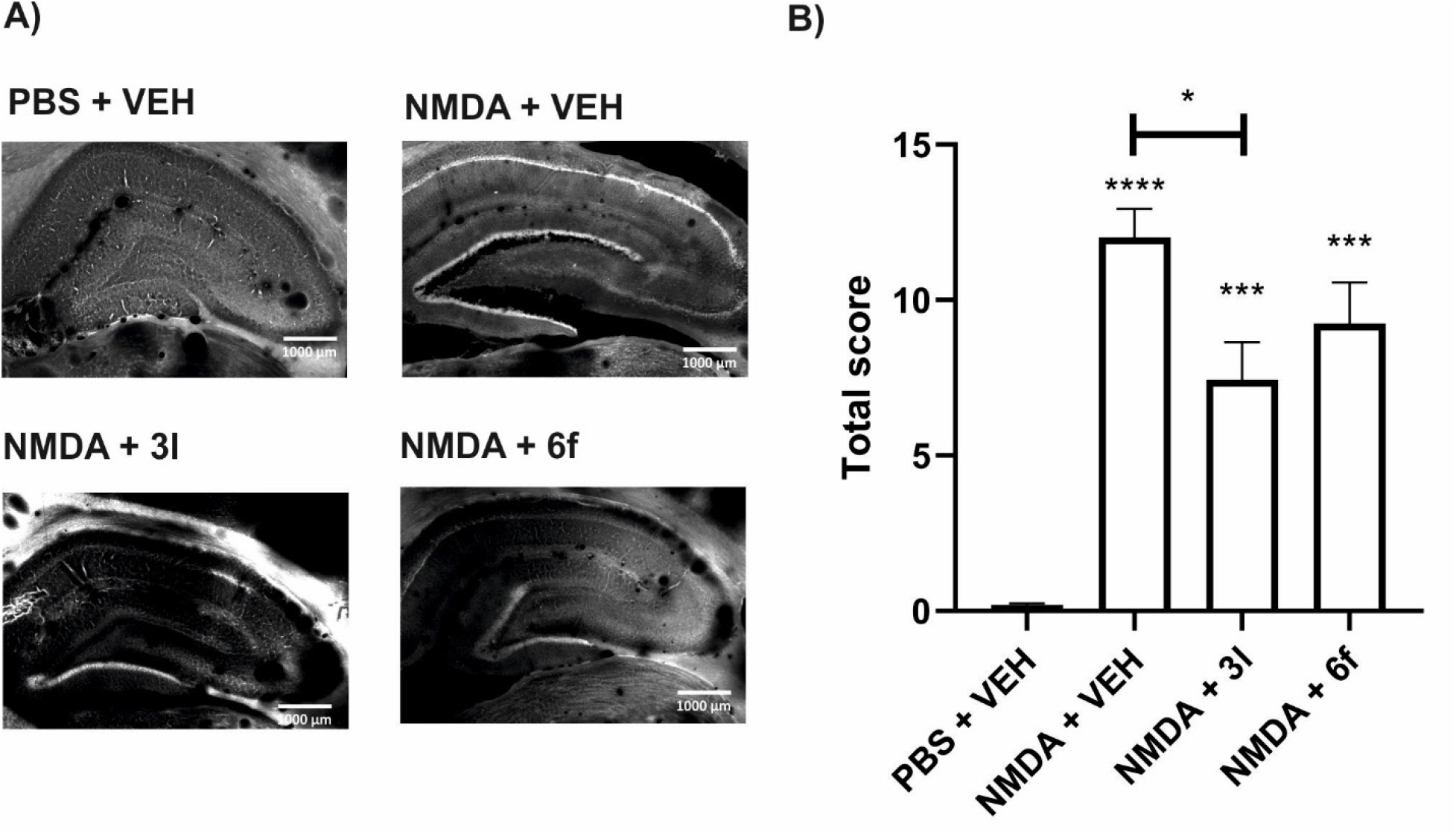
The neuroprotective effect of **3l** and **6f** in the NMDA-induced model of hippocampal neurodegeneration. (A) Representative images of brain slices from rats administered the solutions containing 25 mM NMDA and vehicle (VEH), 30 μM **3l** or **6f** into the dorsal hippocampus. (B) The total scores of the hippocampal damage for the indicated co-applications. * vs. PBS + VEH, *p < 0.05, ***p < 0.001, ****p < 0.0001; W (3.000, 13.14) = 76.62, p < 0.0001; Welch’s ANOVA with Dunnett’s T3 multiple comparisons test.

The efficacy of **3l** and inefficacy of **6f** in our neuroprotective assay is surprising, as **3l** shows approximately half the efficacy in inhibiting both NMDA receptor subtypes at a concentration of 3 µM (Table 2), and the IC_50_ value for hGluN1/hGluN2B receptors (which are associated with neurodegeneration [7] is approximately 27-fold higher (Table 3). The distribution to the brain (Fig. 4) is irrelevant here because both compounds were administered intrahippocampally. However, the high (30 μM) concentration may have erased the differences in inhibitory potency of **3l** and **6f**, as the dose is simply sufficient to elicit the desired neuroprotective effect. Indeed, we observed a (nonsignificant) dose-dependent trend toward hyperlocomotion in the open field (Fig. 5A), suggesting a higher activity of compound **3l**. This suggests that characteristics beyond chemical structure, such as the kinetics of inhibition at NMDA receptors, may play a crucial role in the clinical expression of the compounds’ effects. Notably, both compounds exhibit similar τ_on_ values, but τ_off_ is significantly slower for **3l** compared to **6f**, making compound **3l** wash out faster (Fig. 3). This is corroborated by the irreversible nature of NMDA receptor inhibition by MK-801, which at the behavioral level results in more adverse effects compared to the NMDA receptor antagonist memantine, which has faster τ_off_ kinetics [22–24, 51, 56]. This is further supported by the fact that the activation of extrasynaptic, but not synaptic, NMDA receptors is associated with neurodegeneration [57]. In theory, open-channel blockers with different offset kinetics may distinctly block synaptic (phasic) over extrasynaptic (tonic) NMDA receptors, leading to different neuroprotective effects. In addition, affinity for other targets may also influence the manifestations of the compound. We have recently reported that the inhibitory effect of acetylcholinesterase (AChE) may be important and exacerbate the beneficial effect of compounds acting at NMDA receptors [55]. However, neither **3l** nor **6f** inhibited AChE or butyrylcholinesterase (BChE), suggesting they do not interfere with the cholinergic system. In conclusion, the studied compounds **3l** and **6f** at both doses showed a promising profile consistent with the low risk of adverse behavioral side effects typical of some other NMDA receptor antagonists.

## 3. Conclusion

Our primary goal was to discover novel compounds with high neuroprotective potential and low risk of psychomimetic side effects. The rational design and subsequent electrophysiological testing of 27 novel dibenzosuberane derivatives demonstrated varying inhibitory activity toward the major NMDA receptor subtypes. Based on a combination of *in silico* predictions, *in vitro* assays, and *in vivo* pharmacokinetics, we selected compounds **3l** and **6f** for further *in vivo* analysis based on their potency, bioavailability, and low cytotoxicity. Compound **6f** showed high inhibition of both GluN1/GluN2A and GluN1/GluN2B receptor subtypes, while compound **3l** showed moderate inhibition but exhibited significant neuroprotective effect in a model of NMDA-induced hippocampal neurodegeneration. Behavioral tests confirmed that both compounds **3l** and **6f** did not induce hyperlocomotion and prepulse inhibition deficits, suggesting a lower risk of psychomimetic side effects than other NMDA receptor antagonists, such as MK-801. Thus, compound **3l** became a prime candidate due to its neuroprotective effects and lower toxicity, suggesting its potential for therapeutic use with minimal adverse effects.

## 4. Experimental Section

### 5.1. Chemistry

All chemical solvents and reagents were used in the highest available purity without further purification. They were purchased from Sigma-Aldrich (Prague, Czech Republic) or FluoroChem (Dublin, Ireland). The reactions were monitored by thin layer chromatography (TLC) on silica gel plates (60 F254, Merck, Prague, Czech Republic), and the spots were visualized by ultraviolet light (254 nm). Purification of crude products was carried out using columns of silica gel (silica gel 100, 0.063-0.200 mm, 70-230 mesh ASTM, Fluka, Prague, Czech Republic) and PuriFlash GEN5 column, 5.250 (Interchim, Montluçon, France; silica gel 100, 60 Å, 230–400 mesh ASTM, Sigma-Aldrich, Prague, Czech Republic). NMR spectra were recorded in deuterated chloroform (CDCl_3_), deuterated methanol (CD_3_OD), and deuterated dimethyl sulfoxide (DMSO-*d*_6_) on Bruker Avance NEO 500 MHz spectrometer (499.87 MHz for ^1^H NMR, and 125.71 MHz for ^13^C NMR). Chemical shifts (*δ*) are reported in parts per million (ppm), and spin multiplicities are given as broad singlet (bs), doublet (d), doublet of doublet (dd), triplet (t), quartet (q), pentet (p), or multiplet (m). Coupling constants (*J*) are reported in Hz. The synthesized compounds were analyzed by LC-MS system consisting of UHLPC Dionex Ultimate 3000 RS couplet with Q Exactive Plus mass spectrometer to obtain high-resolution mass spectra (Thermo Fisher Scientific, Bremen, Germany). Melting points were measured using an automated melting point recorder M-565 (Büchi, Switzerland) and are uncalibrated. The final compounds were analyzed by LC-MS consisting of UHPLC Dionex Ultimate 3000 RS coupled with Q Exactive Plus orbitrap mass spectrometer (Thermo Fisher Scientific, Bremen, Germany) to obtain high-resolution mass spectra. Gradient LC analysis confirmed > 95% purity for all the final compounds.

### 5.2. General Procedure for the Preparation of N-substituted-10,11-dihydro-5H-dibenzo[a,d][7]annulene-5-amines (3a-o)

Primary or secondary amine (2a-o)(5.25 mM, 3 eq)(Scheme 1) was placed in a 50 mL flask and dissolved in 10 mL of dry MeCN. The reaction mixture was stirred under an argon atmosphere at room temperature. Commercially available 5-chloro-10,11-dihydro-5*H*-dibenzo[*a,d*][7]annulene (1)(1.75 mM, 1 eq) was dissolved in 5 mL of dry MeCN and added dropwise to the reaction. The reaction was stirred for 3 h under an argon atmosphere at 40 °C. After cooling to room temperature, the solvent was evaporated under reduced pressure, and the crude material was purified using flash chromatography (eluent CH_2_Cl_2_/MeOH with 1% of 25% aq. NH_3_, gradient 98:2 → 94:6) to obtain the final compounds as free bases. These were dissolved in 5 mL of MeOH and 1 mL of HCl (35% aq. sol.) and were added dropwise at 0°C. The reaction was maintained for 1 h at room temperature. The solvent was evaporated under reduced pressure, and the residual water was removed by azeotropic distillation with absolute EtOH. The solid product was washed with acetone for the final hydrochloride salts (3a-o) as a yellowish solid.

#### 5.2.1. N-methyl-10,11-dihydro-5H-dibenzo[a,d][7]annulen-5-amine hydrochloride (3a)(K1934)

Yield 17 %; white solid. M.p.: 238.9 – 240.6 °C. ^1^H NMR (500 MHz, Methanol-*d*_4_) δ 7.54 (dd, *J* = 7.5, 1.4 Hz, 2H), 7.42 (td, *J* = 7.5, 1.4 Hz, 2H), 7.37 – 7.31 (m, 4H), 5.43 (s, 1H), 3.52 – 3.41 (m, 2H), 3.14 – 3.02 (m, 2H), 2.63 (s, 3H). ^13^C NMR (126 MHz, MeOD) δ 140.31, 131.64, 131.07, 130.13, 126.74, 32.15, 31.21. HRMS (ESI^+^ ): [M + H]^+^: calculated for C_16_H_18_N^+^ (m/z): 224.14337; found: 224.14308. LC-MS purity 99.9%.

#### 5.2.2. N-ethyl-10,11-dihydro-5H-dibenzo[a,d][7]annulen-5-amine hydrochloride (3b)(K1935)

Yield 51 %; white solid. M.p.: 226.0 – 227.6 °C. ^1^H NMR (500 MHz, Methanol-*d*_4_) δ 7.54 (dd, *J* = 7.5, 1.4 Hz, 2H), 7.41 (td, *J* = 7.5, 1.4 Hz, 2H), 7.37 – 7.30 (m, 4H), 5.48 (s, 1H), 3.54 – 3.41 (m, 2H), 3.14 – 2.99 (m, 4H), 1.34 (t, *J* = 7.3 Hz, 3H). ^13^C NMR (126 MHz, MeOD) δ 131.02, 130.05, 126.69, 42.00, 32.10, 10.02. HRMS (ESI^+^ ): [M + H]^+^: calculated for C_17_H_20_N^+^ (m/z): 238.15902; found: 238.15881. LC-MS purity 99.9%.

#### 5.2.3. N-isopropyl-10,11-dihydro-5H-dibenzo[a,d][7]annulen-5-amine hydrochloride (3c)(K1936)

Yield 52 %; white solid. M.p.: 202.2 – 203.9 °C. ^1^H NMR (500 MHz, Methanol-*d*_4_) δ 7.56 (dd, *J* = 7.5, 1.4 Hz, 2H), 7.42 (td, *J* = 7.5, 1.4 Hz, 2H), 7.36 – 7.30 (m, 4H), 5.56 (s, 1H), 3.51 – 3.40 (m, 2H), 3.34 – 3.31 (m, 1H), 3.14 – 3.02 (m, 2H), 1.40 (d, *J* = 6.6 Hz, 6H). ^13^C NMR (126 MHz, MeOD) δ 140.32, 132.23, 131.07, 130.04, 126.70, 50.09, 32.05, 18.13. HRMS (ESI^+^ ): [M + H]^+^: calculated for C_18_H_22_N^+^ (m/z): 252.17467; found: 252.17441. LC-MS purity 99.9%.

#### 5.2.4. N-cyclopropyl-10,11-dihydro-5H-dibenzo[a,d][7]annulen-5-amine hydrochloride (3d)(K1937)

Yield 62 %; white solid. M.p.: 204.6 – 205.6 °C. ^1^H NMR (500 MHz, Methanol-*d*_4_) δ 7.59 (dd, *J* = 7.5, 1.4 Hz, 2H), 7.42 (td, *J* = 7.5, 1.4 Hz, 2H), 7.37 – 7.30 (m, 4H), 5.61 (s, 1H), 3.55 – 3.42 (m, 2H), 3.14 – 3.01 (m, 2H), 2.64 – 2.56 (m, 1H), 0.96 – 0.91 (m, 2H), 0.88 – 0.82 (m, 2H). ^13^C NMR (126 MHz, MeOD) δ 129.14, 128.31, 124.90, 30.43, 28.01, 1.42. HRMS (ESI^+^ ): [M + H]^+^: calculated for C_18_H_20_N^+^ (m/z): 250.15902; found: 250.15881. LC-MS purity 99.9%.

#### 5.2.5. N-butyl-10,11-dihydro-5H-dibenzo[a,d][7]annulen-5-amine hydrochloride (3e)(K1938)

Yield 57 %; white solid. M.p.: 220.4 – 223.0 °C. ^1^H NMR (500 MHz, Methanol-*d*_4_) δ 7.54 (dd, *J* = 7.5, 1.4 Hz, 2H), 7.42 (td, *J* = 7.5, 1.4 Hz, 2H), 7.38 – 7.30 (m, 4H), 5.49 (s, 1H), 3.52 – 3.41 (m, 2H), 3.15 – 3.02 (m, 2H), 2.97 – 2.88 (m, 2H), 1.78 – 1.67 (m, 2H), 1.42 – 1.31 (m, 2H), 0.94 (t, *J* = 7.4 Hz, 3H). ^13^C NMR (126 MHz, MeOD) δ 131.02, 130.08, 126.70, 46.57, 32.10, 27.49, 19.57, 12.42. HRMS (ESI^+^ ): [M + H]^+^: calculated for C_19_H_24_N^+^ (m/z): 266.19032; found: 266.18967. LC-MS purity 99.9%.

#### 5.2.6. N-isobutyl-10,11-dihydro-5H-dibenzo[a,d][7]annulen-5-amine hydrochloride (3f)(K1939)

Yield 24 %; white solid. M.p.: 195.6 – 196.2 °C. ^1^H NMR (500 MHz, Methanol-*d*_4_) δ 7.54 (dd, *J* = 7.5, 1.4 Hz, 2H), 7.42 (td, *J* = 7.5, 1.4 Hz, 2H), 7.37 – 7.30 (m, 4H), 5.52 (s, 1H), 3.51 – 3.40 (m, 2H), 3.18 – 3.04 (m, 2H), 2.81 – 2.72 (m, 2H), 2.17 – 2.05 (m, 1H), 1.00 (d, *J* = 6.7 Hz, 6H). ^13^C NMR (126 MHz, MeOD) δ 131.02, 130.11, 126.69, 53.92, 32.10, 25.40, 19.20. HRMS (ESI^+^ ): [M + H]^+^: calculated for C_19_H_24_N^+^ (m/z): 266.19032; found: 266.18994. LC-MS purity 99.9%.

#### 5.2.7. N-cyclohexyl-10,11-dihydro-5H-dibenzo[a,d][7]annulen-5-amine hydrochloride (3g)(K1940)

Yield 19 %; white solid. M.p.: 218.4 – 220.3 °C. ^1^H NMR (500 MHz, Methanol-*d*_4_) δ 7.55 (dd, *J* = 7.5, 1.4 Hz, 2H), 7.41 (td, *J* = 7.5, 1.4 Hz, 2H), 7.37 – 7.29 (m, 4H), 5.61 (s, 1H), 3.51 – 3.40 (m, 2H), 3.15 – 3.02 (m, 2H), 3.00 – 2.90 (m, 1H), 2.26 – 2.17 (m, 2H), 1.90 – 1.81 (m, 2H), 1.72 – 1.64 (m, 1H), 1.52 – 1.37 (m, 2H), 1.31 – 1.14 (m, 3H). ^13^C NMR (126 MHz, MeOD) δ 140.35, 132.33, 131.04, 130.00, 126.68, 56.89, 32.07, 29.14, 24.55, 24.25. HRMS (ESI^+^ ): [M + H]^+^: calculated for C_21_H_26_N^+^ (m/z): 292.20597; found: 292.20572. LC-MS purity 96%.

#### 5.2.8. 1-(10,11-dihydro-5H-dibenzo[a,d][7]annulen-5-yl)piperidine hydrochloride (3h)(K1941)

Yield 38 %; white solid. M.p.: 200.1 – 201.3 °C. ^1^H NMR (500 MHz, Methanol-*d*_4_) δ 7.52 (dd, *J* = 7.5, 1.4 Hz, 2H), 7.43 (td, *J* = 7.5, 1.4 Hz, 2H), 7.39 – 7.27 (m, 4H), 5.36 (s, 1H), 3.56 – 3.44 (m, 2H), 3.41 – 3.34 (m, 2H), 3.14 – 3.01 (m, 4H), 2.01 – 1.89 (m, 2H), 1.87 – 1.74 (m, 3H), 1.64 – 1.50 (m, 1H). ^13^C NMR (126 MHz, MeOD) δ 139.95, 132.86, 131.24, 130.73, 130.48, 126.51, 77.49, 52.63, 31.85, 22.52, 21.16. HRMS (ESI^+^ ): [M + H]^+^: calculated for C_20_H_24_N^+^ (m/z): 278.19032; found: 278.19012. LC-MS purity 99.9%.

#### 5.2.9. 4-(10,11-dihydro-5H-dibenzo[a,d][7]annulen-5-yl)morpholine hydrochloride (3i)(K1943)

Yield 60 %; white solid. M.p.: 185.6 – 187.2 °C. ^1^H NMR (500 MHz, Methanol-*d*_4_) δ 7.53 (dd, *J* = 7.5, 1.4 Hz, 2H), 7.44 (dt, *J* = 7.5, 1.4 Hz, 2H), 7.41 – 7.28 (m, 4H), 5.44 (s, 1H), 4.10 – 3.95 (m, 2H), 3.91 – 3.77 (m, 2H), 3.62 – 3.48 (m, 2H), 3.32 – 3.19 (m, 4H), 3.16 – 2.99 (m, 2H). ^13^C NMR (126 MHz, MeOD) δ 140.23, 133.05, 131.31, 130.70, 129.91, 126.58, 78.60, 63.45, 51.73, 31.94. HRMS (ESI^+^ ): [M + H]^+^: calculated for C_19_H_22_NO^+^ (m/z): 280.16959; found: 280.16928. LC-MS purity 95%.

#### 5.2.10. N-(2-methoxyethyl)-10,11-dihydro-5H-dibenzo[a,d][7]annulen-5-amine hydrochloride (3j)(K1945)

Yield 56 %; white solid. M.p.: 205.8 – 207.2 °C. ^1^H NMR (500 MHz, Methanol-*d*_4_) δ 7.53 (dd, *J* = 7.5, 1.4 Hz, 2H), 7.41 (td, *J* = 7.5, 1.4 Hz, 2H), 7.36 – 7.30 (m, 4H), 5.57 (s, 1H), 3.63 (t, *J* = 5.1 Hz, 2H), 3.52 – 3.41 (m, 2H), 3.38 (s, 3H), 3.14 (t, *J* = 5.1 Hz, 2H), 3.12 – 3.04 (m, 2H). ^13^C NMR (126 MHz, MeOD) δ 131.02, 130.05, 126.68, 66.61, 57.81, 45.92, 32.07. HRMS (ESI^+^ ): [M + H]^+^: calculated for C_18_H_22_NO^+^ (m/z): 268.16959; found: 268.16934. LC-MS purity 99.9%.

#### 5.2.11. 1-(10,11-dihydro-5H-dibenzo[a,d][7]annulen-5-yl)pyrrolidine hydrochloride (3k)(K1948)

Yield 38 %; white solid. M.p.: 222.3 – 223.9 °C. ^1^H NMR (500 MHz, Methanol-*d*_4_) δ 7.54 (dd, *J* = 7.5, 1.4 Hz, 2H), 7.43 (td, *J* = 7.5, 1.4 Hz, 2H), 7.38 – 7.28 (m, 4H), 5.41 (s, 1H), 3.64 – 3.52 (m, 2H), 3.32 – 3.28 (m, 4H), 3.15 – 3.03 (m, 2H), 2.28 – 2.14 (m, 2H), 2.12 – 1.97 (m, 2H). ^13^C NMR (126 MHz, MeOD) δ 139.82, 132.12, 132.05, 131.24, 130.37, 126.63, 75.86, 54.00, 31.98, 22.18. HRMS (ESI^+^ ): [M + H]^+^: calculated for C_19_H_22_N^+^ (m/z): 264.17467; found: 264.17435. LC-MS purity 99.9%.

#### 5.2.12. 2-((10,11-dihydro-5H-dibenzo[a,d][7]annulen-5-yl)amino)ethanol hydrochloride (3l)(K1949)

Yield 53 %; white solid. M.p.: 183.1 – 184.8 °C. ^1^H NMR (500 MHz, Methanol-*d*_4_) δ 7.54 (dd, *J* = 7.5, 1.4 Hz, 2H), 7.41 (td, *J* = 7.5, 1.4 Hz, 2H), 7.36 – 7.30 (m, 4H), 5.60 (s, 1H), 3.84 – 3.78 (m, 2H), 3.54 – 3.43 (m, 2H), 3.15 – 3.02 (m, 4H). ^13^C NMR (126 MHz, MeOD) δ 131.02, 130.02, 126.67, 56.35, 32.11. HRMS (ESI^+^ ): [M + H]^+^: calculated for C_17_H_20_NO^+^ (m/z): 254.15394; found: 254.15373. LC-MS purity 99.9%.

#### 5.2.13. 1-(10,11-dihydro-5H-dibenzo[a,d][7]annulen-5-yl)-4-methylpiperazine dihydrochloride (3m)(K1942)

Yield 33 %; white solid. M.p.: decomposition at 228.7 °C. ^1^H NMR (500 MHz, Methanol-*d*_4_) δ 7.26 – 7.19 (m, 4H), 7.17 (dd, *J* = 7.5, 1.4 Hz, 2H), 7.12 (td, *J* = 7.5, 1.4 Hz, 2H), 4.16 (s, 1H), 4.02 – 3.92 (m, 2H), 3.26 – 3.08 (m, 4H), 2.89 – 2.77 (m, 2H), 2.67 – 2.47 (m, 4H). ^13^C NMR (126 MHz, MeOD) δ 139.64, 137.79, 130.66, 130.61, 128.00, 125.50, 77.69, 53.71, 48.68, 42.06, 31.42. HRMS (ESI^+^ ): [M + H]^+^: calculated for C_20_H_25_N_2_ (m/z): 293.20122; found: 293.20096. LC-MS purity 99.9%.

#### 5.2.14. N,N-diethyl-10,11-dihydro-5H-dibenzo[a,d][7]annulen-5-amine hydrochloride (3n)(K1946)

Yield 55 %; white solid. M.p.: 166.2 – 167.4 °C. ^1^H NMR (500 MHz, DMSO-*d*_6_) δ 9.87 (s, 1H), 7.55 (dd, *J* = 7.5, 1.4 Hz, 2H), 7.39 (td, *J* = 7.5, 1.4 Hz, 2H), 7.30 (dd, *J* = 7.5, 1.4 Hz, 2H), 7.27 (td, *J* = 7.5, 1.4 Hz, 2H), 5.51 (d, *J* = 8.8 Hz, 1H), 3.75 – 3.63 (m, 2H), 3.16 – 3.06 (m, 2H), 3.02 – 2.89 (m, 4H), 1.25 (t, *J* = 7.2 Hz, 6H). ^13^C NMR (126 MHz, DMSO) δ 140.32, 132.94, 132.20, 131.77, 130.63, 126.78, 71.95, 44.32, 31.83, 8.10. HRMS (ESI^+^ ): [M + H]^+^: calculated for C_19_H_24_N^+^ (m/z): 226.19032; found: 266.19046. LC-MS purity 99.9%.

#### 5.2.15. 1-(10,11-dihydro-5H-dibenzo[a,d][7]annulen-5-yl)-4-ethylpiperazine dihydrochloride (3o)(K1947)

Yield 36 %; white solid. M.p.: decomposition at 239.3 °C. ^1^H NMR (500 MHz, Methanol-*d*_4_) δ 7.26 – 7.19 (m, 4H), 7.16 (dd, *J* = 7.5, 1.4 Hz, 2H), 7.11 (td, *J* = 7.5, 1.4 Hz, 2H), 4.14 (s, 1H), 4.04 – 3.90 (m, 2H), 3.24 – 2.96 (m, 6H), 2.89 – 2.75 (m, 2H), 2.67 – 2.42 (m, 4H), 1.28 (t, *J* = 7.3 Hz, 3H). ^13^C NMR (126 MHz, MeOD) δ 139.63, 137.94, 130.64, 130.60, 127.96, 125.48, 77.84, 51.64, 51.50, 48.81, 31.43, 8.47. HRMS (ESI^+^ ): [M + H]^+^: calculated for C_21_H_27_N_2_ (m/z): 307.21687; found: 307.21658. LC-MS purity 99.9%.

### 5.3. General Procedure for the Preparation of N-substituted-5H-dibenzo[a,d][7]annulen-5-amines (6a-l)

Commercially available 5*H*-dibenzo[*a,d*][7]annulen-5-ol (4)(4.8 mM, 1 eq) was placed in 25 mL flask and dissolved in dry DCM, flask was filled with argon and cooled to 0 °C. Solution of SOCl_2_ (20.67 mM, 4.3 eq), 1 mL of dry DCM and 1 drop of pyridine was added dropwise. Reaction mixture was stirred at room temperature for 12 h under argon atmosphere. After the reaction proceeded, solvent was evaporated under reduced pressure and crude material was crystallized from petroleum ether to obtain the final compound (5) as yellow solid. 5 was used directly into the next reaction.

Compounds 6a-l were prepared according to the procedure used in 4.2.

#### 5.3.1. N-methyl-5H-dibenzo[a,d][7]annulen-5-amine hydrochloride (6a)(K2051)

Yield 43 %; white solid. M.p.: 208.7 – 209.7 °C. ^1^H NMR (500 MHz, Methanol-*d*_4_) δ 7.75 – 7.69 (m, 2H), 7.66 – 7.60 (m, 2H), 7.60 – 7.54 (m, 4H), 7.25 (s, 2H), 5.66 (s, 1H), 2.36 (s, 3H). ^13^C NMR (126 MHz, MeOD) δ 133.62, 130.91, 130.44, 130.35, 130.26, 129.54, 129.46, 68.17, 30.74. HRMS (ESI^+^): [M + H]^+^: calculated for (m/z): 222.12772; found: 222.12752. LC-MS purity 98%.

#### 5.3.2. N-ethyl-5H-dibenzo[a,d][7]annulen-5-amine hydrochloride (6b)(K2052)

Yield 44 %; white solid. M.p.: 189.0 – 190.9 °C. ^1^H NMR (500 MHz, Methanol-*d*_4_) δ 7.76 – 7.71 (m, 2H), 7.64 – 7.60 (m, 2H), 7.59 – 7.53 (m, 4H), 7.25 (s, 2H), 5.76 (s, 1H), 2.70 (q, *J* = 7.3 Hz, 2H), 1.21 (t, *J* = 7.3 Hz, 3H). ^13^C NMR (126 MHz, MeOD) δ 133.83, 130.98, 130.44, 130.32, 130.20, 129.47, 129.43, 66.44, 41.31, 9.54. HRMS (ESI^+^): [M + H]^+^: calculated for C_17_H_18_N^+^ (m/z): 236.14337; found: 236.14259. LC-MS purity 99%.

#### 5.3.3. N-isopropyl-5H-dibenzo[a,d][7]annulen-5-amine hydrochloride (6c)(K2053)

Yield 41 %; white solid. Melting point (m.p.): 189.5 – 190.7 °C. ^1^H NMR (500 MHz, Methanol-*d*_4_) δ 7.80 – 7.74 (m, 2H), 7.64 – 7.59 (m, 2H), 7.58 – 7.51 (m, 4H), 7.27 (s, 2H), 5.87 (s, 1H), 2.95 (hept, *J* = 6.6 Hz, 1H), 1.21 (d, *J* = 6.6 Hz, 6H). ^13^C NMR (126 MHz, MeOD) δ 133.97, 131.29, 130.58, 130.10, 129.98, 129.44, 129.36, 64.73, 50.03, 17.97. HRMS (ESI^+^): [M + H]^+^: calculated for C_18_H_20_N^+^ (m/z): 250.15902; found: 250.15836 94%. LC-MS purity 95%.

#### 4.3.4. N-cyclopropyl-5H-dibenzo[a,d][7]annulen-5-amine hydrochloride (6d)(K2054)

Yield 68 %; white solid. M.p.: 195.5 – 196.0 °C. ^1^H NMR (500 MHz, Methanol-*d*_4_) δ 7.79 – 7.74 (m, 2H), 7.64 – 7.59 (m, 2H), 7.59 – 7.53 (m, 4H), 7.24 (s, 2H), 5.84 (s, 1H), 2.17 – 2.08 (m, 1H), 0.83 – 0.76 (m, 2H), 0.75 – 0.69 (m, 2H). ^13^C NMR (126 MHz, MeOD) δ 132.36, 129.43, 128.97, 128.87, 128.54, 127.90, 127.81, 66.85, 27.47, 1.42. HRMS (ESI^+^): [M + H]^+^: calculated for C_18_H_18_N^+^ (m/z): 248.14337; found: 248.14290. LC-MS purity 98%.

#### 5.3.5. N-butyl-5H-dibenzo[a,d][7]annulen-5-amine hydrochloride (6e)(K2055)

Yield 57 %; white solid. M.p.: 202.7 – 203.2 °C. ^1^H NMR (500 MHz, Methanol-*d*_4_) δ 7.76 – 7.71 (m, 2H), 7.65 – 7.60 (m, 2H), 7.59 – 7.53 (m, 4H), 7.26 (s, 2H), 5.77 (s, 1H), 2.65 – 2.56 (m, 2H), 1.64 – 1.53 (m, 2H), 1.31 – 1.18 (m, 2H), 0.87 (t, *J* = 7.4 Hz, 3H). ^13^C NMR (126 MHz, MeOD) δ 133.81, 130.96, 130.45, 130.35, 130.20, 129.49, 129.44, 66.73, 45.85, 26.99, 19.46, 12.33. HRMS (ESI^+^): [M + H]^+^: calculated for C_19_H_22_N^+^ (m/z): 264.17467; found: 264.17410. LC-MS purity 99.9%.

#### 5.3.6. N-isobutyl-5H-dibenzo[a,d][7]annulen-5-amine hydrochloride (6f)(K2056)

Yield 42 %; white solid. M.p.: 183.6 – 184.4 °C. ^1^H NMR (500 MHz, Methanol-*d*_4_) δ 7.77 – 7.72 (m, 2H), 7.65 – 7.61 (m, 2H), 7.60 – 7.54 (m, 4H), 7.28 (s, 2H), 5.79 (s, 1H), 2.45 (d, *J* = 7.1 Hz, 2H), 2.01 – 1.90 (m, 1H), 0.88 (d, *J* = 6.7 Hz, 6H). ^13^C NMR (126 MHz, MeOD) δ 133.71, 130.88, 130.52, 130.38, 130.17, 129.50, 66.78, 53.04, 24.81, 18.98. HRMS (ESI^+^): [M + H]^+^: calculated for C_19_H_22_N^+^ (m/z): 264.17467; found: 264.17392. LC-MS purity 99%.

#### 5.3.7. N-cyclohexyl-5H-dibenzo[a,d][7]annulen-5-amine hydrochloride (6g)(K2057)

Yield 74 %; white solid. M.p.: 199.8 – 200.2 °C. ^1^H NMR (500 MHz, Methanol-*d*_4_) δ 7.78 – 7.73 (m, 2H), 7.65 – 7.60 (m, 2H), 7.59 – 7.52 (m, 4H), 7.27 (s, 2H), 5.91 (s, 1H), 2.63 – 2.52 (m, 1H), 1.98 – 1.90 (m, 2H), 1.83 – 1.73 (m, 2H), 1.65 – 1.57 (m, 1H), 1.36 – 1.24 (m, 2H), 1.18 – 1.03 (m, 3H). ^13^C NMR (126 MHz, MeOD) δ 133.98, 131.24, 130.58, 130.11, 130.00, 129.43, 129.38, 64.60, 56.85, 28.94, 24.41, 24.24. HRMS (ESI^+^): [M + H]^+^: calculated for C_21_H_24_N^+^ (m/z): 290.19032; found: 290.18954. LC-MS purity 99.9%.

#### 5.3.8. 1-(5H-dibenzo[a,d][7]annulen-5-yl)piperidine hydrochloride (6h)(K2058)

Yield 44 %; white solid. M.p.: 169.8 – 171.0 °C. ^1^H NMR (500 MHz, DMSO-*d*_6_) δ 8.52 – 8.40 (m, 1H), 7.86 – 7.81 (m, 2H), 7.69 – 7.64 (m, 2H), 7.60 – 7.53 (m, 4H), 6.26 – 6.15 (m, 1H), 2.78 – 2.61 (m, 4H), 1.86 – 1.73 (m, 2H), 1.70 – 1.58 (m, 2H), 1.56 – 1.36 (m, 2H). ^13^C NMR (126 MHz, DMSO) δ 134.55, 132.18, 131.15, 130.70, 130.13, 129.82, 51.28, 21.71, 20.74. HRMS (ESI^+^): [M + H]^+^: calculated for C_20_H_22_N^+^ (m/z): 276.17467; found: 276.17404. LC-MS purity 99%.

#### 5.3.9. 4-(5H-dibenzo[a,d][7]annulen-5-yl)morpholine hydrochloride (6i)(K2059)

Yield 52 %; white solid. M.p.: 171.1 – 172.7 °C. ^1^H NMR (500 MHz, Methanol-*d*_4_) δ 7.79 – 7.73 (m, 2H), 7.69 – 7.64 (m, 2H), 7.63 – 7.57 (m, 4H), 7.28 (s, 2H), 5.92 (s, 1H), 3.96 – 3.85 (m, 3H), 3.74 – 3.65 (m, 2H), 3.28 – 3.22 (m, 1H), 3.18 – 3.09 (m, 2H), 2.78 – 2.69 (m, 2H). ^13^C NMR (126 MHz, MeOD) δ 134.14, 131.67, 130.48, 130.45, 130.05, 129.50, 128.77, 127.90, 76.97, 63.49, 62.42, 51.12, 43.26. HRMS (ESI^+^): [M + H]^+^: calculated for C_19_H_20_NO^+^ (m/z): 278.15394; found: 278.15356. LC-MS purity 97%.

#### 5.3.10. N-(2-methoxyethyl)-5H-dibenzo[a,d][7]annulen-5-amine hydrochloride (6j)(K2061)

Yield 52 %; white solid. M.p.: 189.1 – 190.8 °C. ^1^H NMR (500 MHz, Methanol-*d*_4_) δ 7.74 – 7.71 (m, 2H), 7.64 – 7.61 (m, 2H), 7.59 – 7.53 (m, 4H), 7.27 (s, 2H), 5.81 (s, 1H), 3.52 – 3.46 (m, 2H), 3.32 (s, 3H), 2.90 – 2.81 (m, 2H). ^13^C NMR (126 MHz, MeOD) δ 133.67, 130.97, 130.48, 130.33, 130.18, 129.48, 66.81, 66.18, 57.75, 45.17. HRMS (ESI^+^): [M + H]^+^: calculated for C_18_H_20_NO^+^ (m/z): 266.15394; found: 266.15411. LC-MS purity 99%.

#### 5.3.11. 1-(5H-dibenzo[a,d][7]annulen-5-yl)pyrrolidine hydrochloride (6k)(K2062)

Yield 58 %; white solid. M.p.: 170.1 – 171.3 °C. ^1^H NMR (500 MHz, Methanol-*d*_4_) δ 7.76 – 7.73 (m, 2H), 7.66 – 7.62 (m, 2H), 7.59 – 7.56 (m, 4H), 7.30 (s, 2H), 5.74 (s, 1H), 3.15 – 3.05 (m, 2H), 2.88 – 2.80 (m, 2H), 2.19 – 2.08 (m, 2H), 1.99 – 1.88 (m, 2H). ^13^C NMR (126 MHz, MeOD) δ 133.63, 131.06, 130.51, 130.50, 130.19, 129.67, 129.49, 74.64, 53.89, 22.66. HRMS (ESI^+^): [M + H]^+^: calculated for C_19_H_20_N^+^ (m/z): 262.15902; found: 262.15942. LC-MS purity 99%.

#### 5.3.12. 2-((5H-dibenzo[a,d][7]annulen-5-yl)amino)ethanol hydrochloride (6l)(K2063)

Yield 69 %; white solid. M.p.: 181.5 – 182.3 °C. ^1^H NMR (500 MHz, Methanol-*d*_4_) δ 7.75 – 7.70 (m, 2H), 7.66 – 7.61 (m, 2H), 7.60 – 7.53 (m, 4H), 7.28 (s, 2H), 5.85 (s, 1H), 3.69 – 3.63 (m, 2H), 2.80 – 2.73 (m, 2H). ^13^C NMR (126 MHz, MeOD) δ 133.62, 131.03, 130.49, 130.29, 130.20, 129.51, 129.47, 66.60, 55.92. HRMS (ESI^+^): [M + H]^+^: calculated for C_17_H_18_NO^+^ (m/z): 252.13829; found: 252.13858. LC-MS purity 99.9%.

### 5.4. Molecular biology and preparation of transfected HEK293 cells

We used cDNA vectors for expression in mammalian cells carrying genes for the human versions of the GluN1-1a (hGluN1), GluN2A (hGluN2A), and GluN2B (hGluN2B) subunits [58]. HEK293 cells were cultured in 24-well plates in Opti-MEM I medium containing 5% fetal bovine serum (FBS; all from Thermo Fischer Scientific, Prague, Czech Republic). Confluent HEK293 cells were transfected using a mixture of 50 μl of Opti-MEM I containing a total of 0.9 μg of cDNA constructs encoding hGluN1/hGluN2 subunits and GFP (for identification of successfully transfected cells; pQBI 25, Takara) in a 1:1:1 ratio, plus 0.9 μl of PolyMag Transfection Reagent (OZ BIOSCIENCES, Marseille, France). After a 20 min incubation on a magnetic plate, HEK293 cells were trypsinized and cultured on 3 cm diameter poly-L-lysine-coated coverslips in Opti-MEM I medium containing 1% FBS, 20 mM MgCl_2,_ and 3 mM kynurenic acid (to prevent cell death caused by excitotoxic NMDA receptor activation) [47].

### 5.5. Electrophysiology

Patch-clamp measurements in the whole-cell configuration were performed 16 - 48 hours after the end of transfection using an Axopatch 200B amplifier (Molecular Devices, California, USA); cell current responses were low-pass filtered at 2 kHz using a 4-pole Bessel low-pass filter, digitized at 5 kHz, and recorded using Clampex 10 software (Axon Instruments, Molecular Devices). The series resistance and capacitance of all measured cells were monitored and compensated to 80% [47]. Glass micropipettes with a tip resistance of 3-6 MΩ were prepared using a micropipette puller model P-1000 (Sutter Instrument Co., California, USA) and were filled with an intracellular solution containing (in mM): 125 gluconic acid, 15 CsCl, 10 BAPTA 10 HEPES, 3 MgCl_2_, 0.5 CaCl_2_ and 2 ATP-Mg salts (pH 7.2 with CsOH). The extracellular recording solution (ECS) contained (in mM) 160 NaCl, 2.5 KCl, 10 HEPES, 10 D-glucose, 0.01 EDTA, and 1 CaCl_2_ (pH 7.3 with NaOH). All chemicals were purchased from Sigma-Aldrich (Prague, Czech Republic). ECS exchange was performed using a rapid application system with a solution exchange time constant per cell of approximately 20 ms. All ECS contained a saturating concentration of the co-agonist – glycine (100 µM), and NMDA receptor responses were induced by applying a saturating concentration of the agonist – L-glutamate (1 mM). The test compounds were dissolved in dimethyl sulfoxide (DMSO) at a concentration of 30 mM; all ECS used in a given experiment contained the same concentration of DMSO (≤1%). Concentration-response inhibition curves were obtained using the

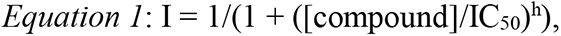

where IC_50_ is the concentration of the compound that caused 50% inhibition of the L-glutamate-induced current, [compound] is the concentration of the tested compound, and h is the Hill coefficient [53].

The time course of inhibition by 6f or 3l was analyzed using a single or double exponential function, and the weighted time constants (τw) were obtained using the

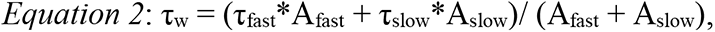

where τ_fast_ and A_fast_ are the time constant and area of the faster component, and τ_slow_ and A_slow_ are the time constant and area of the slower component.

### 5.6. Cell viability assay

Standard MTT assay (Merck) was used according to the manufactureŕs protocol on the CHO-K1 cell line (Chinese hamster ovary) to compare the cytotoxic effect of the studied compounds [59]. The CHO-K1 cells were cultured according to ECACC-recommended conditions and seeded in a density of 8,000 cells per well. Briefly, the tested compounds were dissolved in DMSO and subsequently diluted in a growth medium (F-12) supplemented with 10% FBS and 1% penicillin/streptomycin so that the final concentration of DMSO did not exceed 0.5% (v/v). The CHO-K1 cells were exposed to the tested compounds for 24 h. The medium was then replaced with medium containing 10 μM MTT, and the cells were allowed to produce formazan for another approximately 3 h under surveillance. Thereafter, the medium with MTT was removed, and the formazan crystals were dissolved in DMSO (100 µL). Cell viability was assessed spectrophotometrically by the amount of formazan produced. Absorbance was measured at a wavelength of 570 nm with a reference wavelength of 650 nm on Synergy HT (BioTek, Winooski, USA). The IC_50_ values were then calculated from the control-subtracted triplicates using non-linear regression (four parameters) in GraphPad Prism 5 software (GraphPad Software, San Diego, USA). The final IC_50_ values were obtained as a mean ± standard error of the mean (SEM) from three independent measurements.

### 5.7 In Vitro Anti-Cholinesterase Assay

The inhibitory activity of novel compounds against human recombinant AChE (hAChE, E.C. 3.1.1.7, purchased from Merck) and human plasmatic butyrylcholinesterase (hBChE, E.C. 3.1.1.8, purchased from Merck) were determined using the modified Ellman’s method, according to the previously published protocol with the highest concentration tested 100 µM [60]. The results are expressed as IC_50_ values (the concentration of the compound that is required to reduce cholinesterase activity by 50%). For the calculation of the degree of inhibition (I), the following Equation 3 was used:

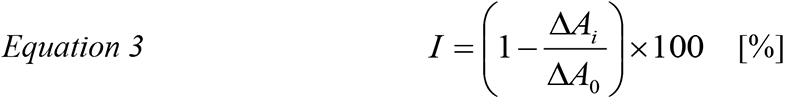

where ΔAi indicates the change in absorbance observed for adequate cholinesterase exposed to its corresponding inhibitor, and ΔA0 indicates the change in absorbance when a phosphate buffer solution (PBS) was added instead of an inhibitor. Microsoft Excel 10 software (Microsoft Corporation, Redmont, USA) and Prism v. 5.02 (GraphPad Software, San Diego, USA) were used for statistical evaluation of the obtained data.

### 5.8. In vivo *pharmacokinetic study*

#### 5.8.1. Animals

Adult male Wistar rats (200-300 g) purchased from the Velaz breeding colony (Czech Republic) were used for the pharmacokinetic assessment of four selected compounds (3g, 3l, 6f, and 6g). The experiments were conducted in accordance with the guidelines of the European Union directive 2010/63/EU and Act No. 246/1992 Coll. The experimental animals were handled under the supervision of the Ethics Committee of the Faculty of Military Health Sciences, Czech Republic (approvals reference No. 15/21 and 19/21).

The rats were injected intraperitoneally (*i.p.*) with the tested compounds (5 mg/kg in 5% DMSO/5% kolliphor/saline (v/v) mixture). Blood samples were collected under deep terminal anesthesia directly by cardiac puncture into heparinized 1.5 mL tubes at 15 and 60 min (three animals per time interval). Four animals were used for zero time or blank control. The animals were perfused transcardially with a saline solution (0.9% NaCl) for 5 min (1 mL/min), and after the wash-out, the skull was opened and the brain carefully removed; brains were stored at −80 °C until homogenization. The brains were weighed, and PBS was added four times. Subsequently, the brains were homogenized by a T-25 Ultra Turrax disperser (IKA, Staufen, Germany), ultrasonicated by a UP 50H needle homogenizer (Hielscher, Teltow, Germany), and stored at -80°C before extraction.

#### 5.8.2. Sample extraction

190 µL of brain homogenate or plasma were spiked with 10 µL of internal standard (IS) isotope-labeled donepezil-d_7_ in methanol (IS, final concentration was 1 µM). The sample was alkalized with 100 µL of 1M sodium hydroxide, and 0.7 mL of tert-butyl methyl ether was added. The samples were then shaken (1,200 RPM, 10 min, Wizard Advanced IR Vortex Mixer, Velp Scientifica, Usmate, Italy) and centrifuged (12,000 RPM, 5 min, Universal 320 R centrifuge, Hettich, Tuttlingen, Germany). 500 µL of supernatant was transferred to a microtube and evaporated to dryness in a CentriVap concentrator (Labconco Corporation, Kansas City, USA). Calibration samples were prepared by the spiking of 180 µL of blank brain homogenate or plasma with 10 µL of the studied compound dissolved in methanol (final concentrations range from 0.5 nM to 50 µM) and with 10 µL of IS, with a final concentration of 1 µM). The calibration samples were then vortexed and extracted as above. The analyzed samples were reconstituted in 100 µL of acetonitrile/water mixture of 50/50 (v/v).

#### 5.8.3. Pharmacokinetic study - HPLC-MS analysis

Compound levels in plasma and brain homogenate were measured by the above-mentioned UHPLC system with mass spectrometric detection. The results were obtained by the gradient elution method with a reverse phase on an RP Amide column (Ascentis Express, 2.1×100 mm, 2.7 µm, Merck, Darmstadt, Germany). Mobile phase A was ultrapure water of ASTM I type (resistance 18.2 MΩ.cm at 25°C) prepared by Barnstead Smart2Pure 3 UV/UF apparatus (Thermo Fisher Scientific, Bremen, Germany) with 0.1% (v/v) formic acid (LC-MS grade, Sigma-Aldrich, Steinheim, Germany); mobile phase B was acetonitrile (MS grade, VWR International, Fontenay-sous-Bois, France) with 0.1% (v/v) of formic acid. The gradient was as follows. Initially, 10% of mobile phase B flowed for 0.3 min; the gradient of mobile phase B then increased to 60% during 2.7 min and then rose to 100% in 0.7 min. After this 0.7 min of 100% B flow, the composition returned to 10 % B and was equilibrated for 3.5 min. The total run time of the gradient elution method was 7.5 min. The rate of flow of the mobile phase was set to 0.45 mL/min, and the column was tempered to 40°C. The injection volume was 5 µL. Samples were analyzed using the previously mentioned Orbitrap mass spectrometer in total ion current in positive mode. The calibration curves had 11 points, and concentrations ranged from 0.5 nM to 50 µM; curves in both cases were linear within the measured range [55].

### 5.9. Acute toxicity evaluation-NOAEL

The acute toxicities of the compounds were assessed at a no-observed-adverse-effect level (NOAEL). Compounds were administered intraperitoneally (i.p., 5% DMSO with 5% Kolliphor in saline) in adult male Wistar rats. Rats were randomly assigned to the experimental groups consisting of four males per one administered dose of tested compounds. Selected doses were administered to identify NOEL and NOAEL with starting dose levels based on their solubility or according to our previous experience. All tested compounds were administered via i.p. injection in a standardized volume of 1 mL.kg-1.

Treated rats were extensively observed for signs of toxicity in the first hour and then periodically for two days. Clinical signs such as cardiovascular, respiratory, and nervous system disability, weight loss, or reduction of food consumption were observed according to Laboratory Animal Science Association (UK) guidelines. The severity of symptoms was classified as mild, moderate, and substantial [61]. During all acute toxicity studies, we followed the rules: i) if a category of substantial severity was achieved, the animal was immediately euthanized by CO_2_, and a lower/more appropriate dose was selected to continue the study; ii) if a severe adverse effect or death occurred within a few minutes after administration to the first animal in the group, the other animal was not treated, and a lower dose was selected as well; iii) in the case of mortality or surviving 48 hours, all animals were euthanized by CO_2_ and subjected to basic macroscopic necropsy [62].

### 5.10. Evaluation of behavioral side effects of 3l and 6f

#### 5.10.1. Animals

Adult male Wistar rats (280–375 g, 2–3 months old) purchased from the Velaz breeding colony were used. The rats were housed in pairs in standard plexiglass boxes (42 × 21 × 20 cm) in the animal facility of the National Institute of Mental Health, Czech Republic, with a 12:12 h light/dark cycle and defined temperature and humidity. Water and food were available *ad libitum*. All experiments were conducted in the light phase of the day after a week-long acclimatization period. Before the experiments, the animals were habituated to manipulation by the experimenter by handling. The experiments were performed in accordance with the guidelines of European Union directive 2010/63/EU and Act No. 246/1992 Coll. on the protection of animals against cruelty and were approved by the Animal Care and Use Committee of the National Institute of Mental Health (reference number MZDR 9928/2023-5/OVZ).

#### 5.10.2. Treatment and experimental design

The compounds 3l and 6f were administered at 1 and 10 mg/kg body weight doses 15 min before the experiment. MK-801 (+)-MK-801 hydrogen maleate, Sigma-Aldrich) was administered at the dose of 0.2 mg/kg body weight (open field test) and 0.3 mg/kg body weight (prepulse inhibition test) 30 min before the experiment. All compounds were first dissolved in DMSO (5% of total vehicle volume; Dimethysulfoxid Rotipuran® p.a., Carl Roth, Germany), and then a solution consisting of 5% Kolliphor EL (Glentham Life Sciences, UK) in physiological saline was added. Control animals received vehicle (5% DMSO in 5% Kolliphor EL in saline) 15 min before the experiment. All drugs were administered intraperitoneally (*i.p.*). Injection volume was 1 ml/kg. The animals were pseudo-randomly assigned to six treatment groups: VEH (vehicle), MK-801, 3l (1), 3l (10), 6f (1), and 6f (10); the numbers in the brackets denote the doses used (1 or 10 mg/kg body weight). We used 8 animals in the VEH group and 6 animals in each group. Each animal underwent an open field test and then, after 3–4 days, a prepulse inhibition test with the same treatment.

#### 5.10.3. Open field test

The apparatus consisted of a square arena (80 × 80 × 40 cm) made of black plastic, located in a separate room with defined light conditions. The animal was placed in the center of the apparatus and allowed to move freely for 10 min. The animal was recorded by a camera above the arena, connected to tracking software (EthoVision XT 16, Noldus, Netherlands). The arena was cleaned thoroughly between animals. The dependent variable was the distance moved by the animal [55].

#### 5.10.4. Prepulse inhibition of acoustic startle response

The startle response is a reflex response to a sudden, intense stimulus. The prepulse inhibition of startle response is a measure of sensorimotor gating and refers to the physiological reduction of the amplitude of the startle response caused by the presentation of a weak stimulus (prepulse) immediately before the startle-inducing stimulus (pulse). During the prepulse inhibition test, acoustic stimuli are presented, and the motor startle responses of the rats are analyzed. The dependent variable is the percent of prepulse inhibition. The experiment was performed as described (the only difference being the time interval between the compound administration and the testing session) [53].

### 5.11. Evaluation of the neuroprotective effect of 3l and 6f

#### 5.11.1. Animals

Experiments were conducted in accordance with the guidelines of the European Union directive 2010/63/EU and approved by the Animal Care and Use Committee, which possesses the National Institutes of Health Statement of Compliance with Standards for Humane Care and Use of Laboratory Animals. Adult male Wistar rats (350–500 g) were used for experiments to assess the neuroprotective activity of compounds 3l and 6f. The rats were kept in a controlled environment described above.

#### 5.11.2. Compounds and experimental groups (treatment)

NMDA was dissolved in 10 mM sterile PBS to achieve a final concentration of 25 mM. Compounds were diluted in a vehicle consisting of sterile 0.9 % physiological saline solution containing 5 % DMSO (Merck Millipore) and 5 % Kolliphor EL to achieve a final concentration of 30 μM. The solutions containing NMDA at 25 mM and compound 3l of 6f at 30 μM were thoroughly mixed in a 1:1 ratio. The control group of animals received a mixed solution of NMDA (25 mM) and vehicle. The second control group received a mixed solution of 10 mM PBS and vehicle. A total of n = 37 animals were used for the experiment ascribed to the following treatment groups: (1) PBS + VEH (received PBS and vehicle; n = 9); (2) NMDA + VEH (received NMDA and vehicle; n = 7); (3) NMDA + 3l (received NMDA and compound 3l; n = 12) and (4) NMDA + 6f (received NMDA and compound 6f; n = 9).

#### 5.11.3. NMDA lesion and histology

The animals underwent surgical preparation under 1.5–2.5 % of isoflurane anesthesia (VetPharma). They were secured onto a stereotaxic apparatus (TSE systems, Berlin, Germany), with their eyes covered with a medical petroleum jelly (Vaseline, Unilever). They were carefully shaved, and the scalp was incised. A unilateral craniotomy was performed at the coordinates of - 4 mm AP and 2.5 mm ML relative to the bregma [63]. The right dorsal hippocampus was injected using a Hamilton 10 μL syringe (DV = 4.6 mm). An infusion pump (model 540310 plus, TSE Systems, Berlin, Germany) maintained a constant flow rate of 0.25 μL/min to apply the mixed solutions. The total volume administered to each animal was 2 μL. After surgery, the animals had free access to food and water containing analgesics. At 24 h after the application of the mixed NMDA and compounds 3l or 6f solutions, the rats were euthanized with an overdose of the anesthetics Narketan (ketamine; 100 mg/kg) and Rometar (xylazine; 20 mg/kg). Transcranial perfusion was performed with 4% paraformaldehyde (PFA) diluted in 0.2 M phosphate buffer (PB). Brains were post-fixed 24 h in 4% PFA and gradually cryoprotected using increasing concentrations of sucrose solution (10, 20, and 30%; the brain was immersed in the solution until it sank to the bottom of the tube). Frozen cryoprotected brains were sliced in the coronal plane (50 μm, 1-in-5 series) from the beginning of the hippocampus (Bregma – 1.80 mm). All sections were collected and stored at -20°C in cryoprotectant solution (ethylene glycol, glycerol, 0.2 M PB, distilled water - 3:3:1:3). Each series of slices underwent evaluation of the neurodegenerative damage of the hippocampus. The severity of hippocampal damage was determined in the following hippocampal regions: granular cell layer of the lower part of the dentate gyrus (DGl), granular cell layer of the higher part of the dentate gyrus (DGh), hilus, CA1, and CA3, as previously described [64].

### 5.12. Statistics

Electrophysiological data are presented as mean ± standard error of the mean (SEM), and group differences were analyzed by t-test using SigmaStat 3.5 (Systat Software Inc.). Differences with a p-value < 0.05 were considered statistically significant. The behavioral data were analyzed in GraphPad Prism 8 (San Diego, USA) using ANOVA with Dunnett’s multiple comparisons test (vs. VEH group) and presented as mean + SEM. p-value < 0.05 was considered statistically significant. In the data from prepulse inhibition, one outlier in the 6f (10) group was removed (Grubbs test). The data concerning the neuroprotective effect of the compounds were also analyzed in GraphPad Prism 8 (San Diego, USA) and presented as mean + SEM; p < 0.05 was considered statistically significant. These data were assessed for normality (Shapiro-Wilk test) and homogeneity of variances (Brown-Forsythe test). As these data did not meet the assumption of homogeneity of variances, Welch’s ANOVA was employed, followed by Dunnett’s T3 multiple comparisons test.

## Supporting information

Pharmacokinetics, blood brain permeability, chemistry (purity)

## CRediT authorship contribution statement

**Jan Konecny:** Investigation, Conceptualization, Methodology, Writing – review & editing. **Anna Misiachna:** Investigation, Conceptualization, Methodology, Data curation, Writing – review & editing. **Marketa Chvojkova:** Methodology, Investigation, Data curation. **Lenka Kleteckova:** Methodology, Investigation, Data curation. **Marharyta Kolcheva:** Methodology, Investigation. **Martin Novak:** Methodology, Investigation. **Lukas Prchal:** Methodology, Investigation. **Marek Ladislav:** Methodology, Data curation. **Katarina Hemelikova:** Methodology. **Jakub Netolicky:** Investigation. **Tereza Kobrlova:** Investigation, Data curation. **Jana Zdarova Karasova:** Conceptualization, Investigation, Data curation. **Jaroslav Pejchal:** Investigation, Data curation. **Pavla Jendelova:** Writing – original draft, Conceptualization. **Yuan-Ping Pang:** Conceptualization. **Karel Vales:** Writing – review & editing, Supervision, Funding acquisition. **Jan Korabecny:** Writing – review & editing, Validation, Supervision, Funding acquisition. **Ondrej Soukup:** Writing – review & editing, Writing – original draft, Supervision, Project administration, Methodology, Funding acquisition, Conceptualization. **Martin Horak:** Writing – review & editing, Writing – original draft, Supervision, Funding acquisition, Formal analysis, Data curation, Conceptualization.

## Declaration of competing interest

The authors declare that they have no known competing financial interests or personal relationships that could have appeared to influence the work reported in this paper.

## Data availability

Data will be made available on request.

## Acknowledgment

Supported by the Czech Science Foundation (24-10026S), the Ministry of Health of the Czech Republic (grant number NU20-08-00296), and project registration number CZ.02.01.01/00/22_008/0004562 (ExRegMed, MEYS CR). Supported by MH CZ – DRO (UHHK, 00179906).

## Notes

### Competing Interest Statement

The authors have declared no competing interest.

