## Supplementary material for "Dizocilpine derivatives with neuroprotective effect lacking the psychomimetic side effects": Pharmacokinetics, blood brain permeability, chemistry (purity)

**Table S1**. Physicochemical parameters of compounds **3a-o, 6a-l** and memantine.

| Compound | **MW** | **pKa** | **logP** | **HBA** | **HBD** | **TPSA** |
|---|---|---|---|---|---|---|
| **3a** | 223.31 | 8.90 | 3.88 | 1 | 1 | 12.03 |
| **3b** | 237.34 | 8.97 | 4.24 | 1 | 1 | 12.03 |
| **3c** | 251.37 | 9.06 | 4.65 | 1 | 1 | 12.03 |
| **3d** | 249.35 | 8.65 | 4.34 | 1 | 1 | 12.03 |
| **3e** | 265.39 | 9.17 | 5.20 | 1 | 1 | 12.03 |
| **3f** | 265.39 | 9.30 | 5.12 | 1 | 1 | 12.03 |
| **3g** | 291.43 | 9.37 | 5.68 | 1 | 1 | 12.03 |
| **3h** | 277.40 | 9.84 | 5.11 | 1 | 0 | 3.24 |
| **3i** | 279.38 | 7.64 | 4.04 | 2 | 0 | 12.47 |
| **3j** | 267.37 | 8.36 | 3.83 | 2 | 1 | 21.26 |
| **3k** | 263.38 | 9.96 | 4.67 | 1 | 0 | 3.24 |
| **3l** | 253.34 | 8.42 | 3.19 | 2 | 2 | 32.26 |
| **3m** | 292.42 | 8.46 | 4.11 | 2 | 0 | 6.48 |
| **3n** | 265.39 | 9.54 | 4.98 | 1 | 0 | 3.24 |
| **3o** | 306.44 | 8.51 | 4.47 | 2 | 0 | 6.48 |
| **6a** | 221.30 | 8.78 | 3.68 | 1 | 1 | 12.03 |
| **6b** | 235.32 | 8.86 | 4.04 | 1 | 1 | 12.03 |

|  |  |  |  |  |  |  |
|---|---|---|---|---|---|---|
| **6c** | 249.35 | 8.94 | 4.46 | 1 | 1 | 12.03 |
| **6d** | 247.33 | 8.54 | 4.15 | 1 | 1 | 12.03 |
| **6e** | 263.38 | 9.06 | 5.01 | 1 | 1 | 12.03 |
| **6f** | 263.38 | 9.19 | 4.93 | 1 | 1 | 12.03 |
| **6g** | 289.41 | 9.25 | 5.48 | 1 | 1 | 12.03 |
| **6h** | 275.39 | 9.61 | 4.92 | 1 | 0 | 3.24 |
| **6i** | 277.36 | 7.41 | 3.85 | 2 | 0 | 12.47 |
| **6j** | 265.35 | 8.25 | 3.64 | 2 | 1 | 21.26 |
| **6k** | 261.36 | 9.73 | 4.47 | 1 | 2 | 3.24 |
| **6l** | 251.32 | 8.31 | 2.99 | 2 | 2 | 32.26 |
| **Memantine** | 179.30 | 10.70 | 2.07 | 1 | 1 | 26.02 |

MW = molecular weight, pKa = acid dissociation constant, logP = partition coefficient, HBA = hydrogen bond acceptor, HBB = hydrogen bond donor, TPSA = topological polar surface area

**Table S2.** Drug-likeness of derivatives **3a-o**, **6a-l** and memantine.

|  | **Drug-likeness** |  |  |  |  |
|---|---|---|---|---|---|
| **Compound** | **Lipinski** [a] | **Ghose** [b] | **Veber** [c] | **Egan** [d] | **Muegge** [e] |
| **3a** | Yes | Yes | Yes | Yes | No [f] |
| **3b** | Yes | Yes | Yes | Yes | No [f] |
| **3c** | Yes | Yes | Yes | Yes | No [f] |
| **3d** | Yes | Yes | Yes | Yes | No [f] |
| **3e** | No | Yes | Yes | Yes | No [f] |
| **3f** | No | Yes | Yes | Yes | No [f] |
| **3g** | No | Yes | Yes | Yes | No [g] |
| **3h** | No | Yes | Yes | Yes | No [f] |
| **3i** | Yes | Yes | Yes | Yes | Yes |

| | | | | | |
|---|---|---|---|---|---|
| **3j** | Yes | Yes | Yes | Yes | Yes |
| **3k** | No | Yes | Yes | Yes | No [f] |
| **3l** | Yes | Yes | Yes | Yes | Yes |
| **3m** | Yes | Yes | Yes | Yes | Yes |
| **3n** | No | Yes | Yes | Yes | No [f] |
| **3o** | Yes | Yes | Yes | Yes | Yes |
| **6a** | Yes | Yes | Yes | Yes | No [f] |
| **6b** | Yes | Yes | Yes | Yes | No [f] |
| **6c** | Yes | Yes | Yes | Yes | No [f] |
| **6d** | Yes | Yes | Yes | Yes | No [f] |
| **6e** | Yes | Yes | Yes | Yes | No [f] |
| **6f** | Yes | Yes | Yes | Yes | No [f] |
| **6g** | No | Yes | Yes | Yes | No [g] |
| **6h** | No | Yes | Yes | Yes | No [f] |
| **6i** | Yes | Yes | Yes | Yes | Yes |
| **6j** | Yes | Yes | Yes | Yes | Yes |
| **6k** | Yes | Yes | Yes | Yes | No [f] |
| **6l** | Yes | Yes | Yes | Yes | Yes |
| **Memantine** | Yes | Yes | Yes | Yes | No [g] |

[a] Lipinski represent,[1] MW < 500 Da, logP <5, HBD <5, HBA <10; [b] Ghose,[2] MW = 180 – 480, MR (molar refractivity) = 40 – 130, logP = -0.4 – +5.6, number of atoms = 20 – 70; [c] Veber,[3] rotatable bonds <10, TPSA <140; [d] Egan,[4] TPSA <132, logP <5.88; [e] Muegge,[5] MW = 200 – 500, log P = -2 - +5, HBD <5, HBA <10, MR = 40 – 130, rotatable bonds <8, heavy atoms = 20 – 70, TPSA <120, net charge -2 - +2; [f] 1 violation, [g] 2 violations.

### 1. [1]H and [13]C NMR spectra

*N*-methyl-10,11-dihydro-5*H*-dibenzo[*a,d*][7]annulen-5-amine hydrochloride (**3a**):

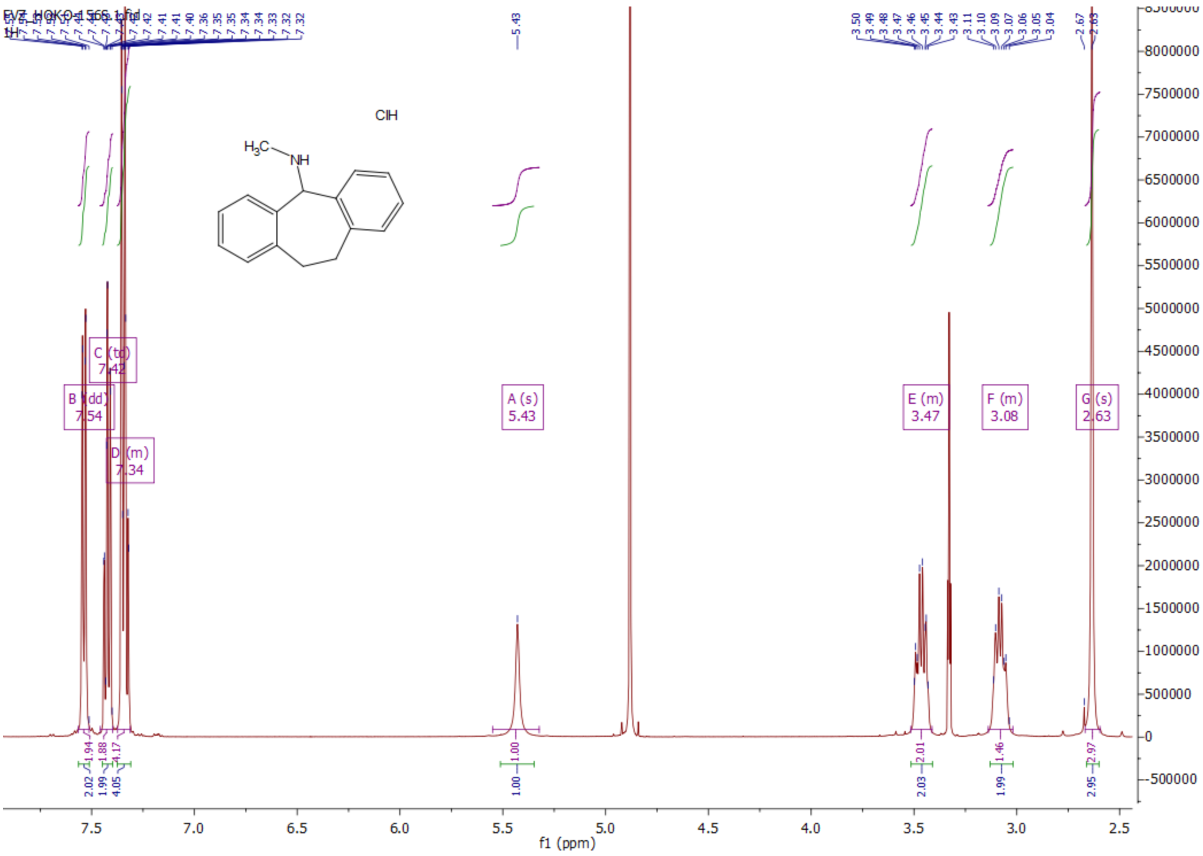

*N*-methyl-10,11-dihydro-5*H*-dibenzo[*a,d*][7]annulen-5-amine hydrochloride (**3a**):

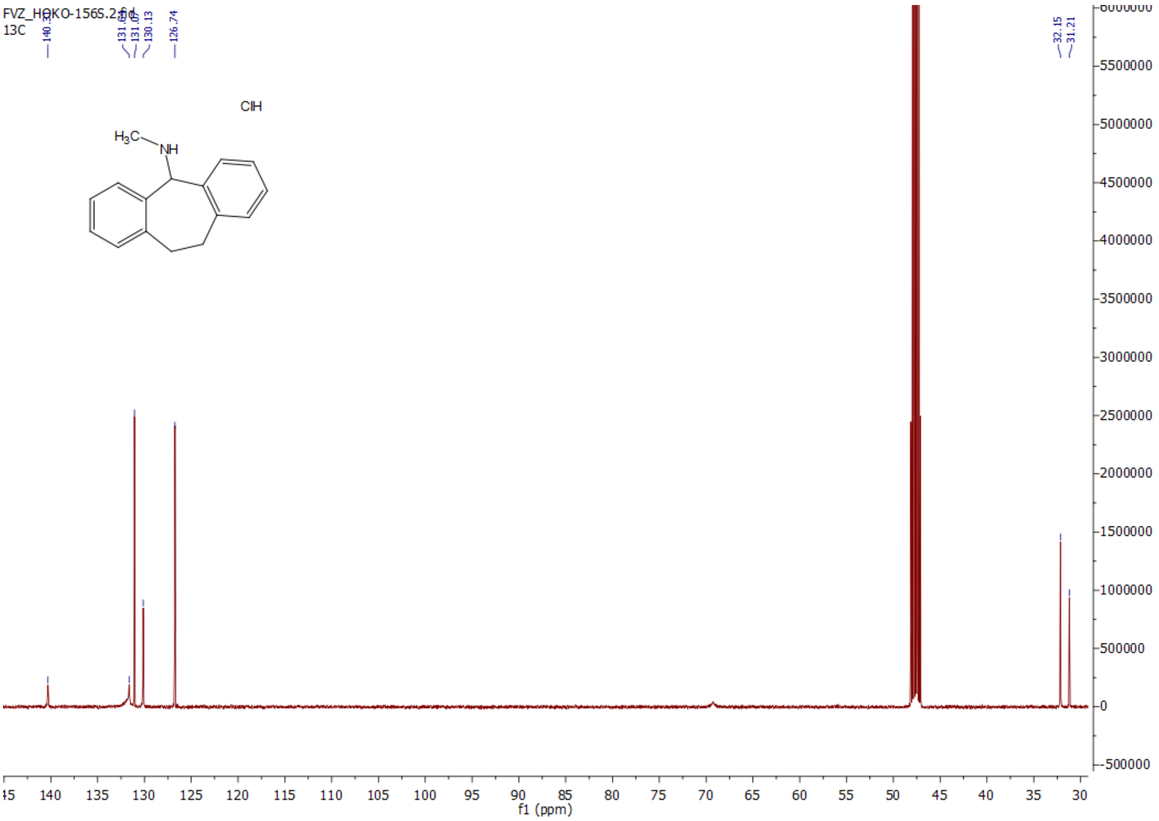


*N*-ethyl-10,11-dihydro-5*H*-dibenzo[*a,d*][7]annulen-5-amine hydrochloride (**3b**):

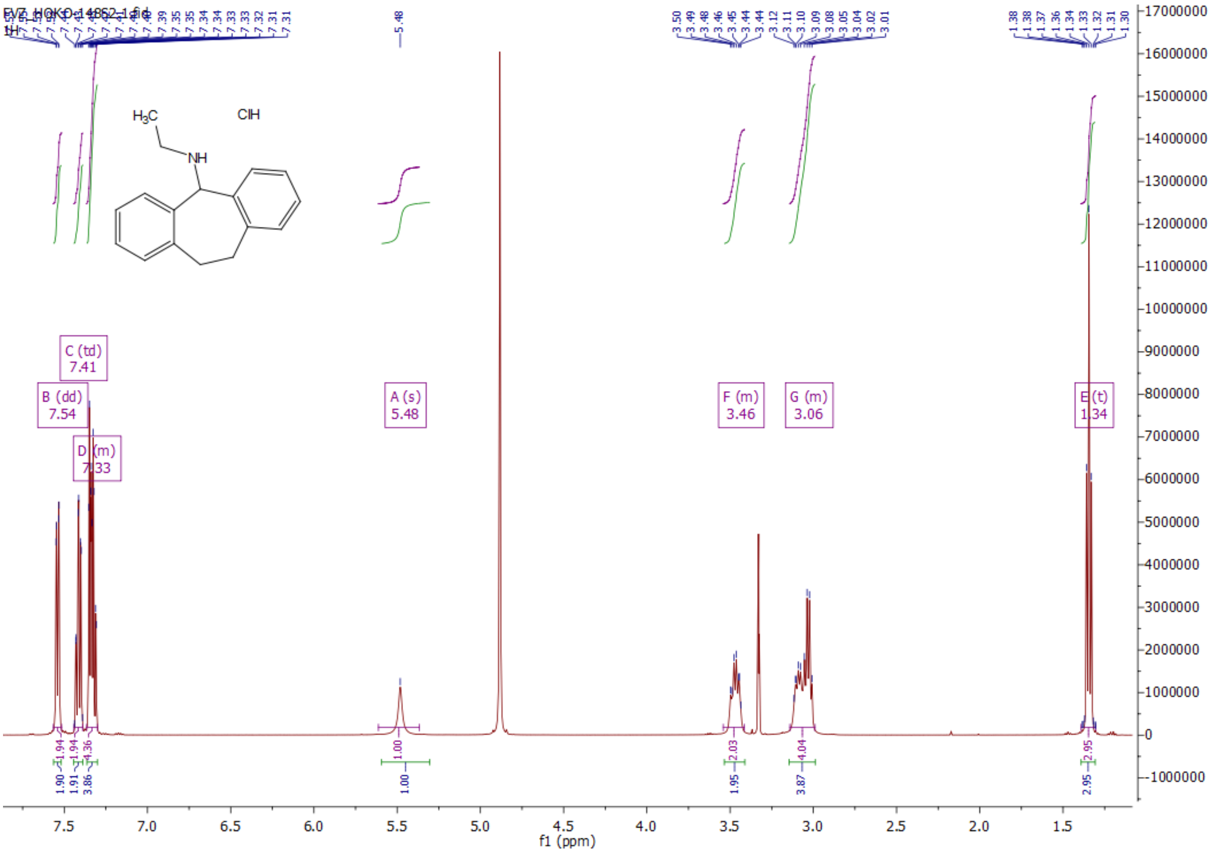


*N*-ethyl-10,11-dihydro-5*H*-dibenzo[*a,d*][7]annulen-5-amine hydrochloride (**3b**):

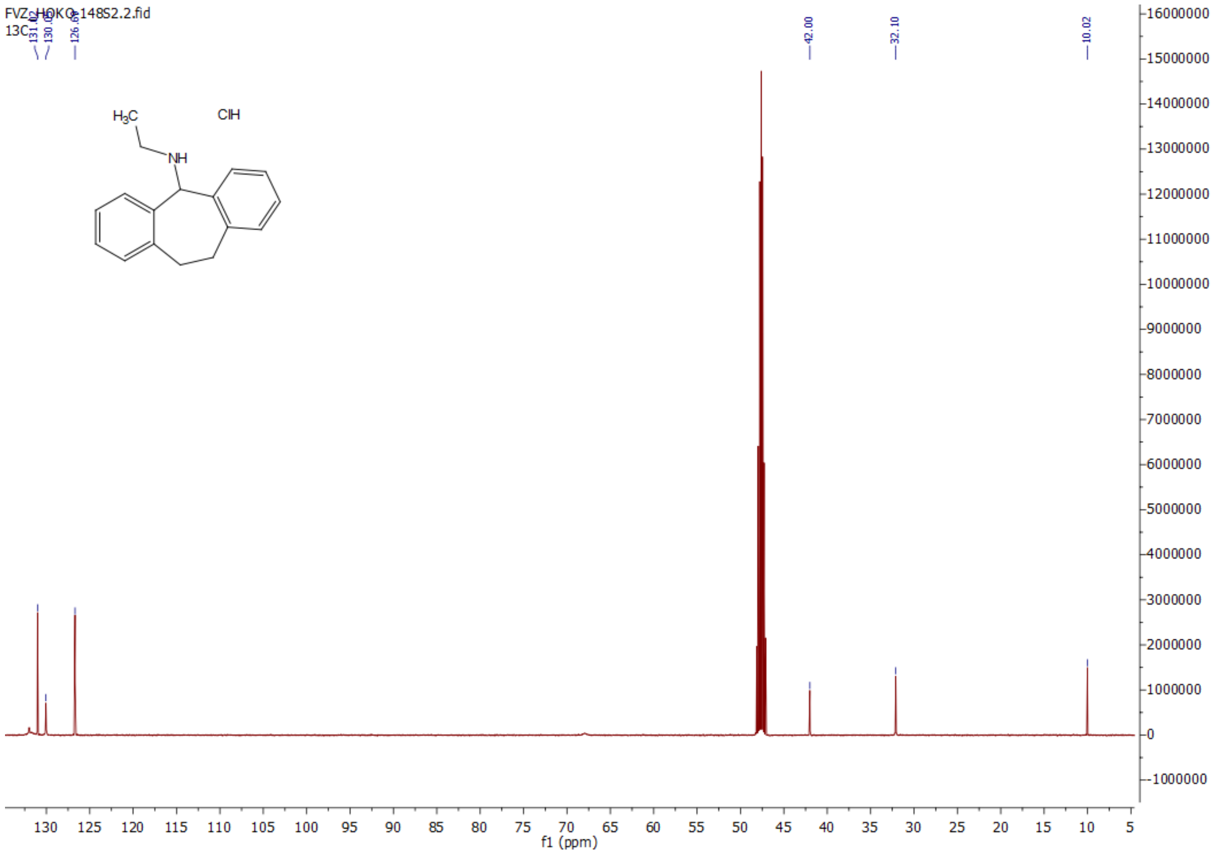


*N*-isopropyl-10,11-dihydro-5*H*-dibenzo[*a,d*][7]annulen-5-amine hydrochloride (**3c**):

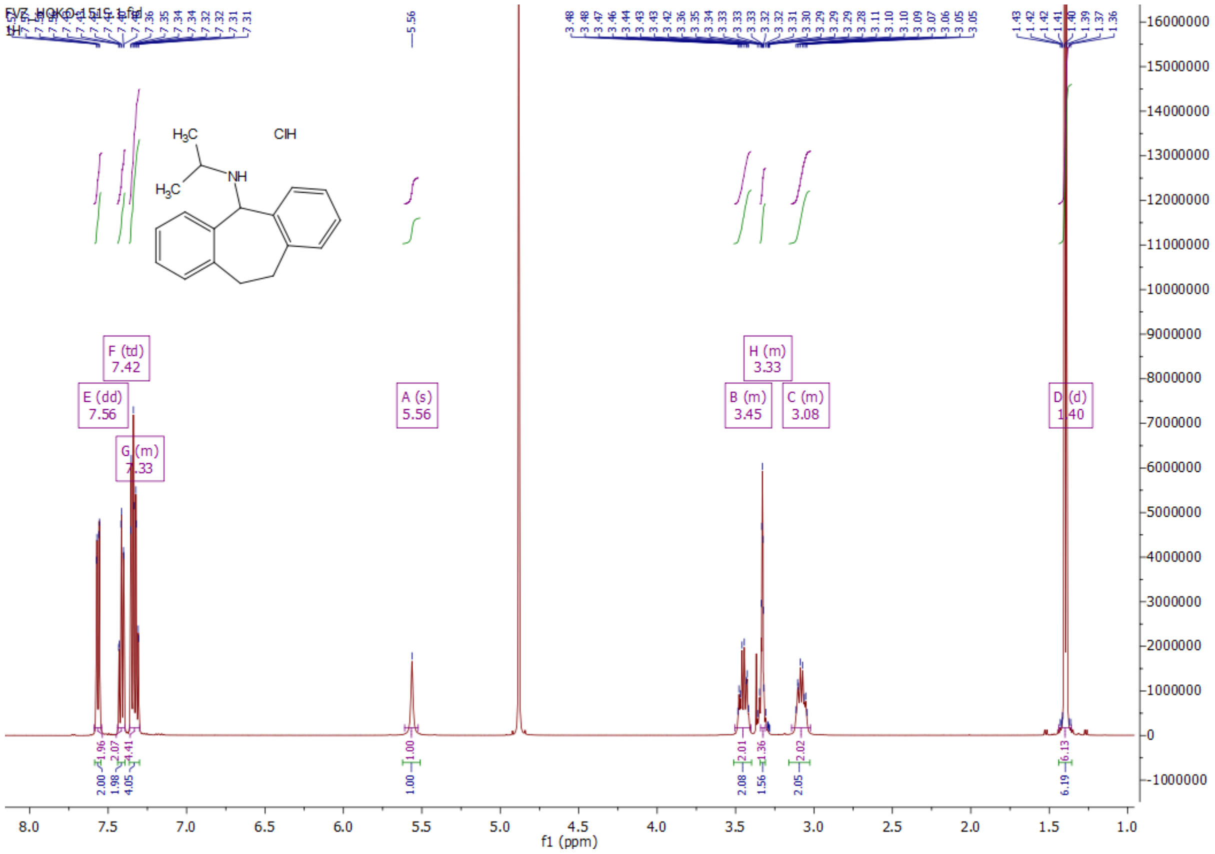


*N*-isopropyl-10,11-dihydro-5*H*-dibenzo[*a,d*][7]annulen-5-amine hydrochloride (**3c**):

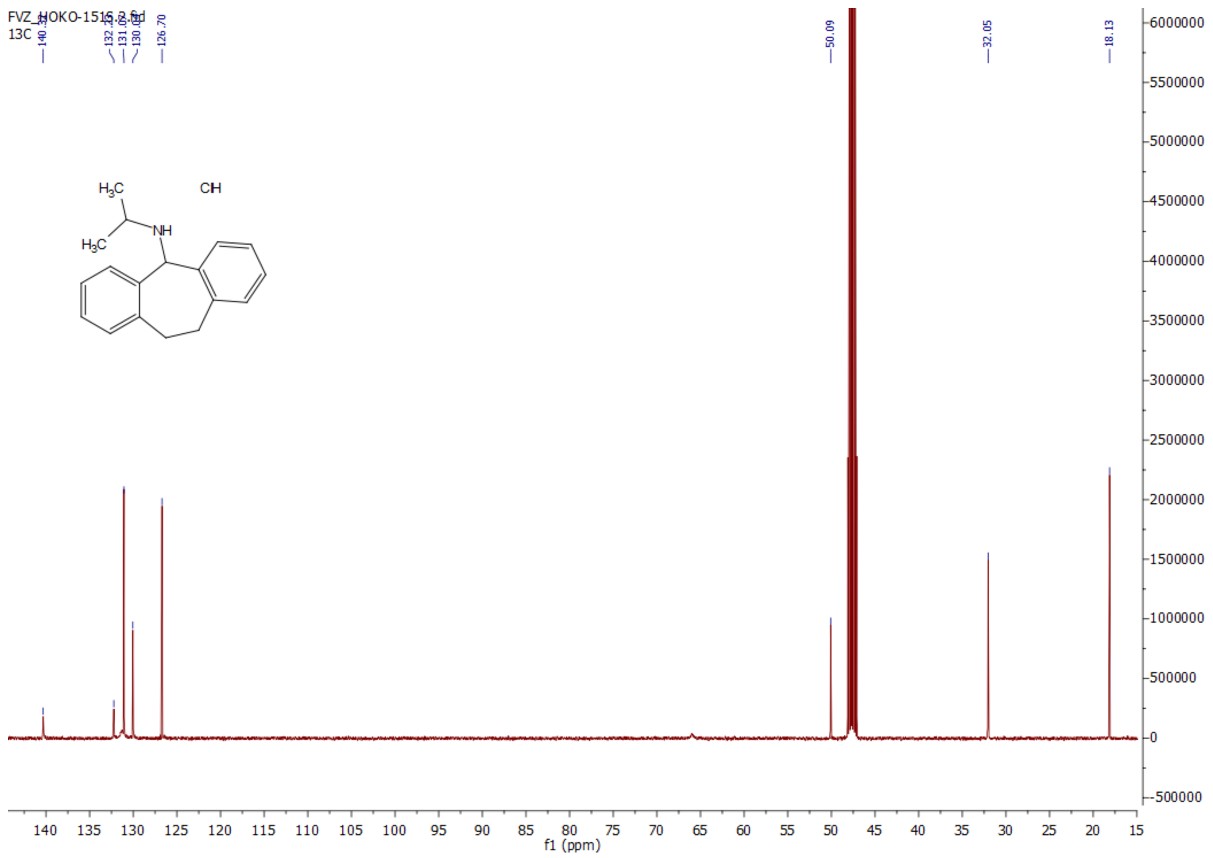


*N*-cyclopropyl-10,11-dihydro-5*H*-dibenzo[*a,d*][7]annulen-5-amine hydrochloride (**3d**):

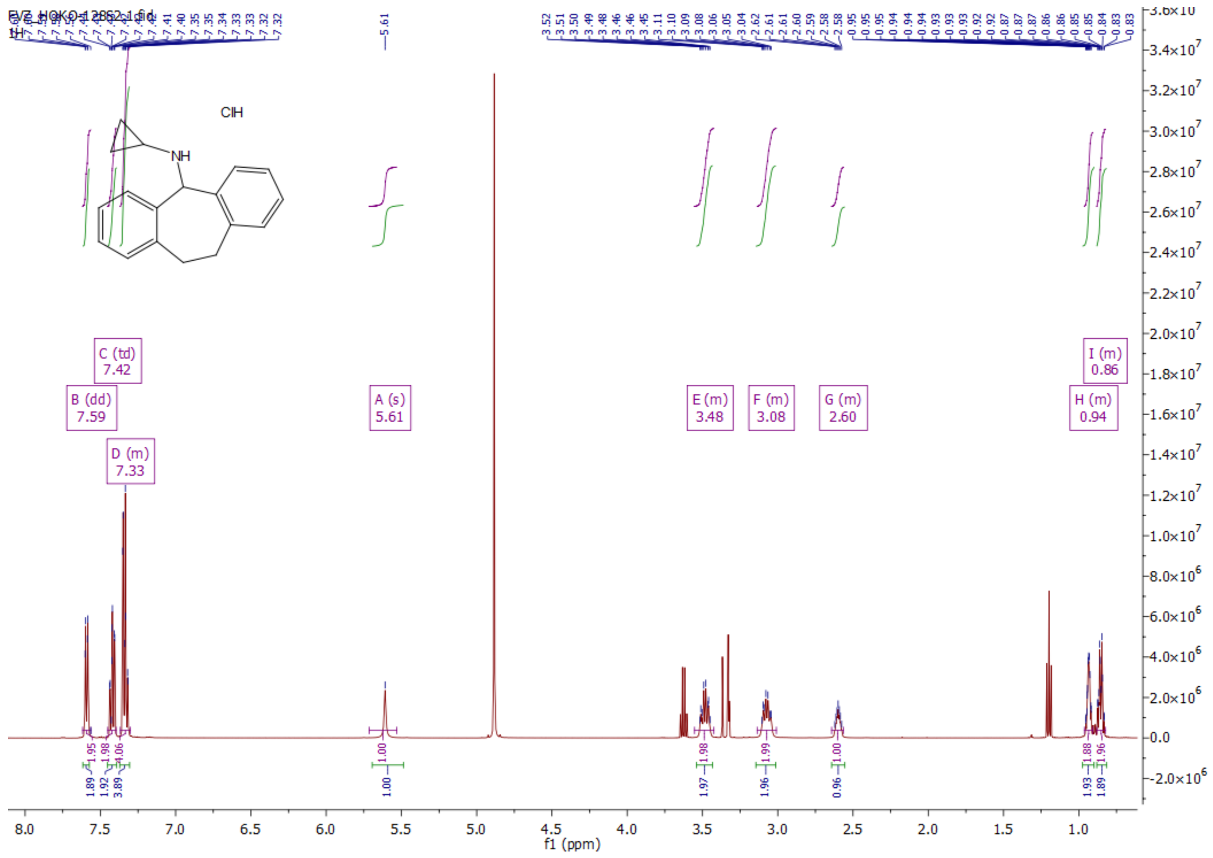


*N*-cyclopropyl-10,11-dihydro-5*H*-dibenzo[*a,d*][7]annulen-5-amine hydrochloride (**3d**):

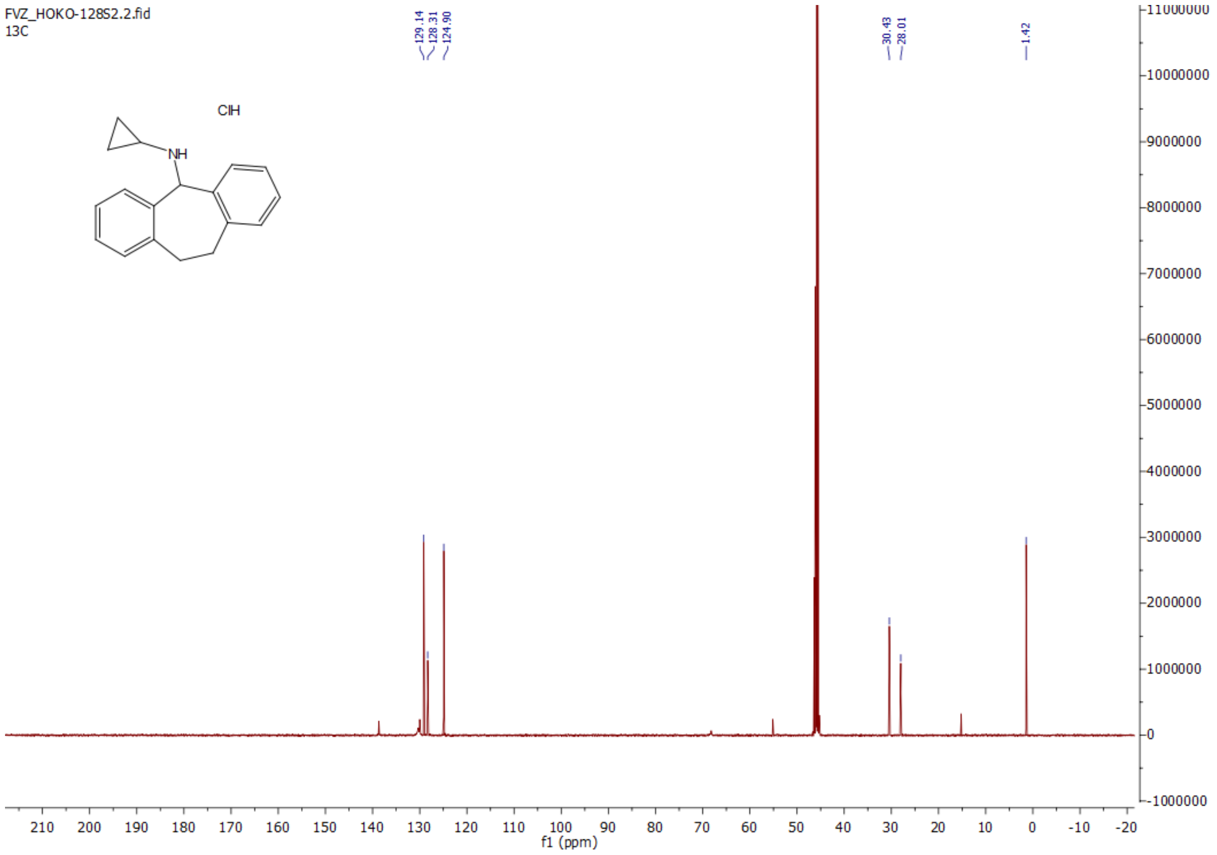


*N*-butyl-10,11-dihydro-5*H*-dibenzo[*a,d*][7]annulen-5-amine hydrochloride (**3e**):

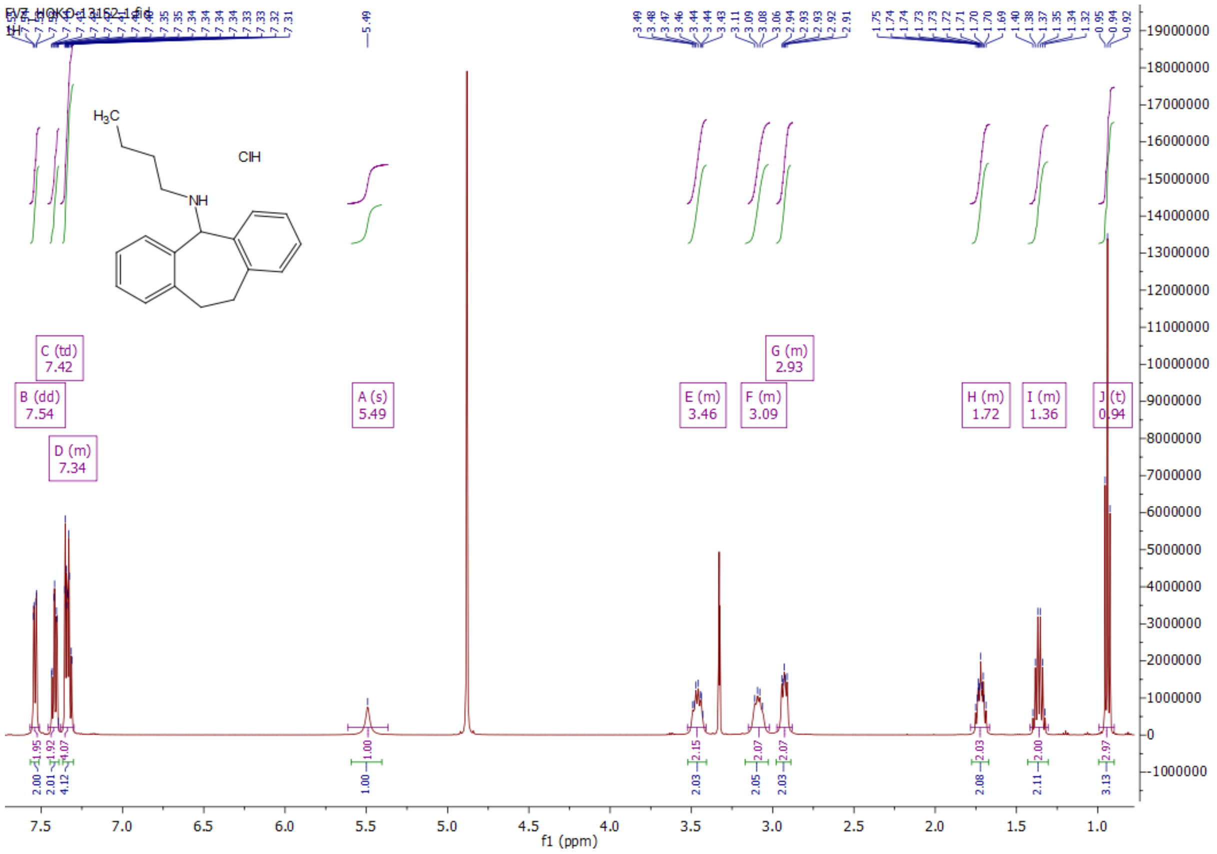


*N*-butyl-10,11-dihydro-5*H*-dibenzo[*a,d*][7]annulen-5-amine hydrochloride (**3e**):

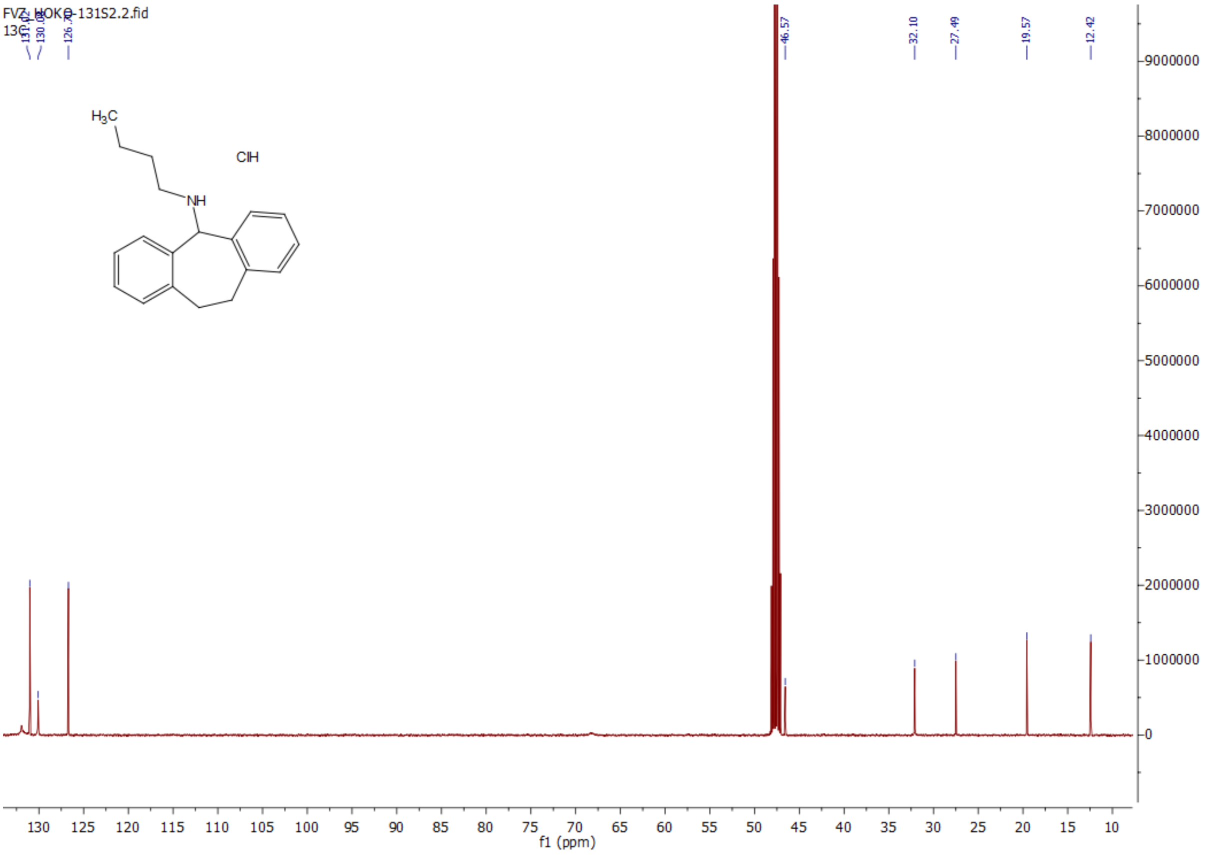


*N*-isobutyl-10,11-dihydro-5*H*-dibenzo[*a*,*d*][7]annulen-5-amine hydrochloride (**3f**):

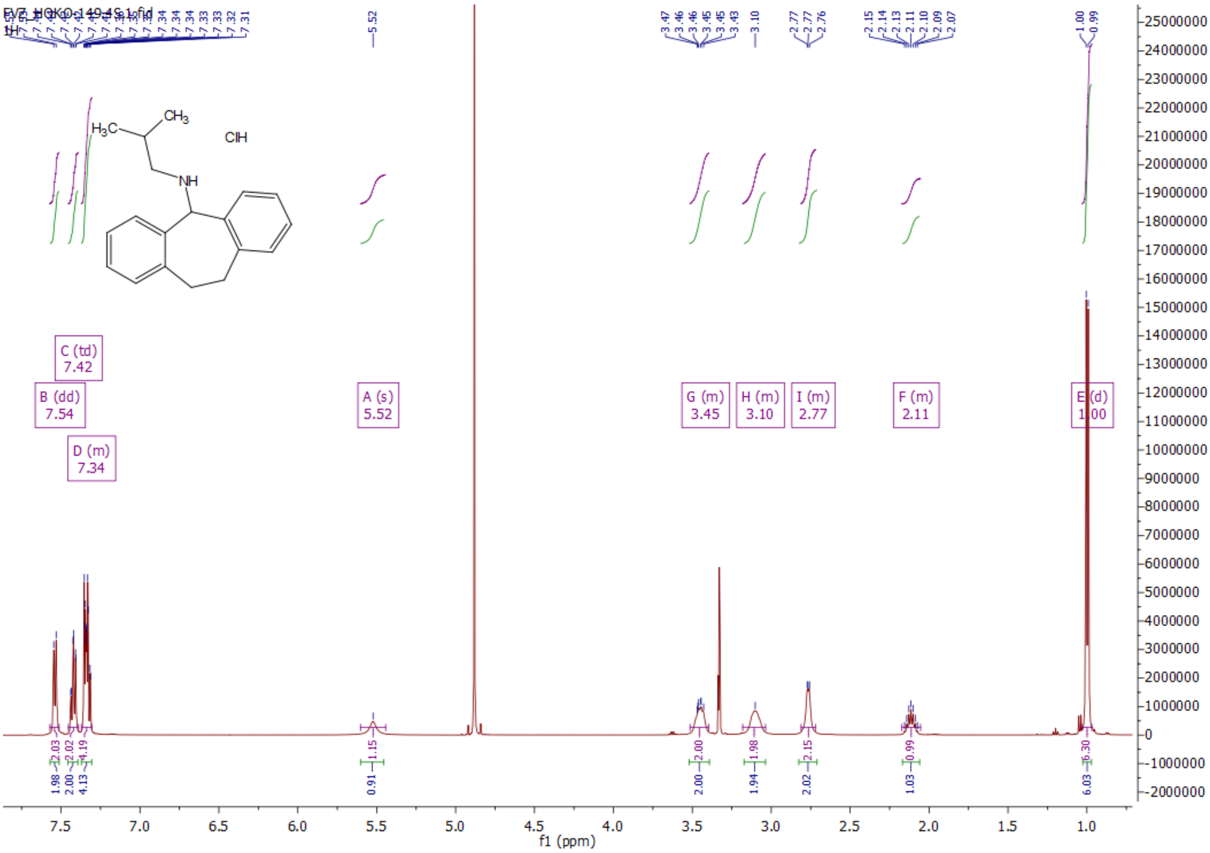


*N*-isobutyl-10,11-dihydro-5*H*-dibenzo[*a*,*d*][7]annulen-5-amine hydrochloride (**3f**):

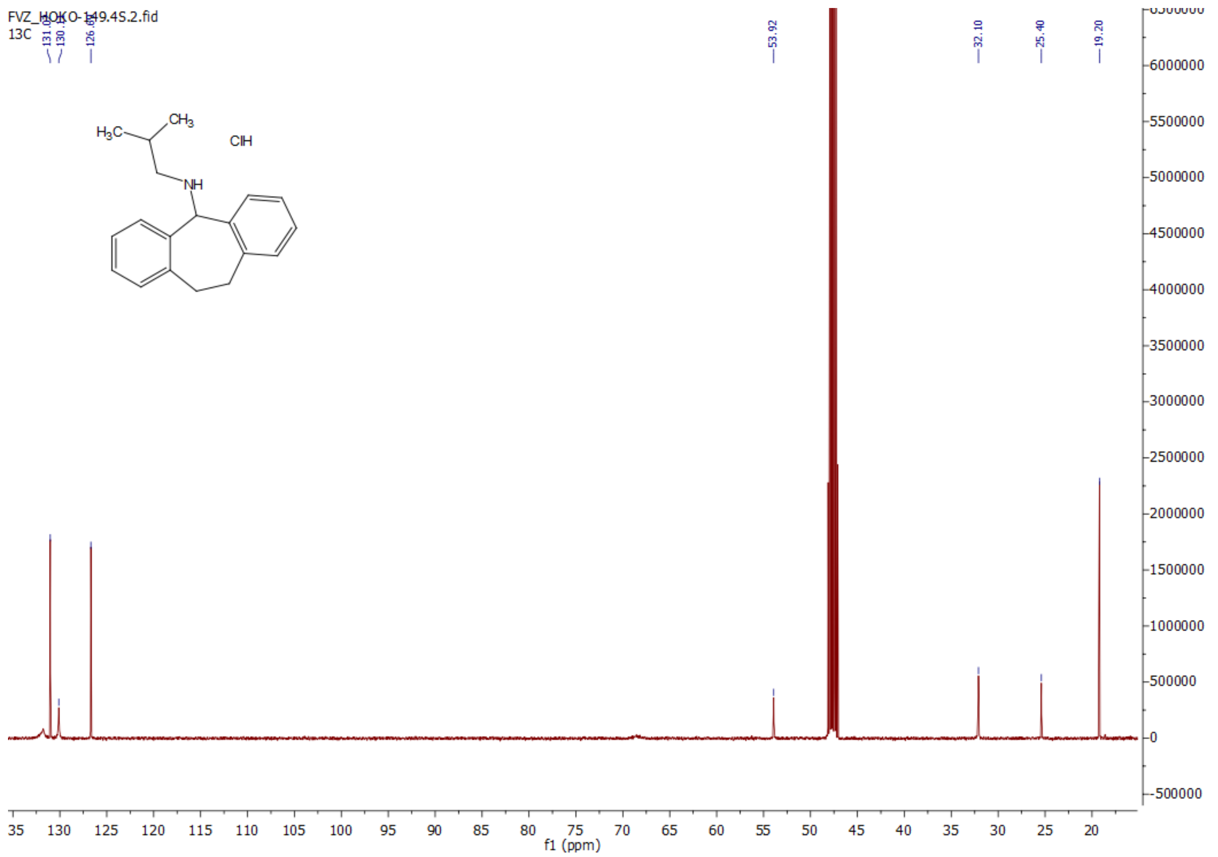

*N*-cyclohexyl-10,11-dihydro-5*H*-dibenzo[*a,d*][7]annulen-5-amine hydrochloride (**3g**):

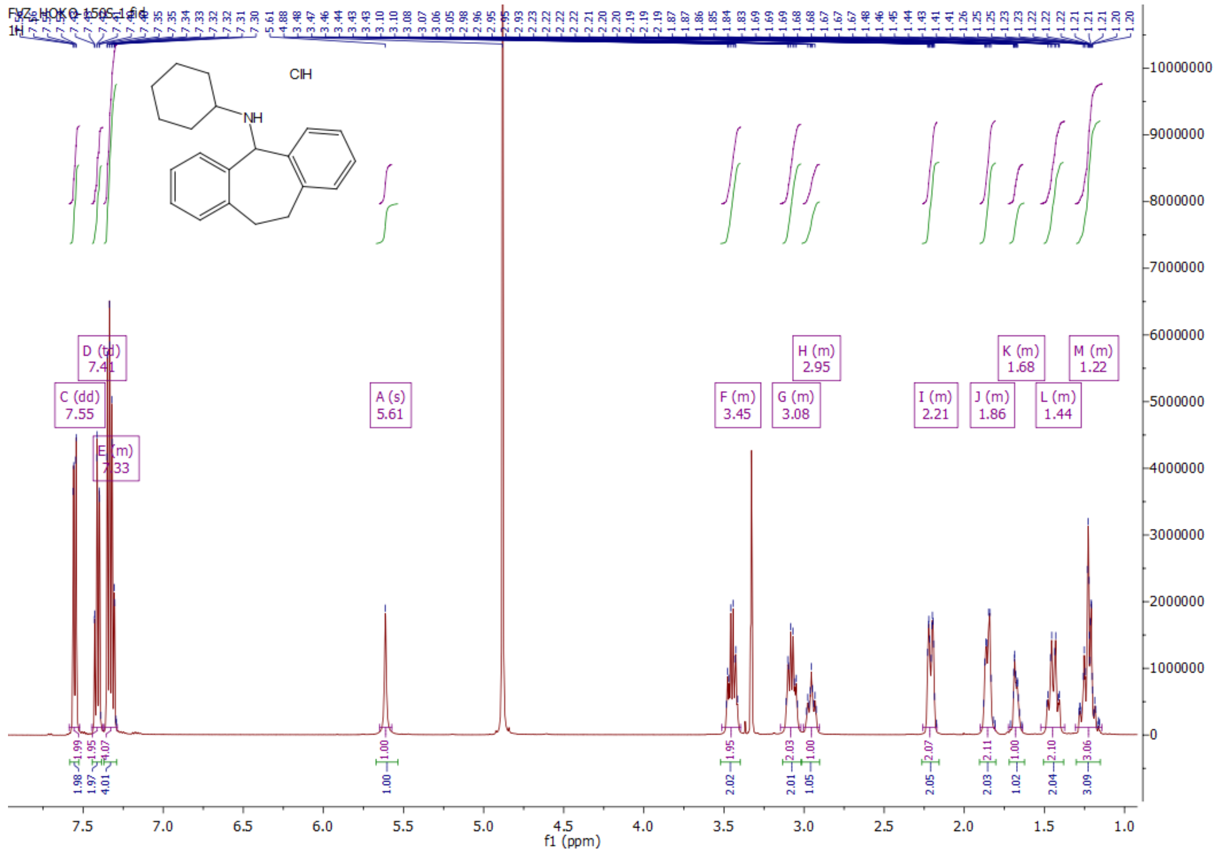

*N*-cyclohexyl-10,11-dihydro-5*H*-dibenzo[*a,d*][7]annulen-5-amine hydrochloride (**3g**):

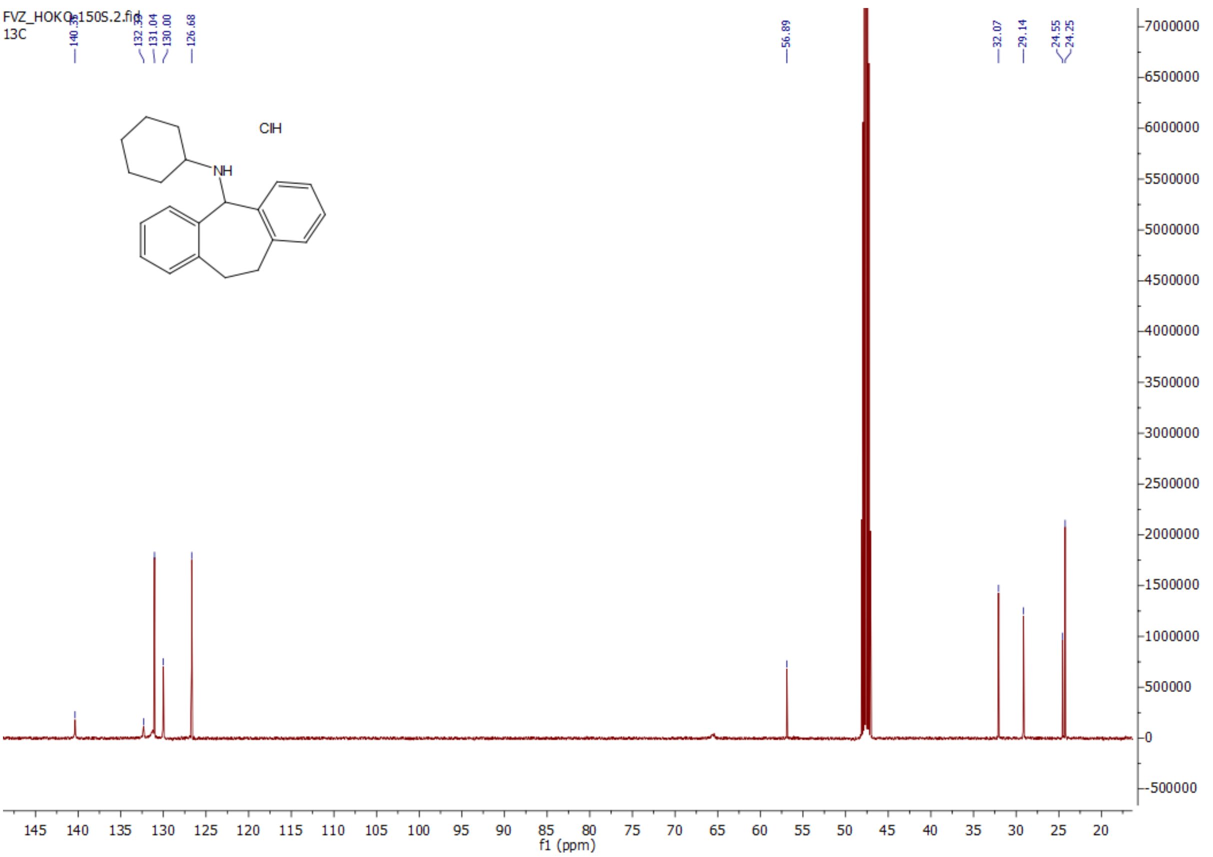


1-(10,11-dihydro-5*H*-dibenzo[*a,d*][7]annulen-5-yl)piperidine hydrochloride (**3h**):

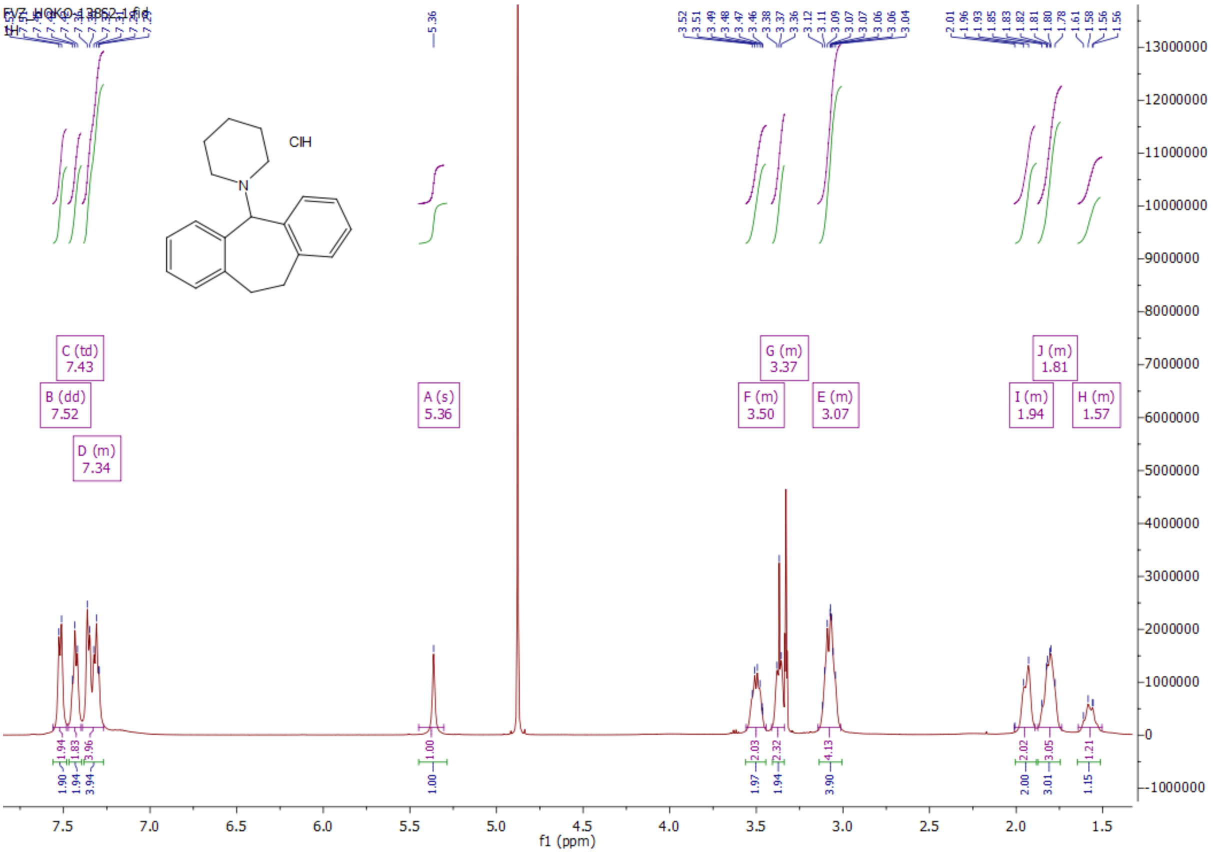


1-(10,11-dihydro-5*H*-dibenzo[*a,d*][7]annulen-5-yl)piperidine hydrochloride (**3h**):

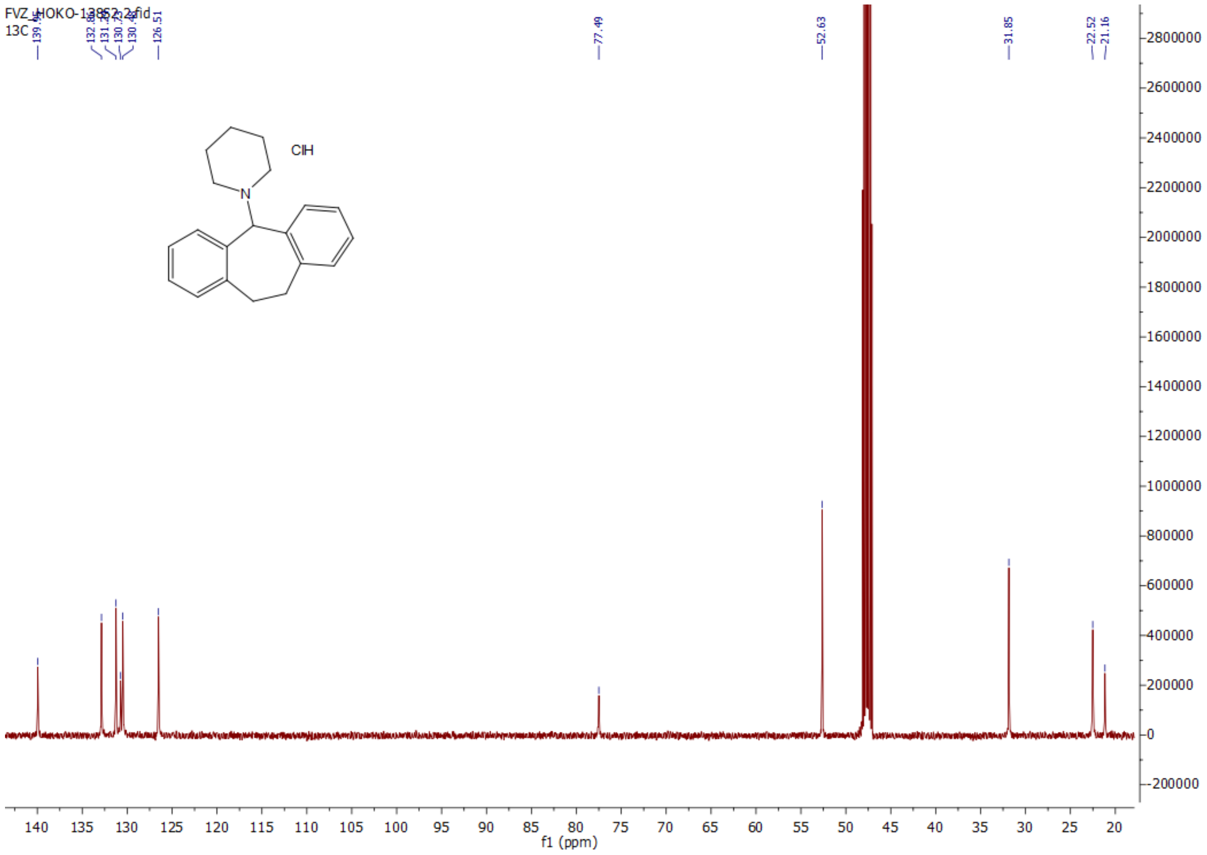


4-(10,11-dihydro-5*H*-dibenzo[*a,d*][7]annulen-5-yl)morpholine hydrochloride (**3i**):

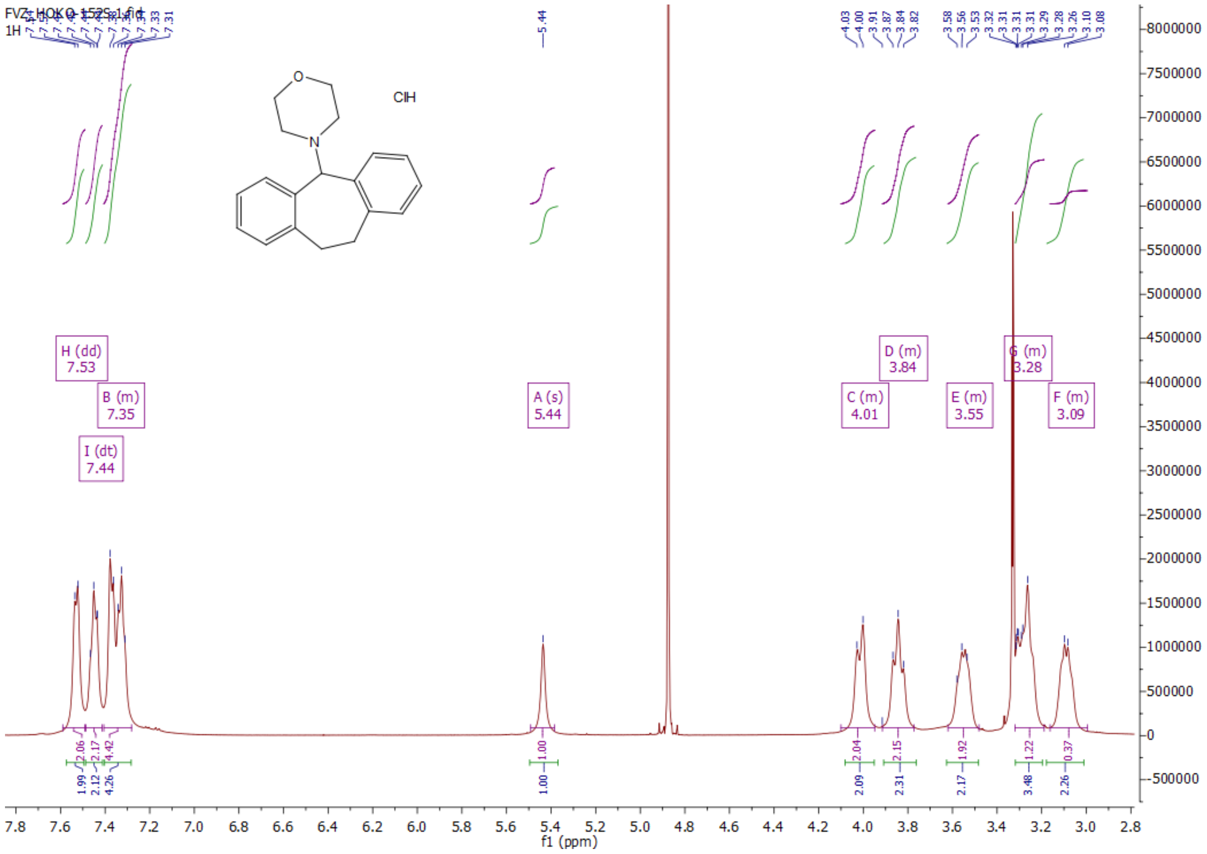


4-(10,11-dihydro-5*H*-dibenzo[*a,d*][7]annulen-5-yl)morpholine hydrochloride (**3i**):

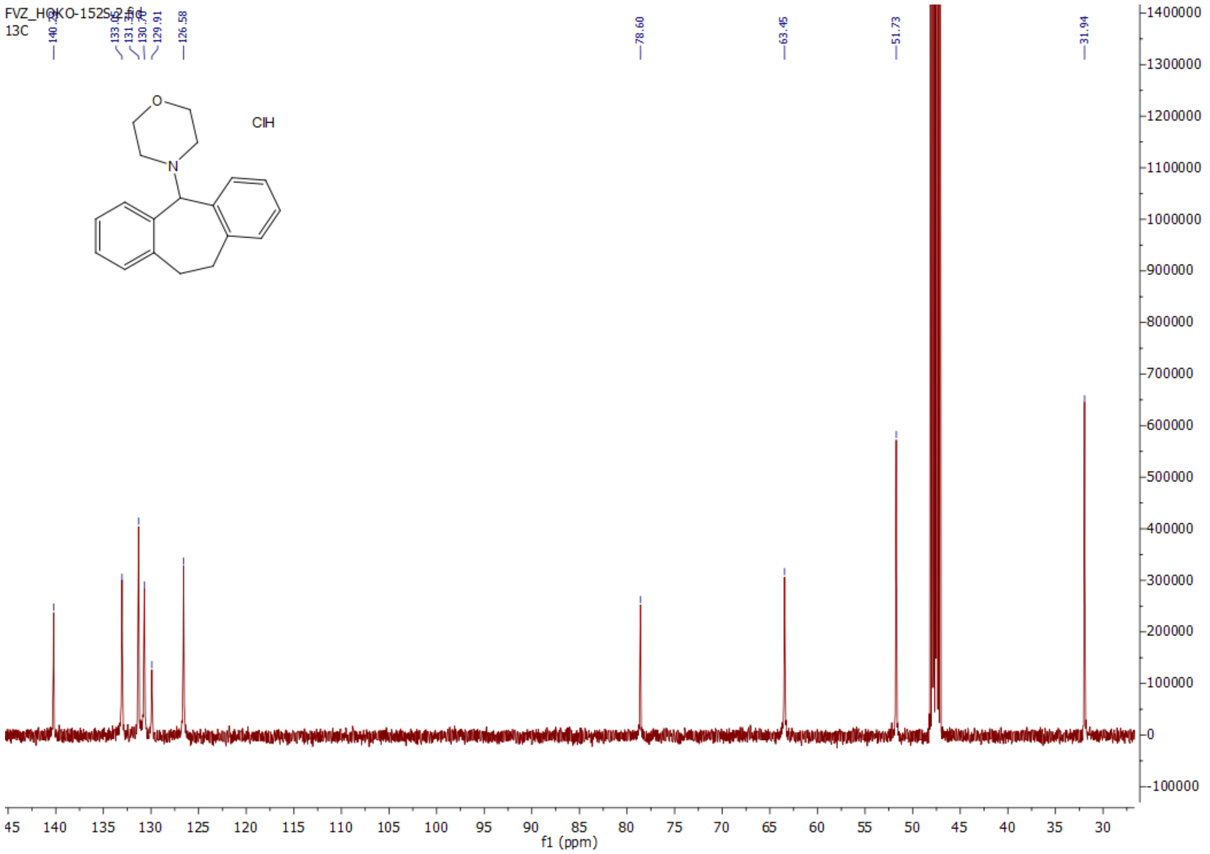


*N*-(2-methoxyethyl)-10,11-dihydro-5*H*-dibenzo[*a,d*][7]annulen-5-amine hydrochloride (**3j**):

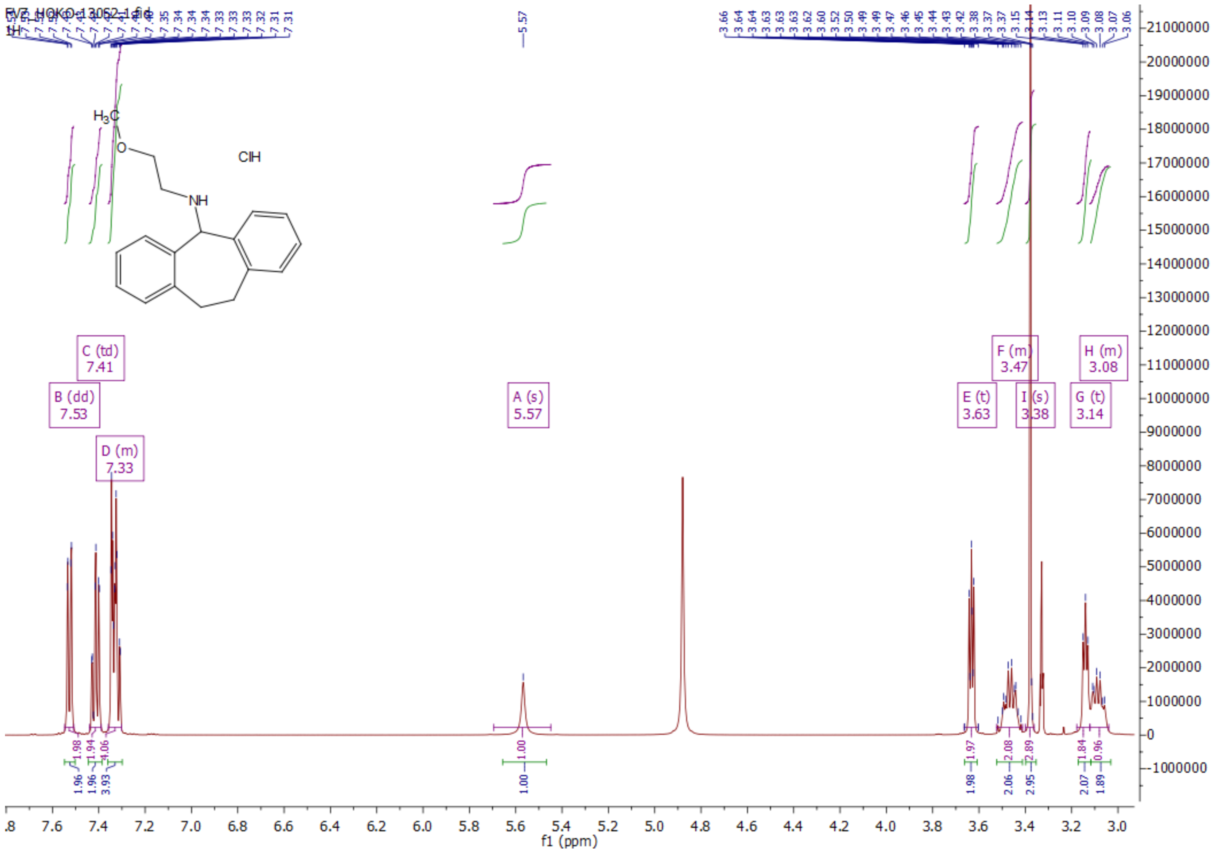

*N*-(2-methoxyethyl)-10,11-dihydro-5*H*-dibenzo[*a,d*][7]annulen-5-amine hydrochloride (**3j**):

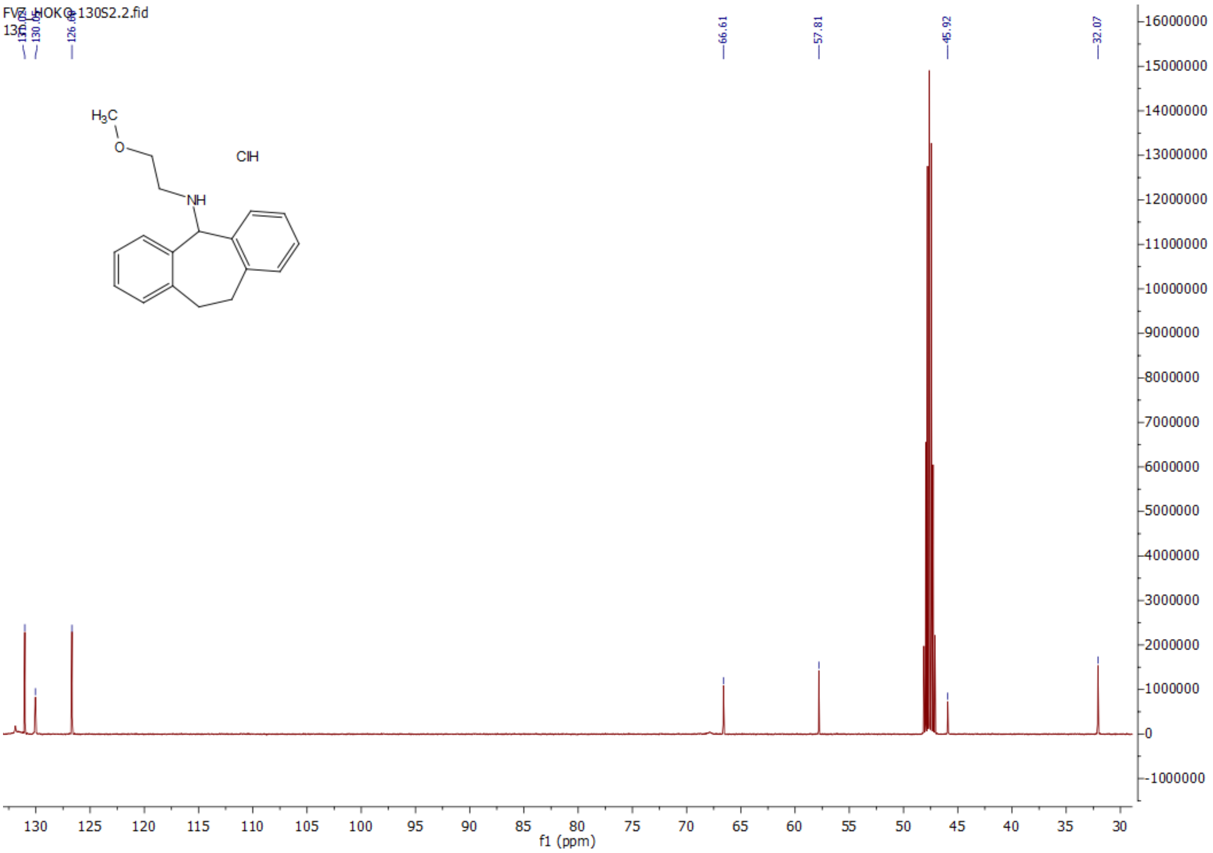

1-(10,11-dihydro-5*H*-dibenzo[*a,d*][7]annulen-5-yl)pyrrolidine hydrochloride (**3k**):

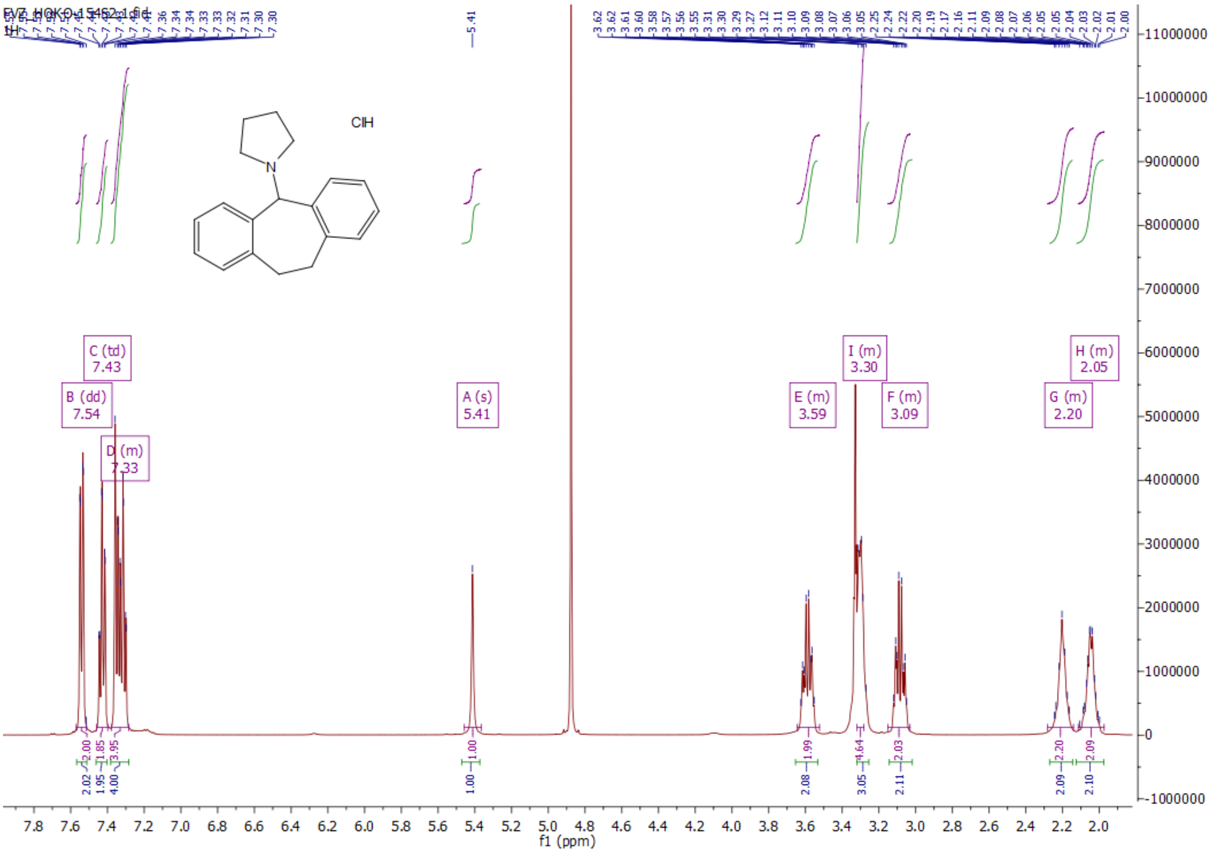

1-(10,11-dihydro-5*H*-dibenzo[*a,d*][7]annulen-5-yl)pyrrolidine hydrochloride (**3k**):

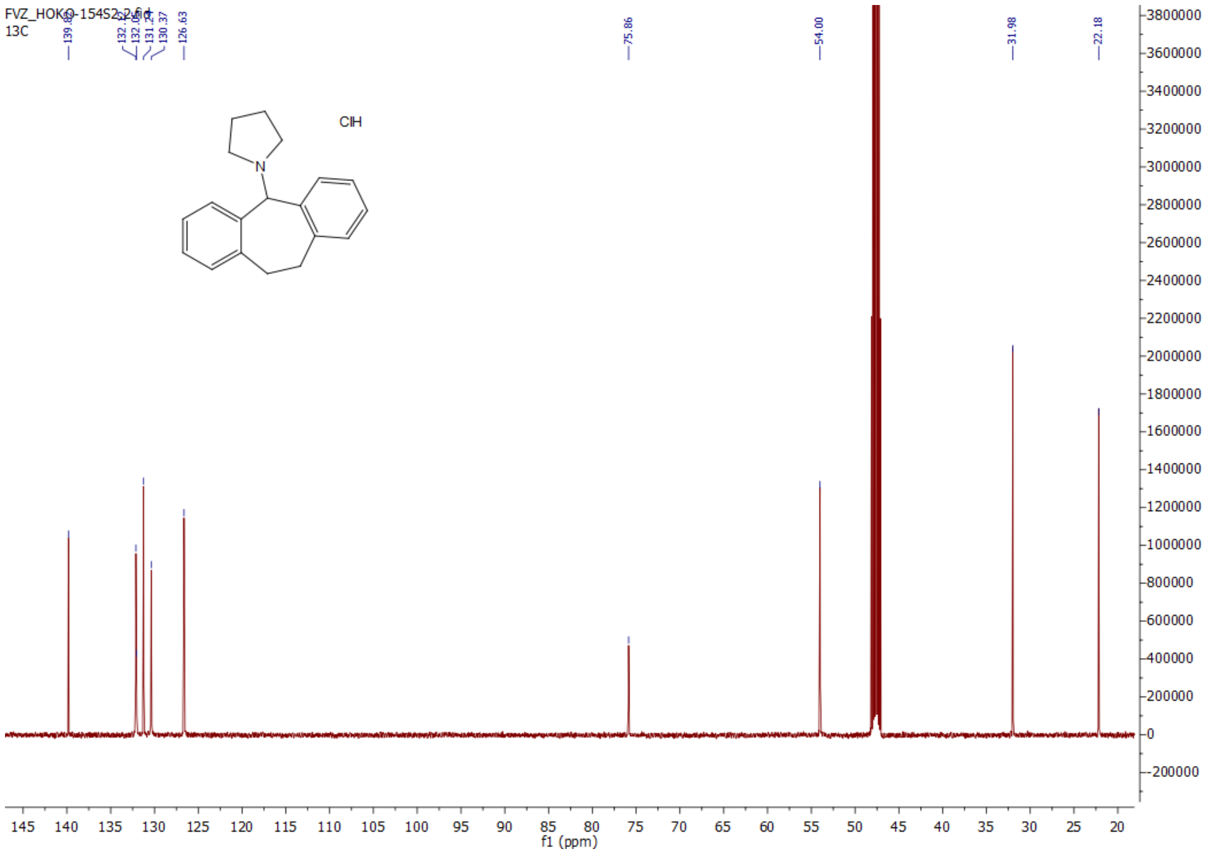


2-((10,11-dihydro-5*H*-dibenzo[*a,d*][7]annulen-5-yl)amino)ethanol hydrochloride (**3l**):

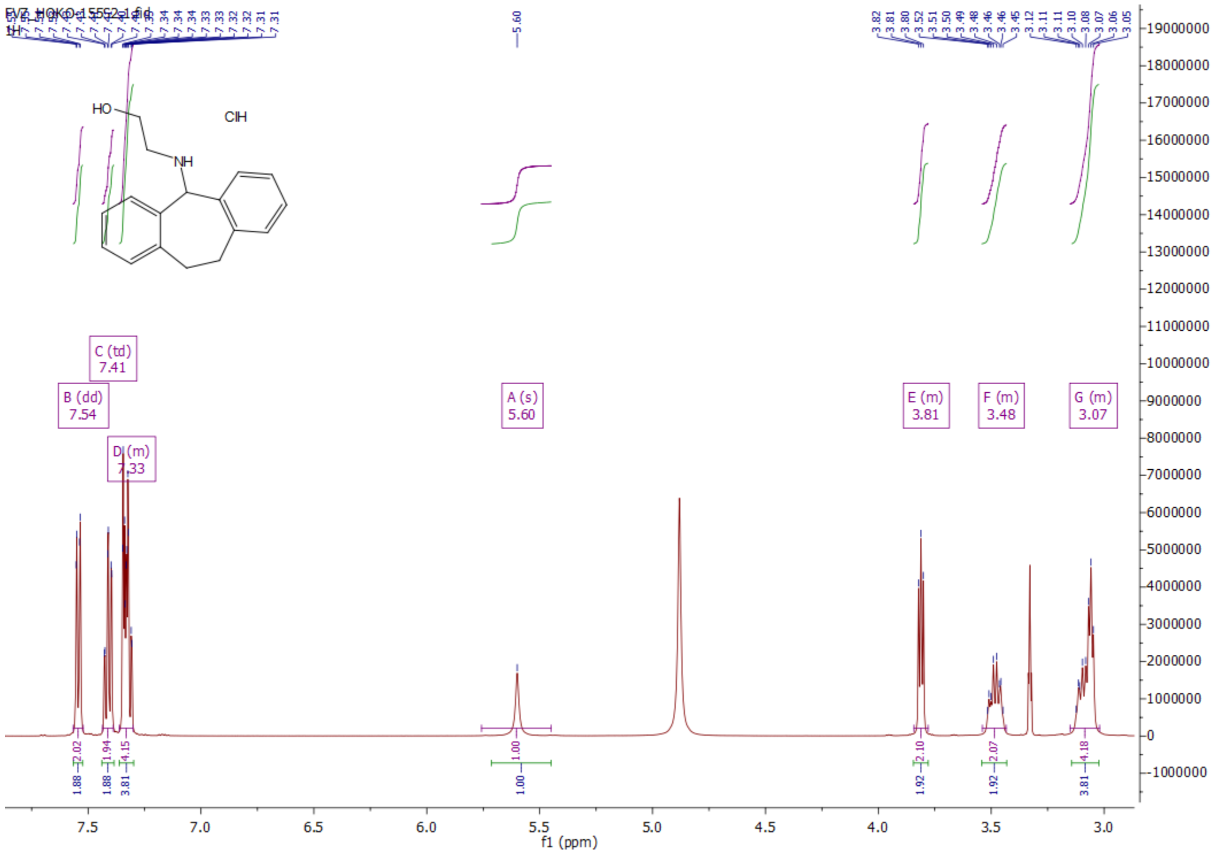

2-((10,11-dihydro-5*H*-dibenzo[*a,d*][7]annulen-5-yl)amino)ethanol hydrochloride (**3l**):

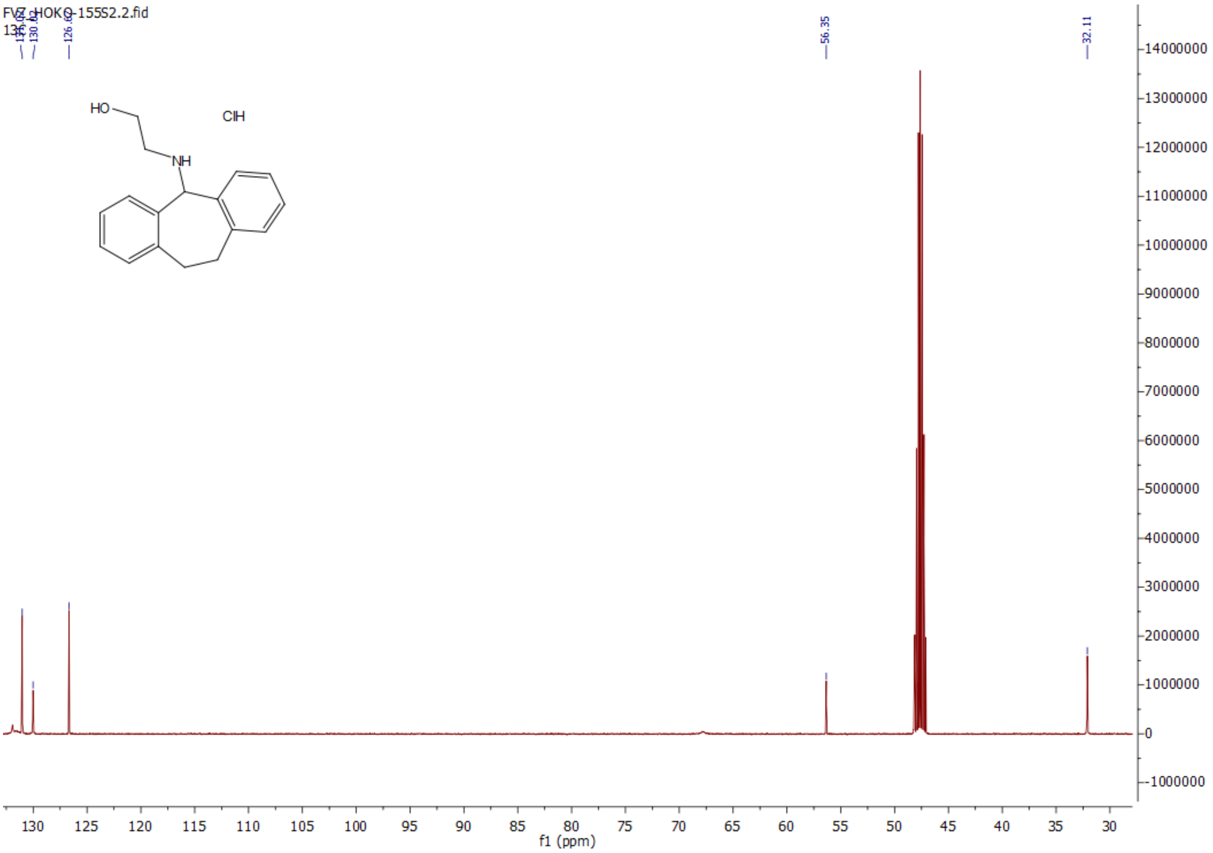


1-(10,11-dihydro-5*H*-dibenzo[*a,d*][7]annulen-5-yl)-4-methylpiperazine dihydrochloride (**3m**):

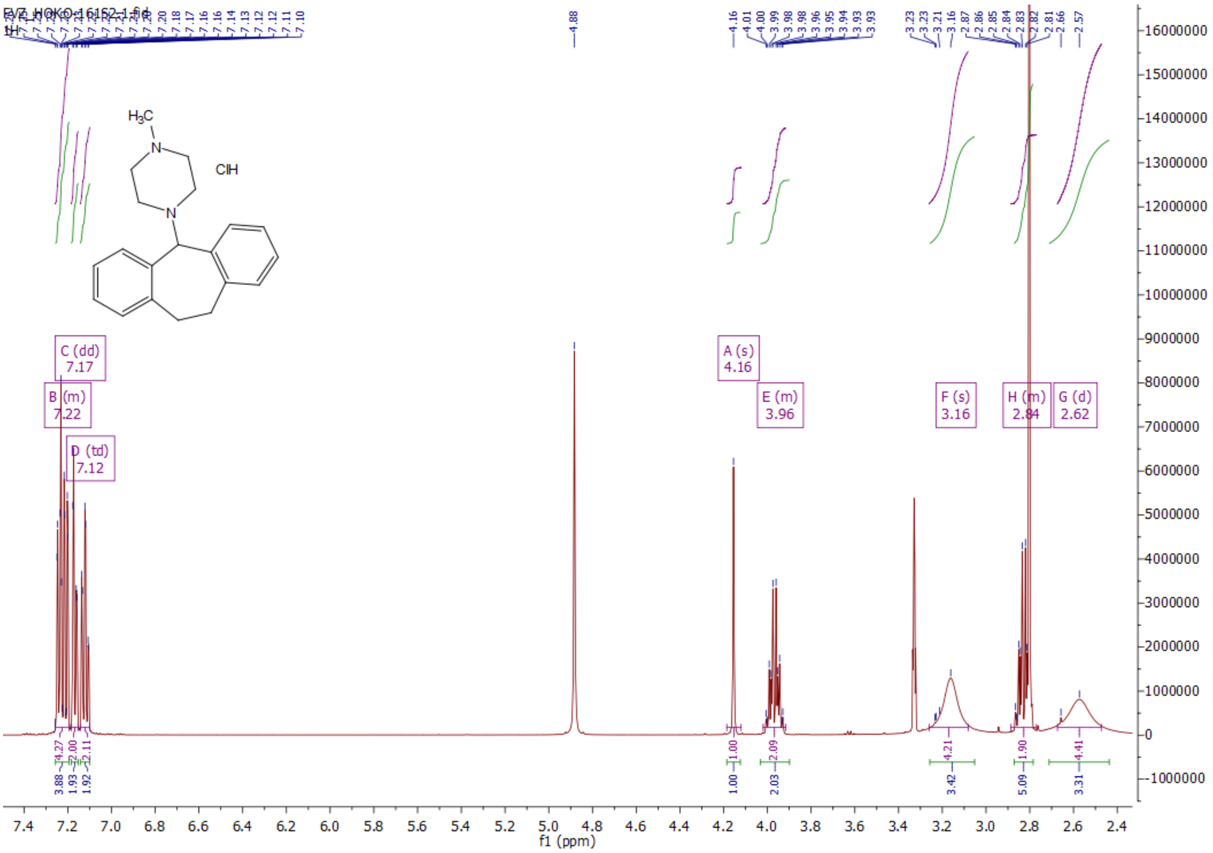

1-(10,11-dihydro-5*H*-dibenzo[*a,d*][7]annulen-5-yl)-4-methylpiperazine dihydrochloride (**3m**):

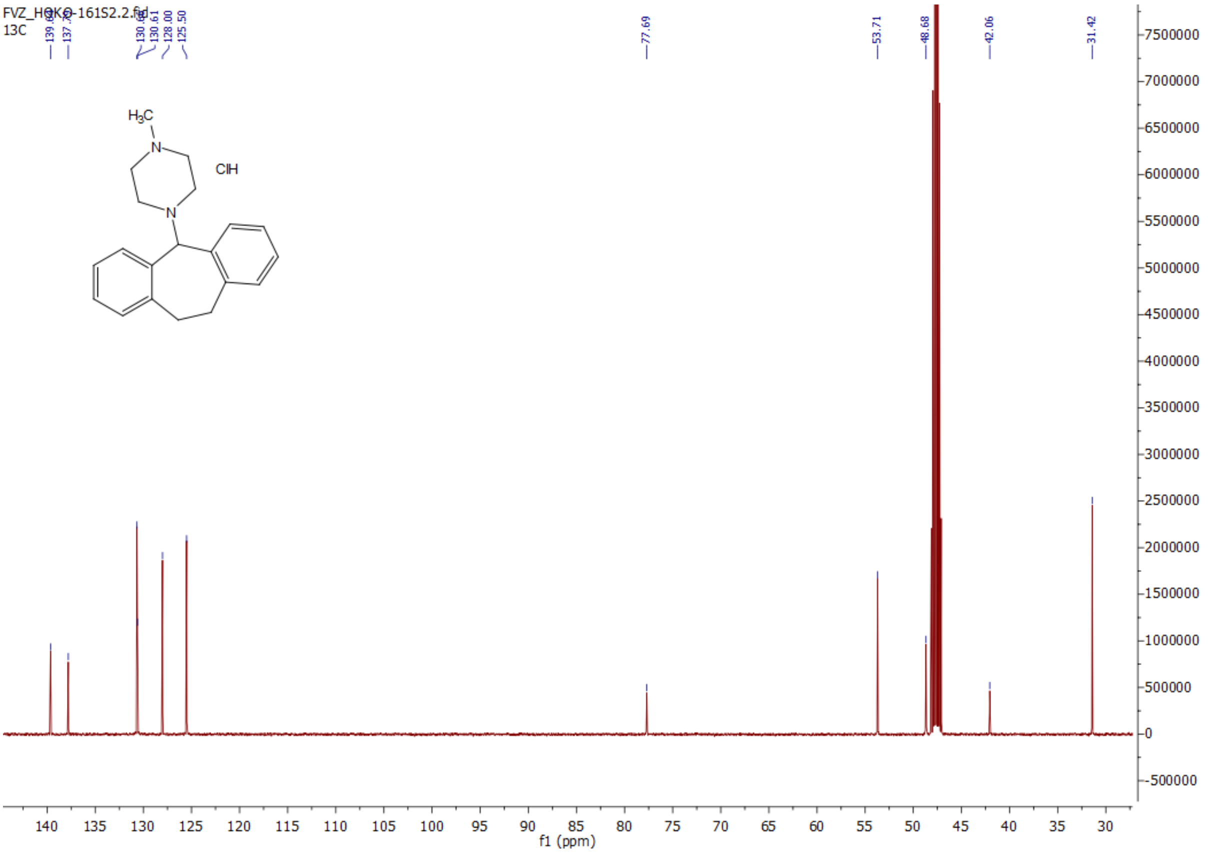


*N,N*-diethyl-10,11-dihydro-5*H*-dibenzo[*a,d*][7]annulen-5-amine hydrochloride (**3n**):

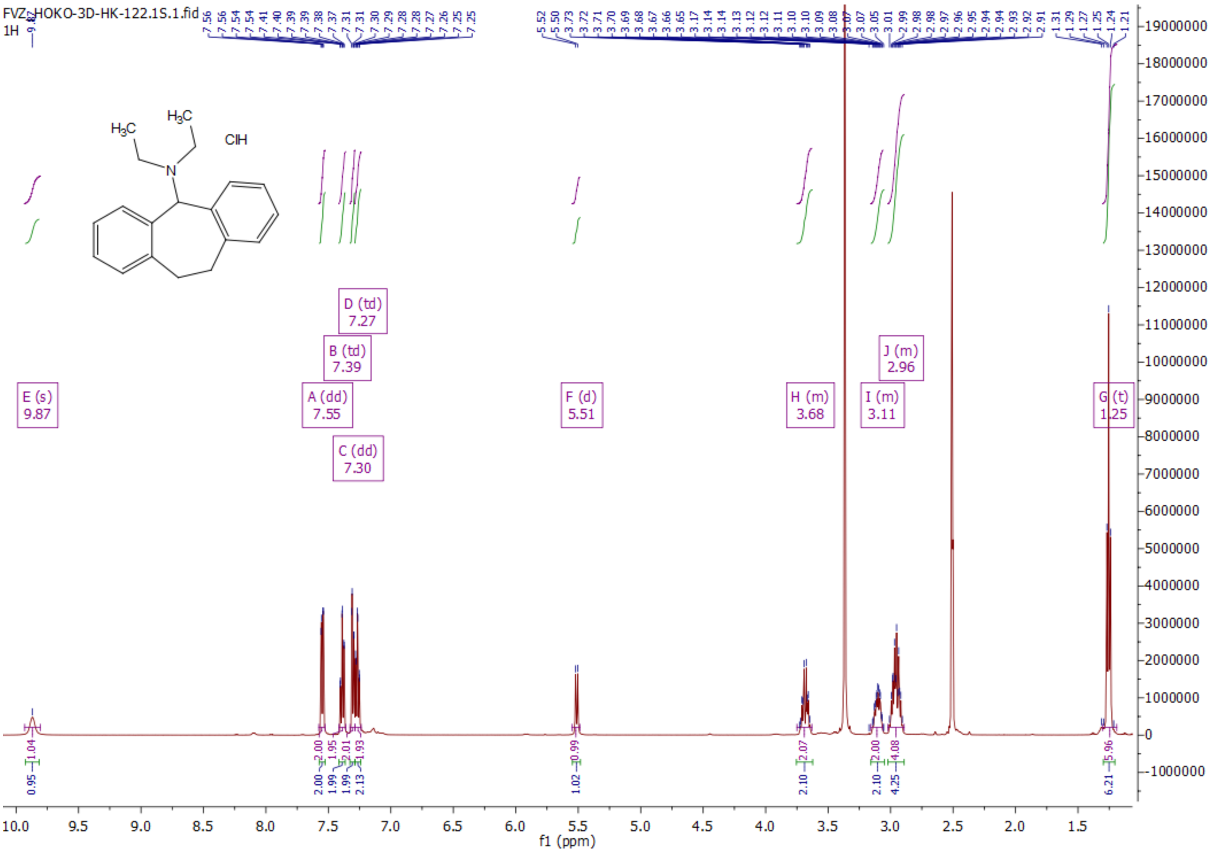

*N,N*-diethyl-10,11-dihydro-5*H*-dibenzo[*a,d*][7]annulen-5-amine hydrochloride (**3n**):

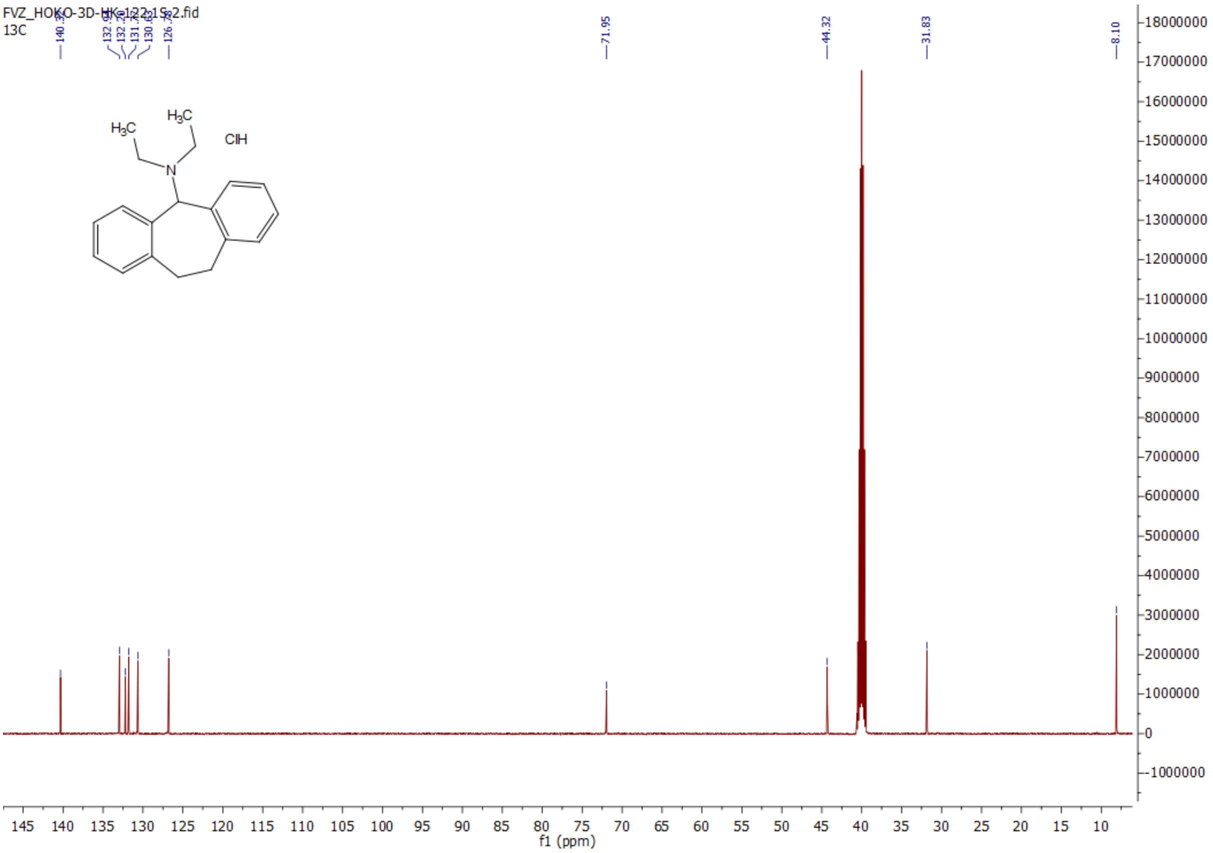


1-(10,11-dihydro-5*H*-dibenzo[*a,d*][7]annulen-5-yl)-4-ethylpiperazine dihydrochloride (**3o**):

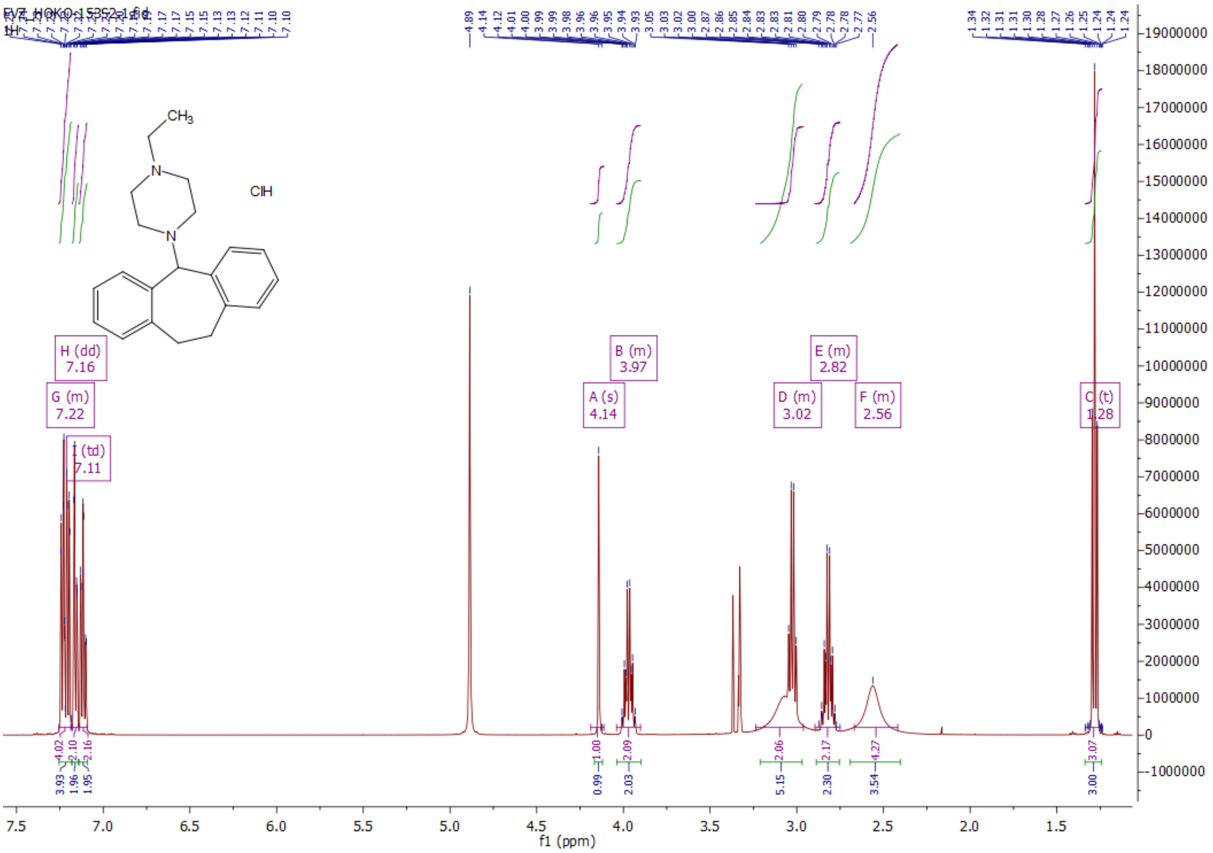

1-(10,11-dihydro-5*H*-dibenzo[*a,d*][7]annulen-5-yl)-4-ethylpiperazine dihydrochloride (**3o**):

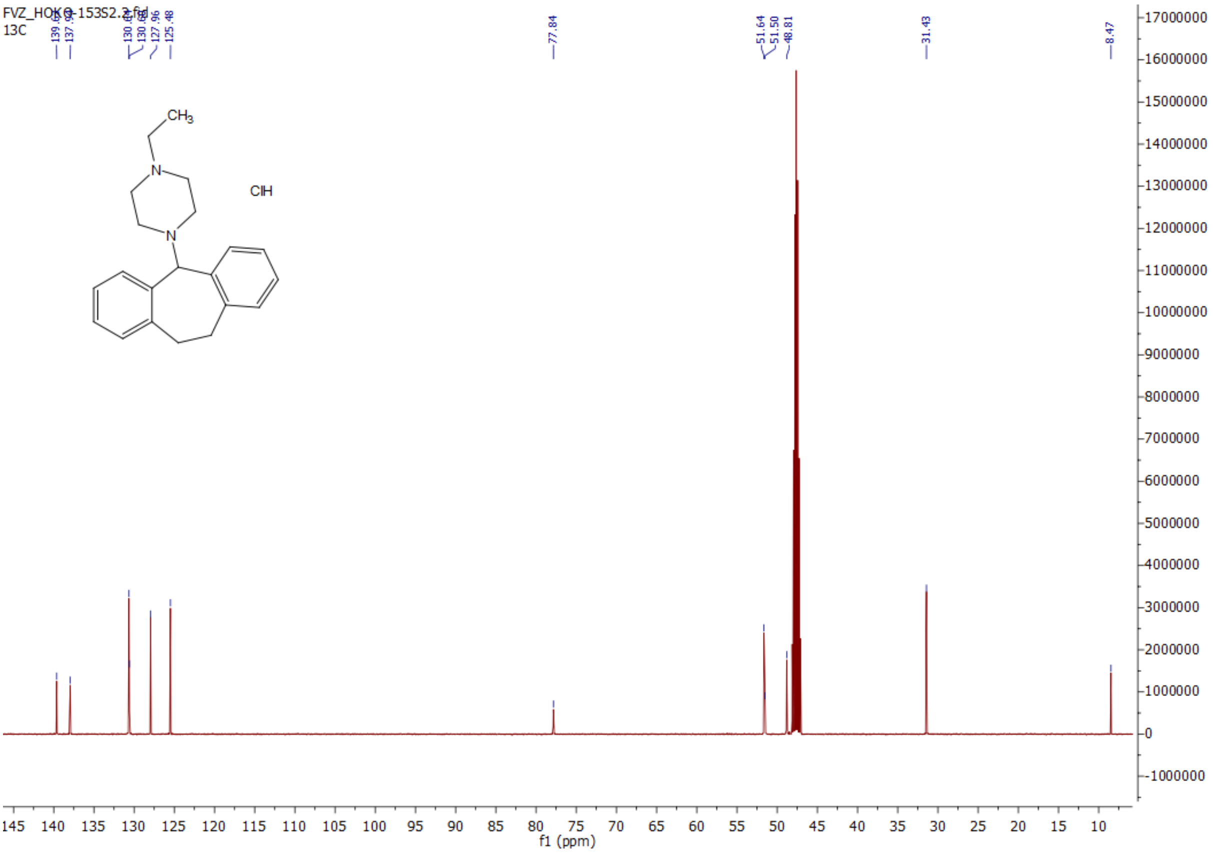

*N*-methyl-5*H*-dibenzo[*a,d*][7]annulen-5-amine hydrochloride (**6a**)

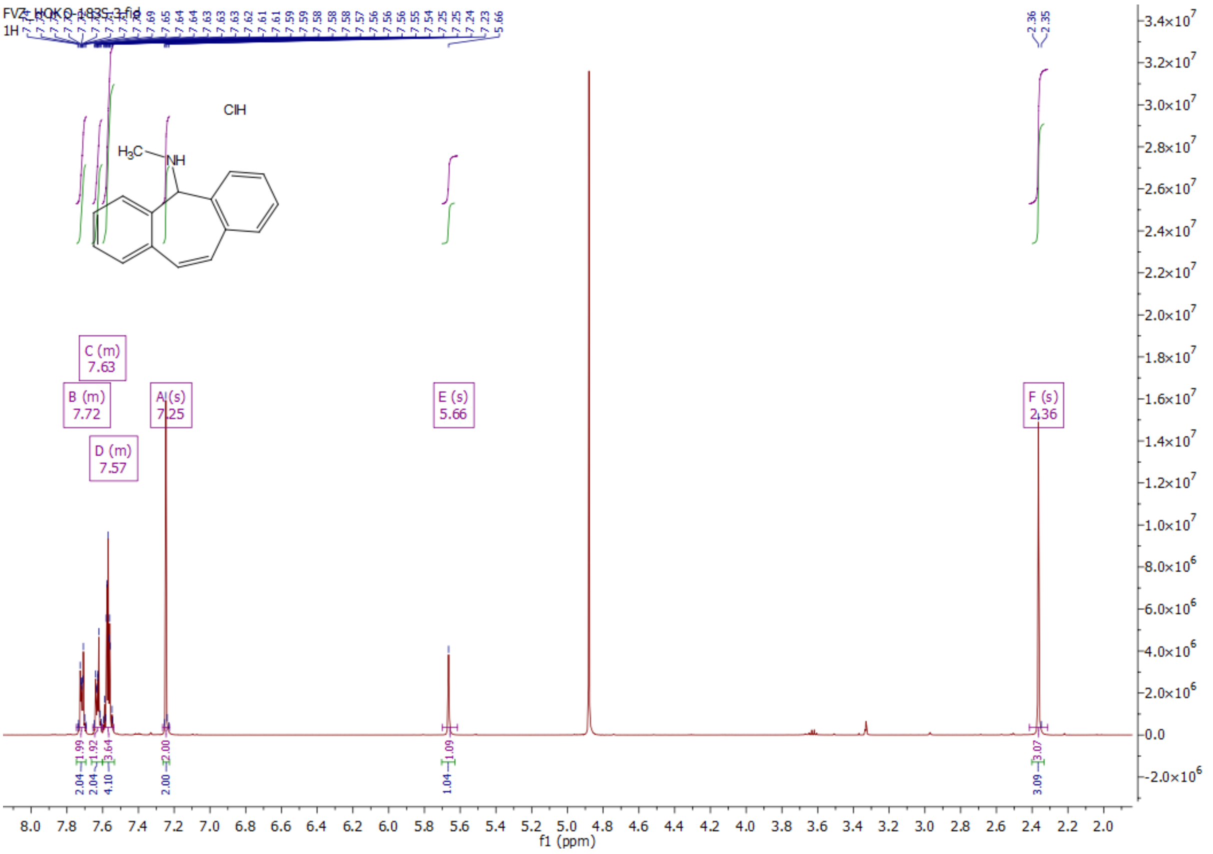

*N*-methyl-5*H*-dibenzo[*a,d*][7]annulen-5-amine hydrochloride (**6a**)

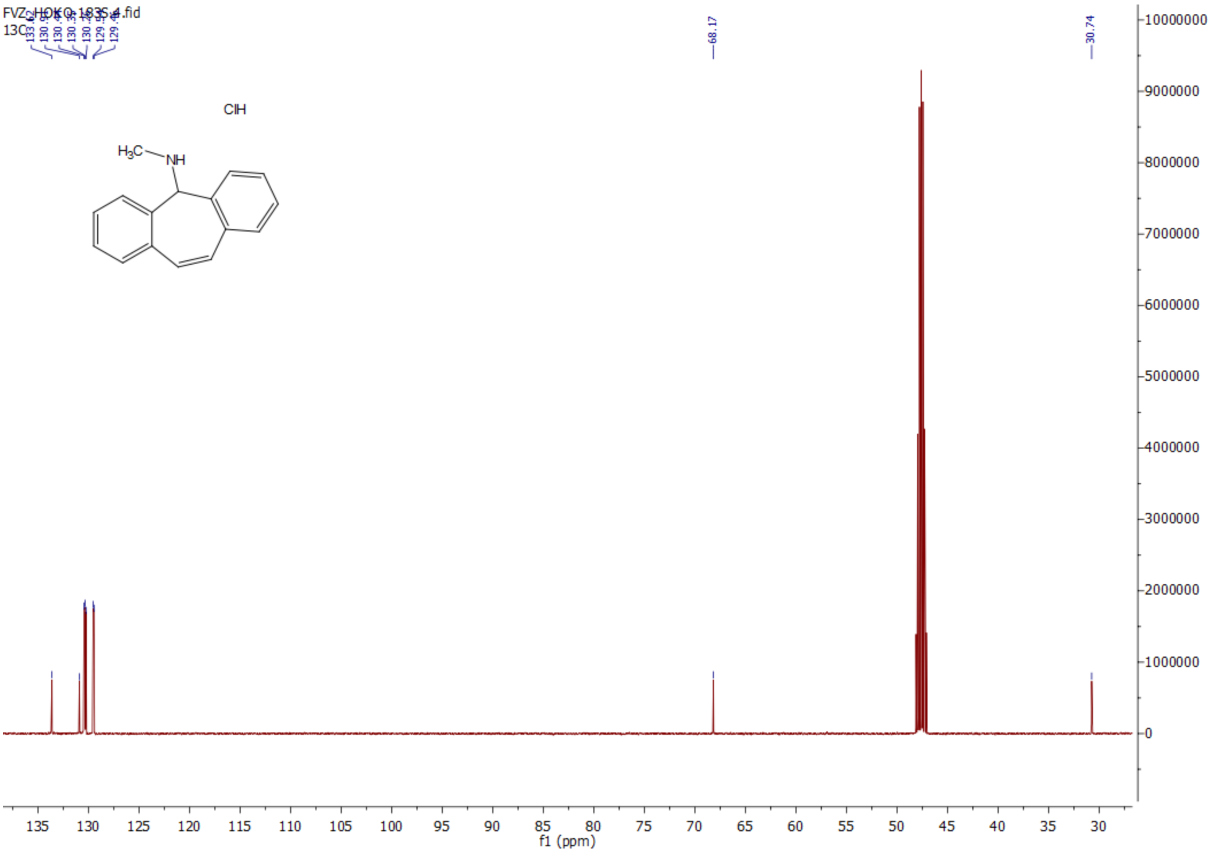

*N*-ethyl-5*H*-dibenzo[*a,d*][7]annulen-5-amine hydrochloride (**6b**)

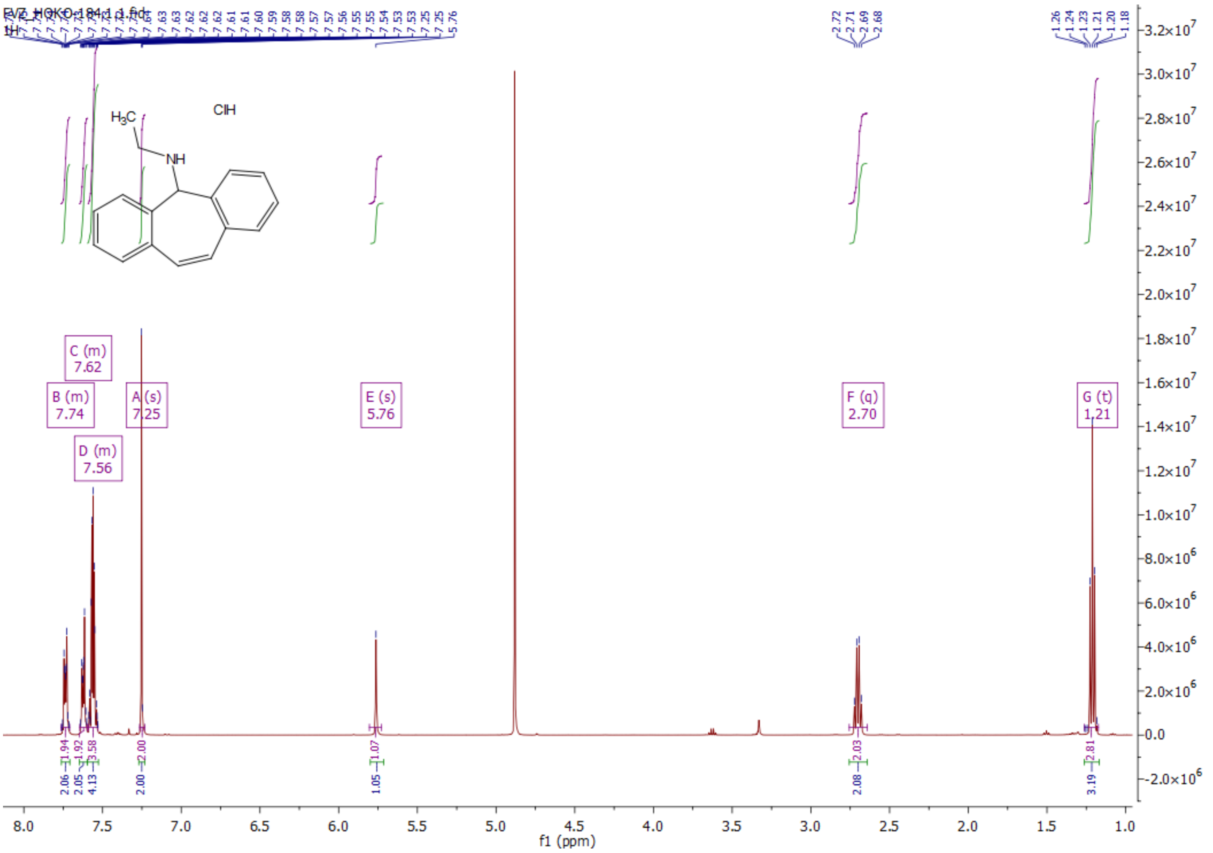

*N*-ethyl-5*H*-dibenzo[*a,d*][7]annulen-5-amine hydrochloride (**6b**)

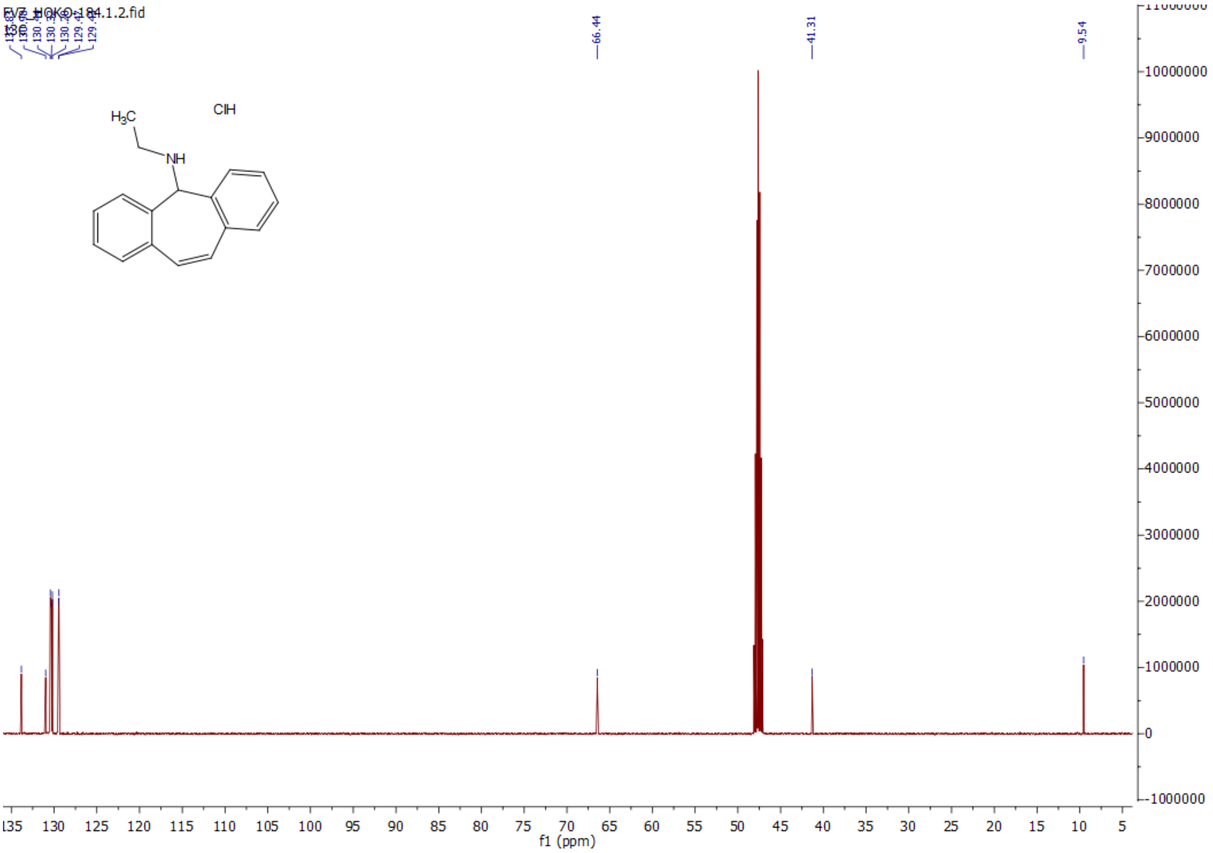

*N*-isopropyl-5*H*-dibenzo[*a,d*][7]annulen-5-amine hydrochloride (**6c**)

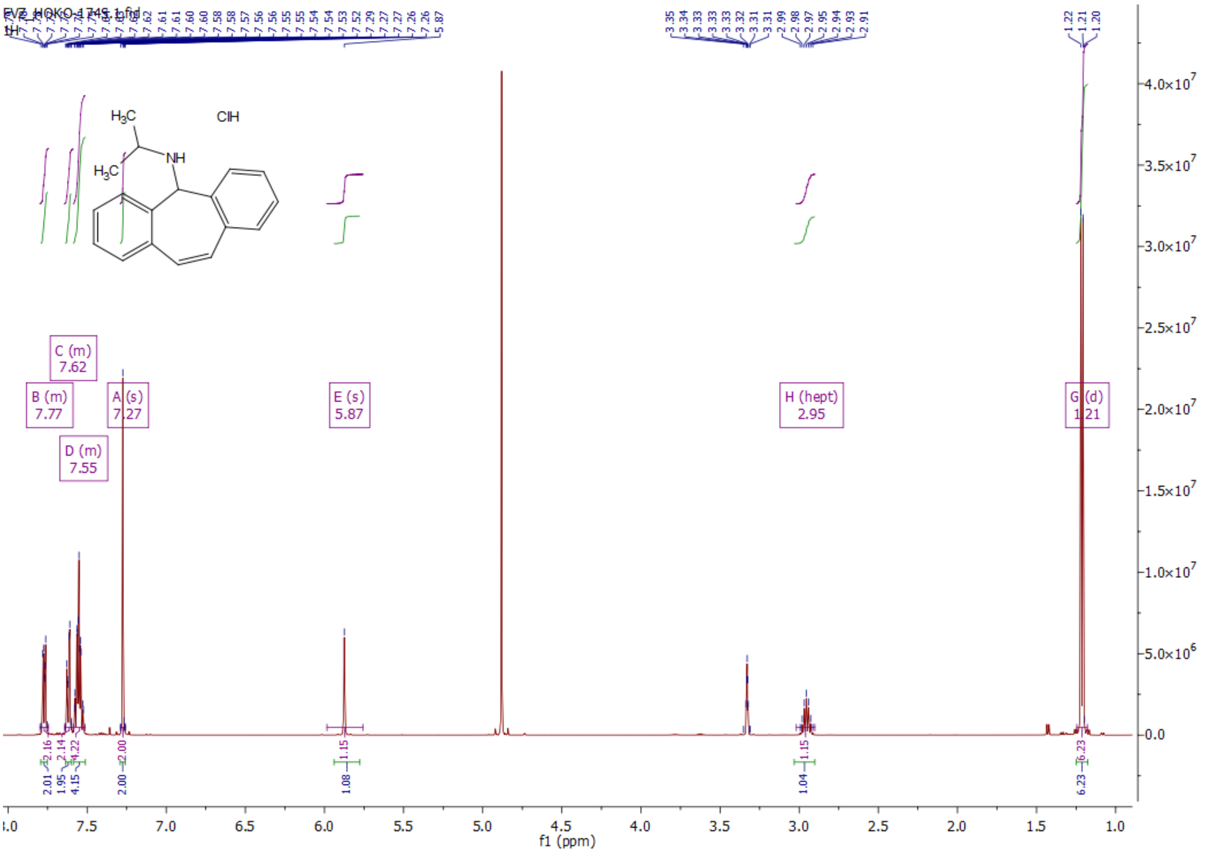

*N*-isopropyl-5*H*-dibenzo[*a,d*][7]annulen-5-amine hydrochloride (**6c**)

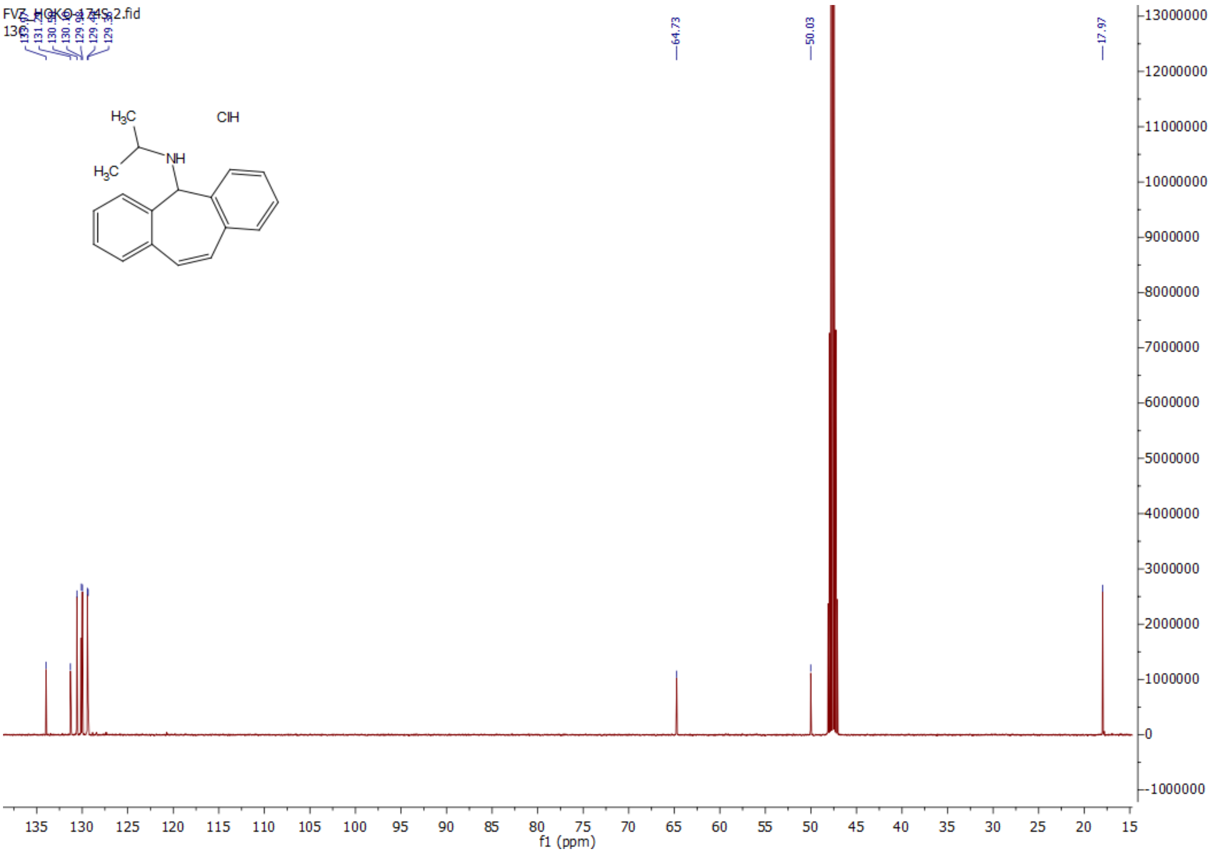

*N*-cyclopropyl-5*H*-dibenzo[*a,d*][7]annulen-5-amine hydrochloride (**6d**)

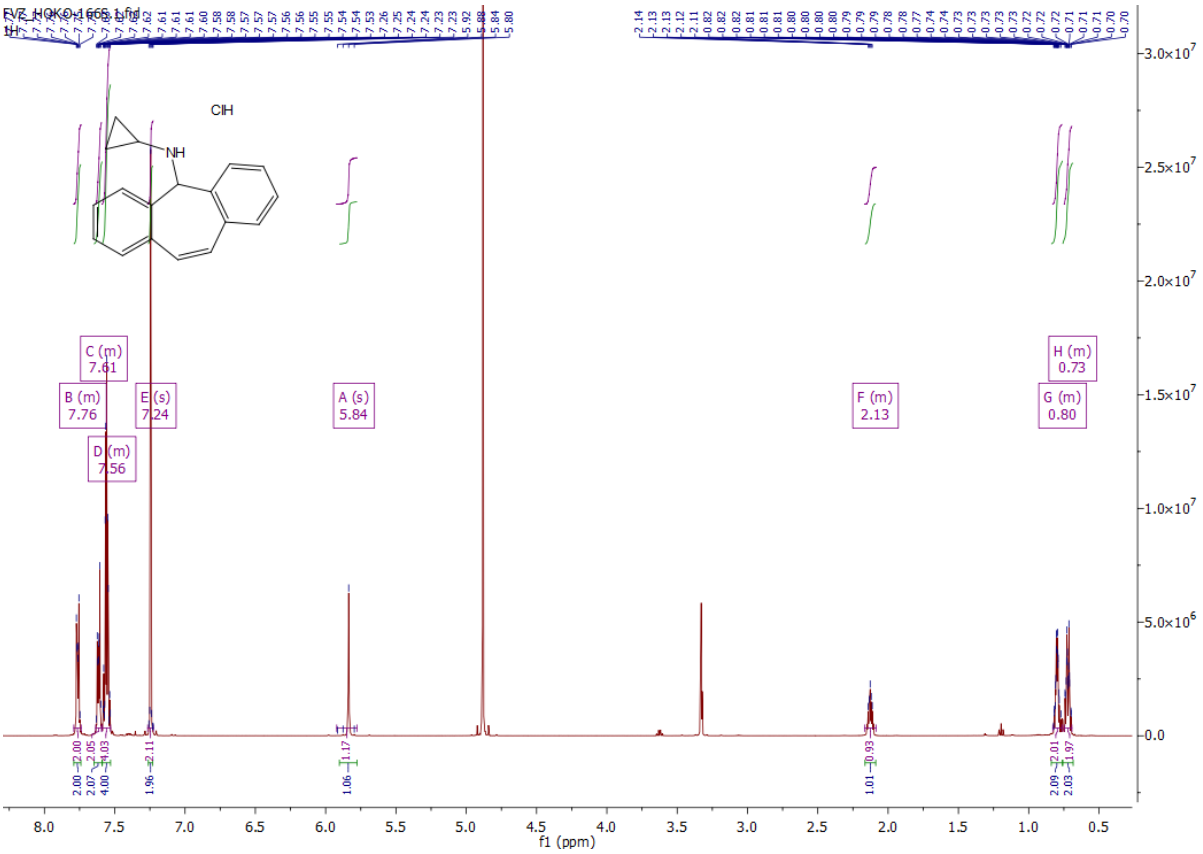

*N*-cyclopropyl-5*H*-dibenzo[*a,d*][7]annulen-5-amine hydrochloride (**6d**)

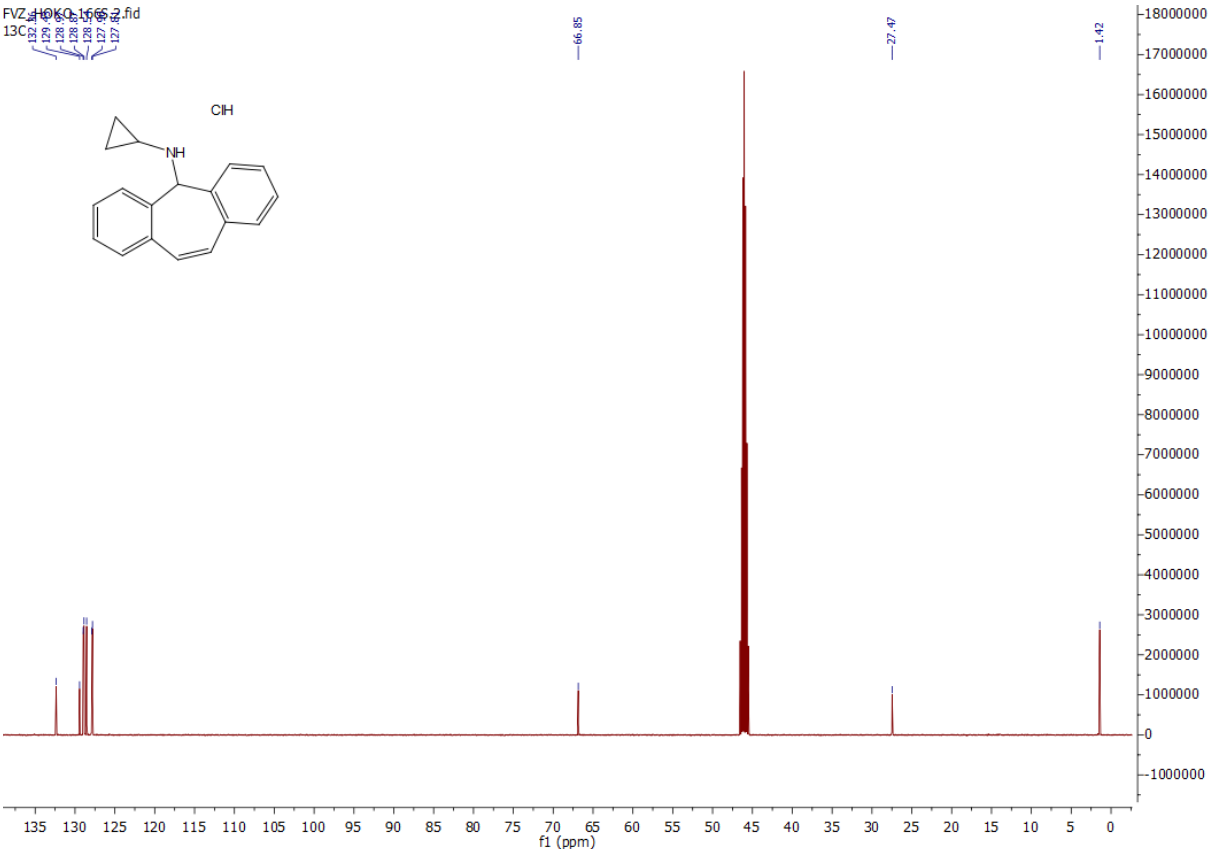

*N*-butyl-5*H*-dibenzo[*a,d*][7]annulen-5-amine hydrochloride (**6e**)

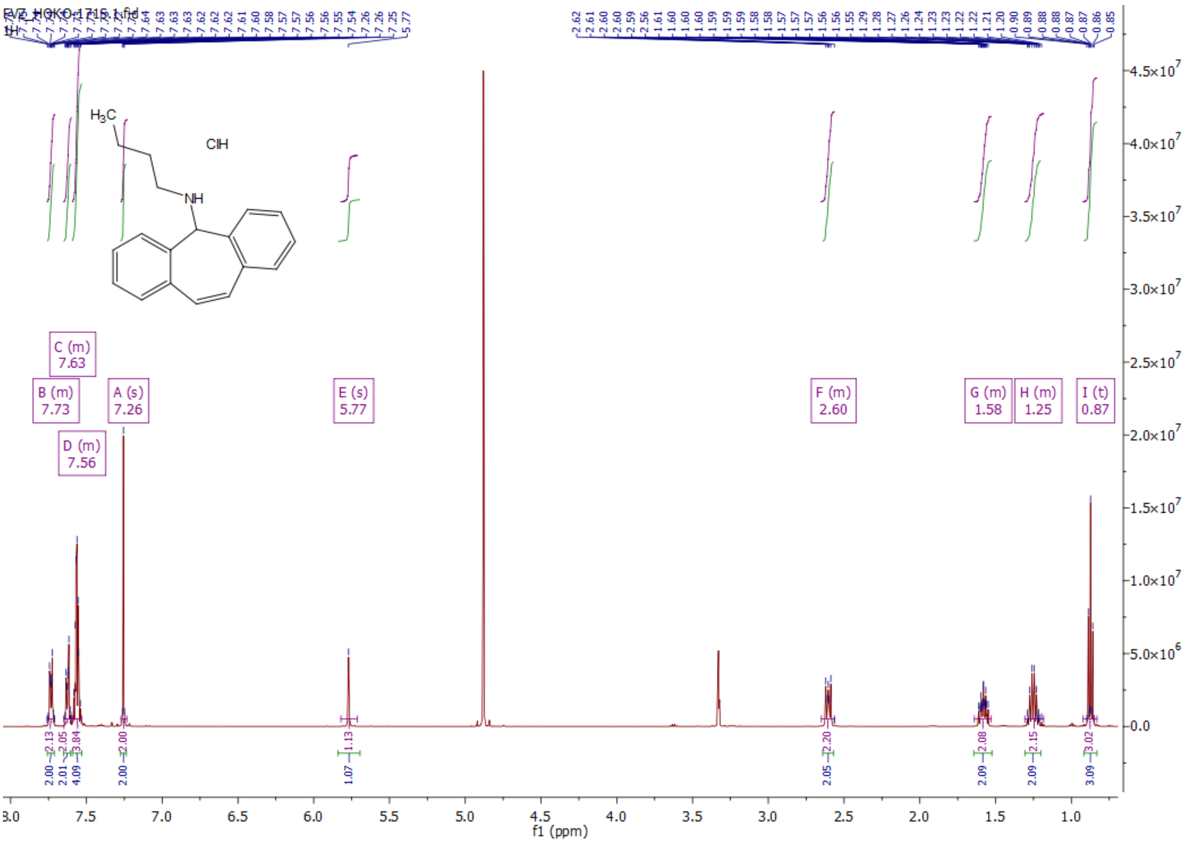


*N*-butyl-5*H*-dibenzo[*a,d*][7]annulen-5-amine hydrochloride (**6e**)

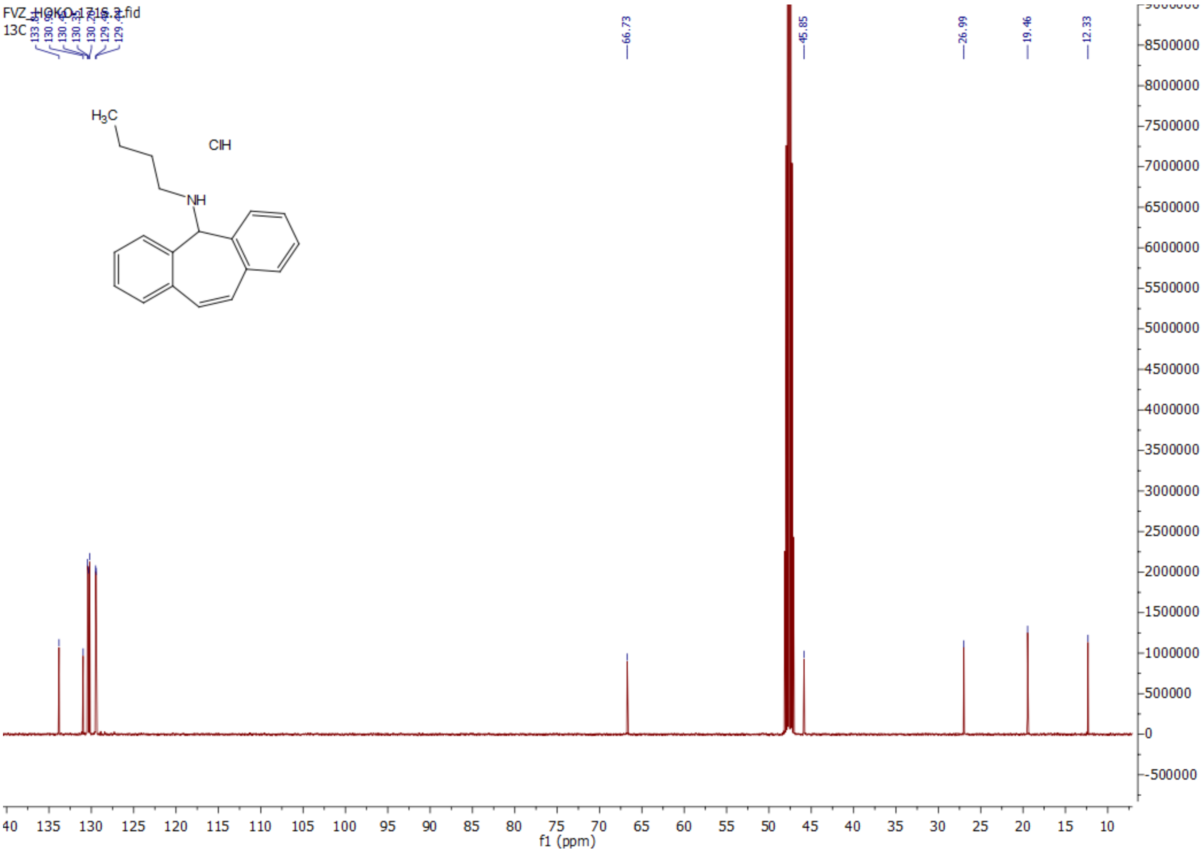

*N*-isobutyl-5*H*-dibenzo[*a,d*][7]annulen-5-amine hydrochloride (**6f**)

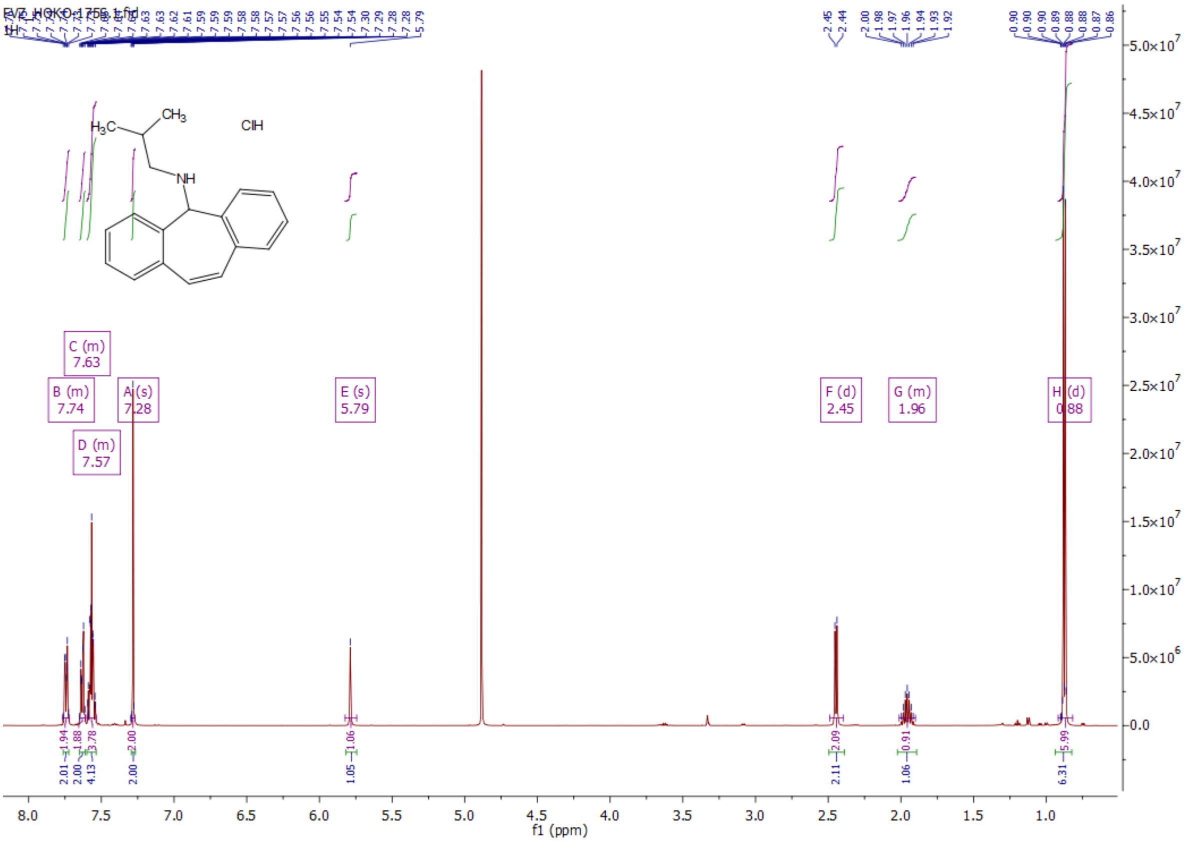


*N*-isobutyl-5*H*-dibenzo[*a,d*][7]annulen-5-amine hydrochloride (**6f**)

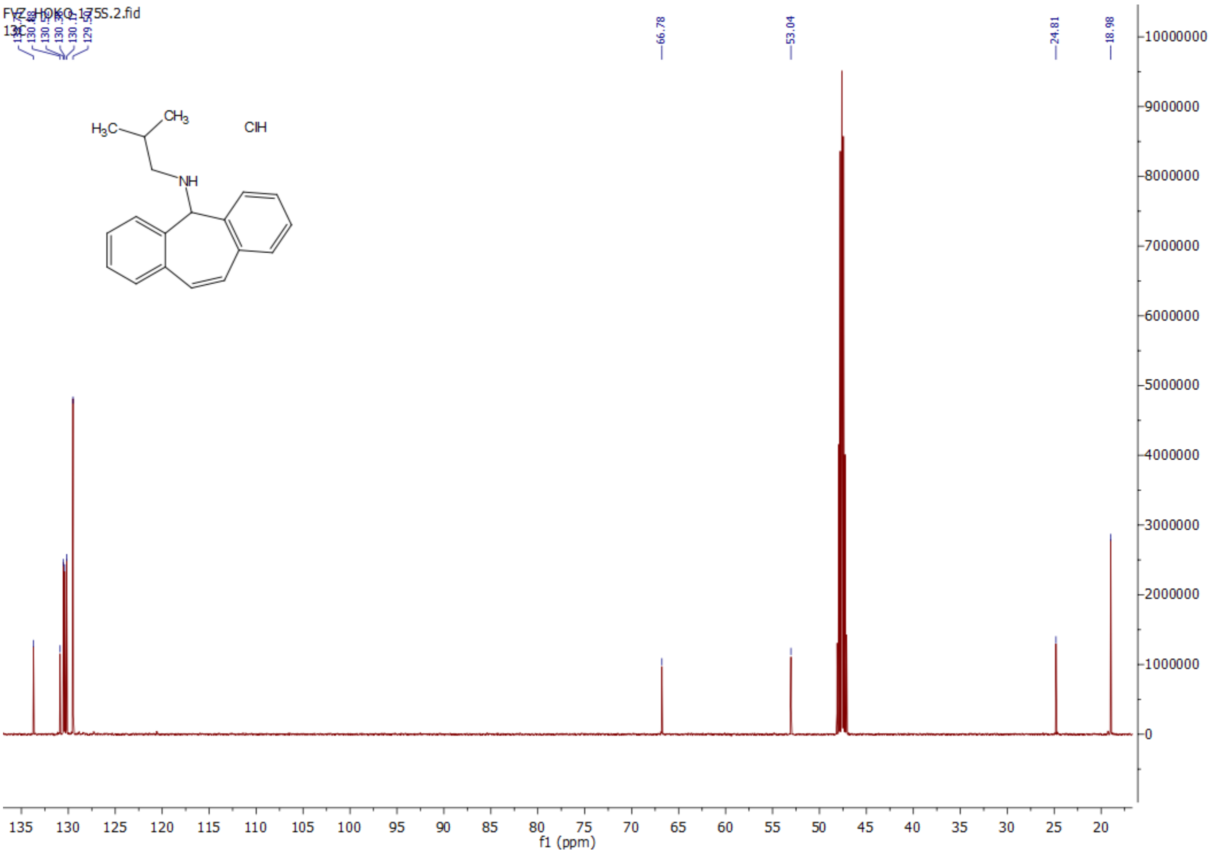

*N*-cyclohexyl-5*H*-dibenzo[*a,d*][7]annulen-5-amine hydrochloride (**6g**)

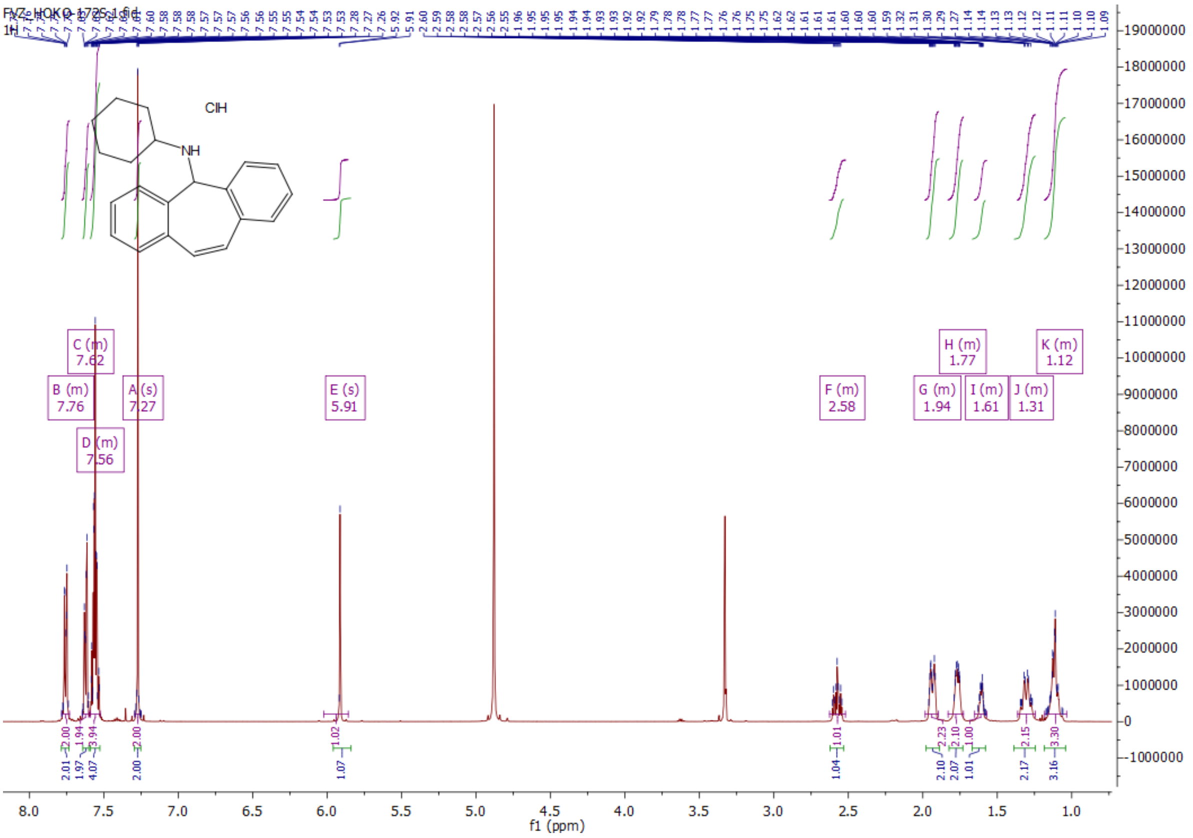

*N*-cyclohexyl-5*H*-dibenzo[*a,d*][7]annulen-5-amine hydrochloride (**6g**)

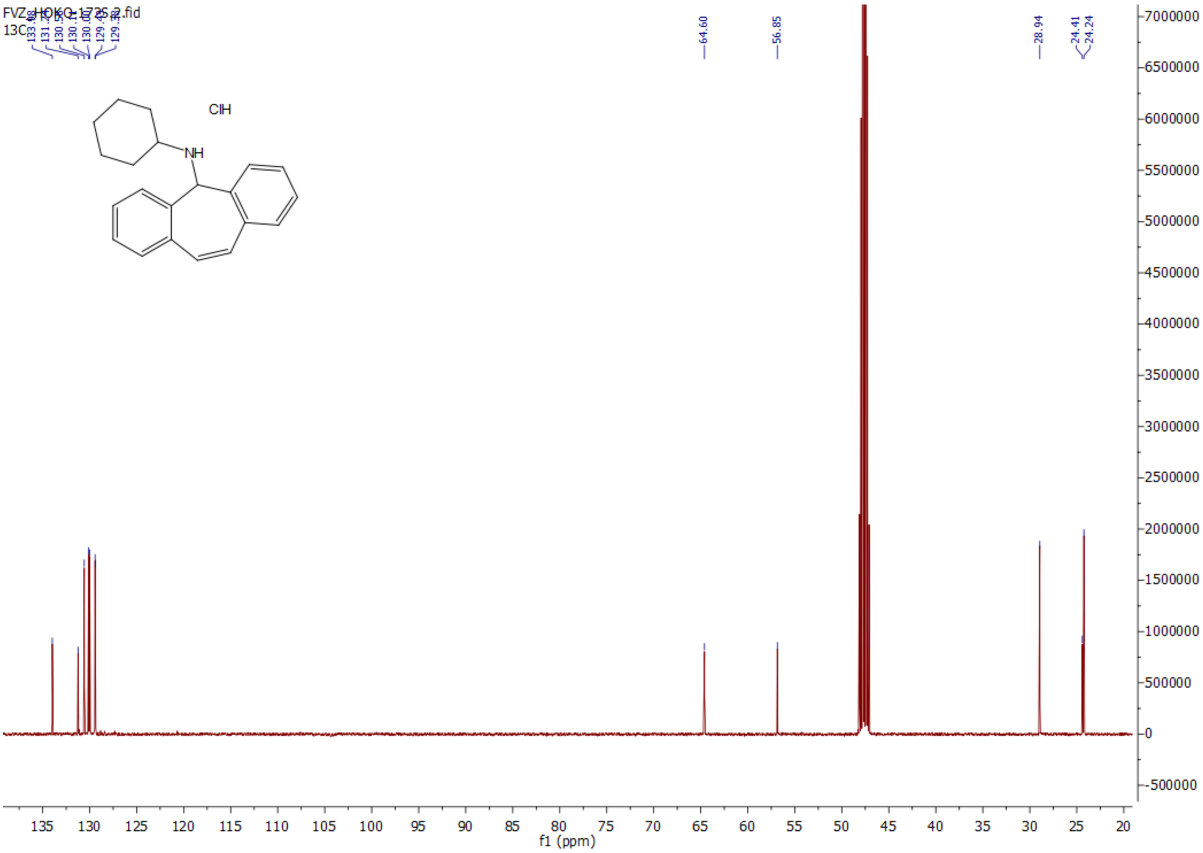

1-(5*H*-dibenzo[*a*,*d*][7]annulen-5-yl)piperidine hydrochloride (**6h**)

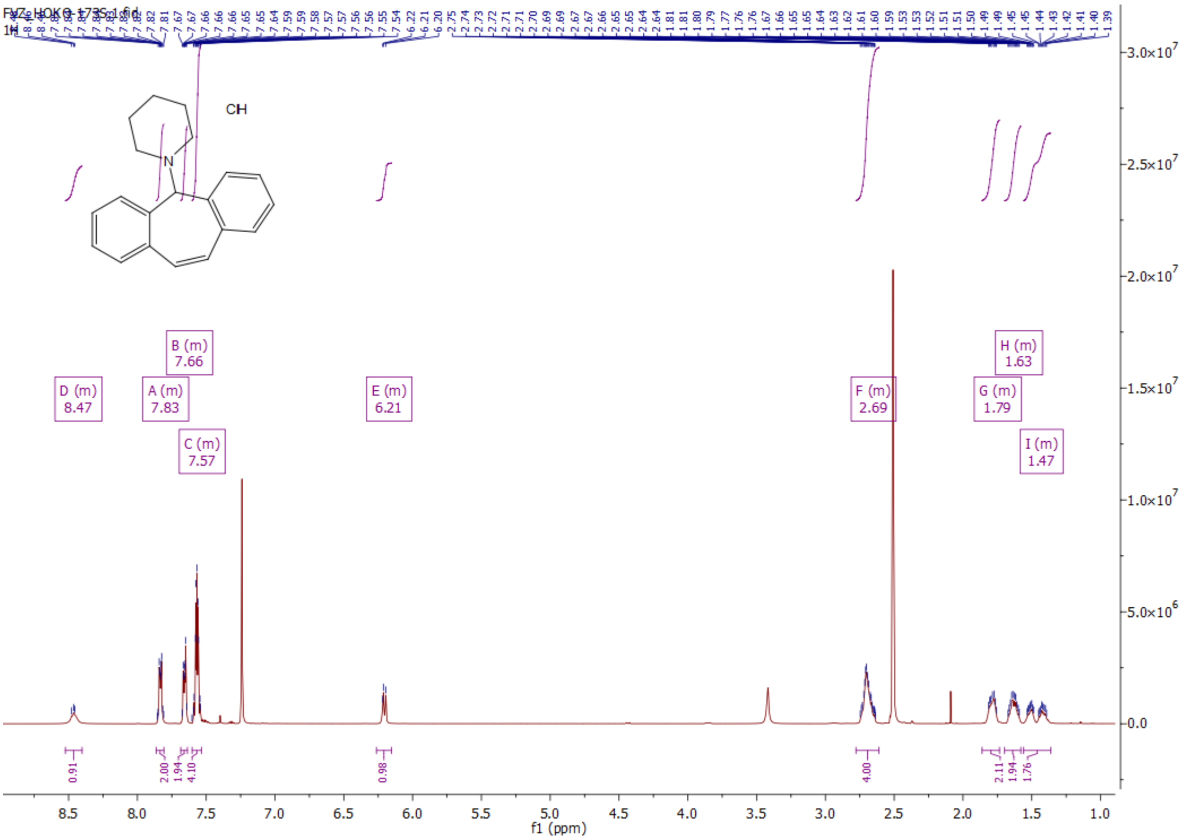


1-(5*H*-dibenzo[*a*,*d*][7]annulen-5-yl)piperidine hydrochloride (**6h**)

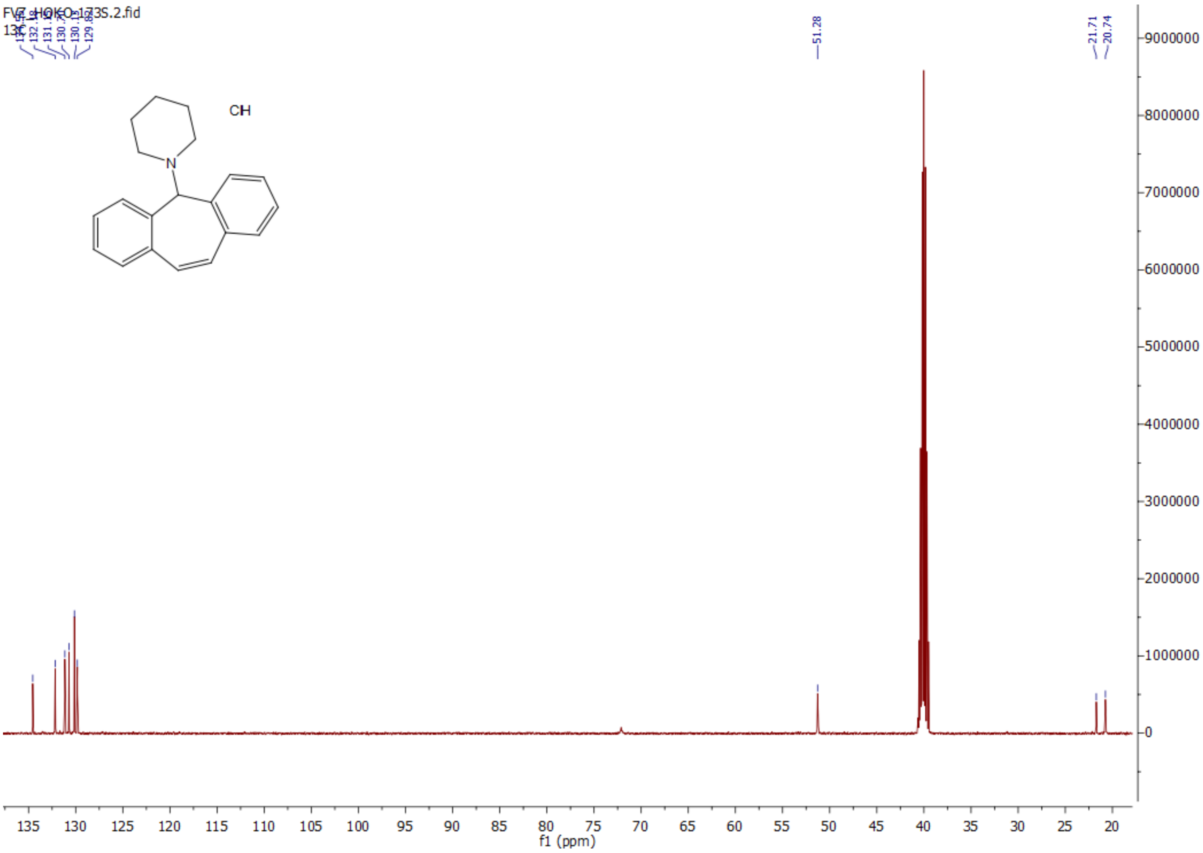

4-(5*H*-dibenzo[*a*,*d*][7]annulen-5-yl)morpholine hydrochloride (**6i**)

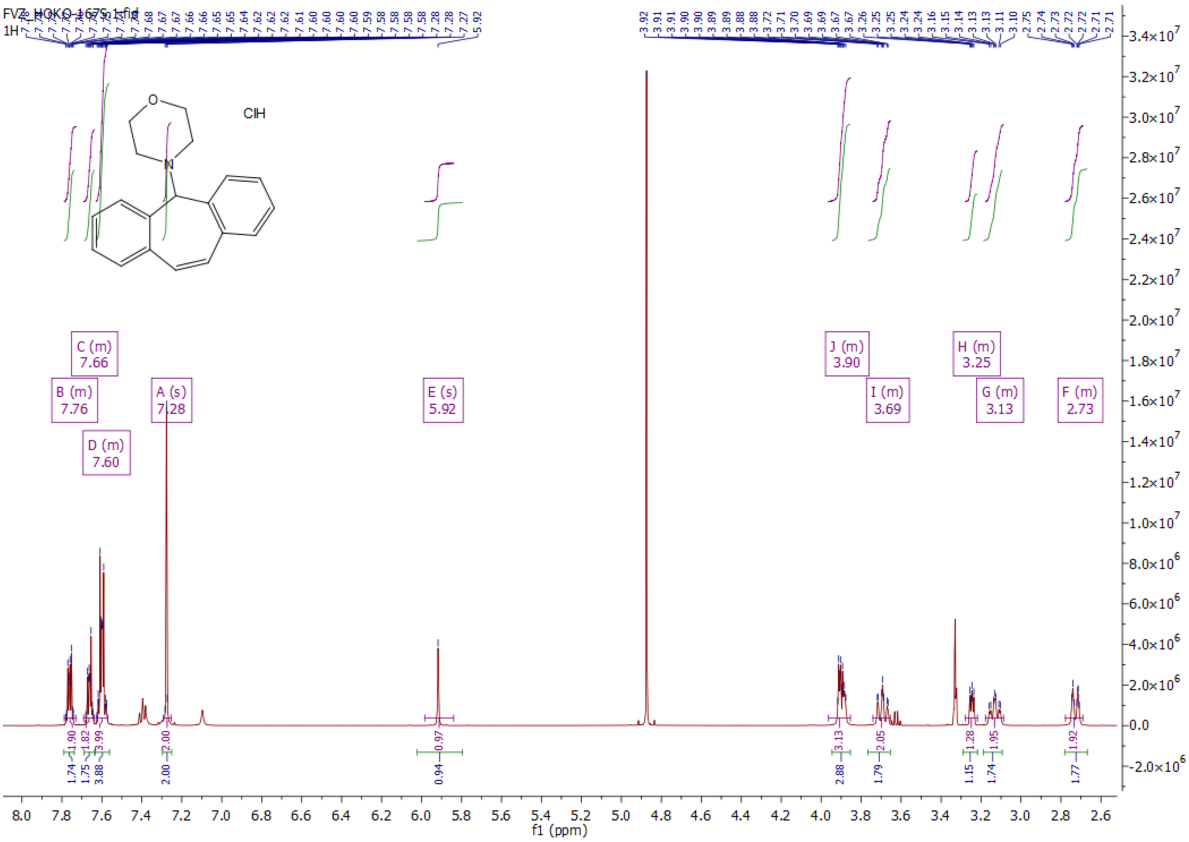

4-(5*H*-dibenzo[*a*,*d*][7]annulen-5-yl)morpholine hydrochloride (**6i**)

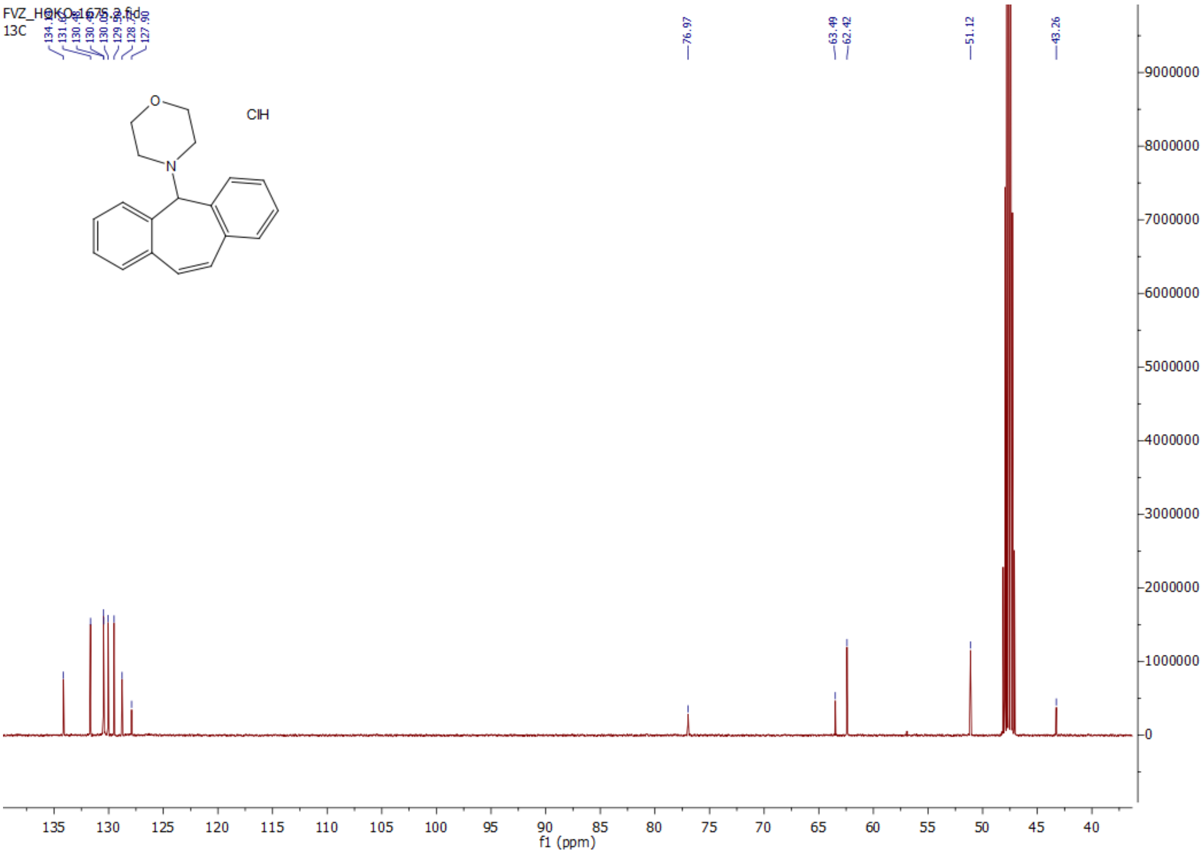

*N*-(2-methoxyethyl)-5*H*-dibenzo[*a,d*][7]annulen-5-amine hydrochloride (**6j**)

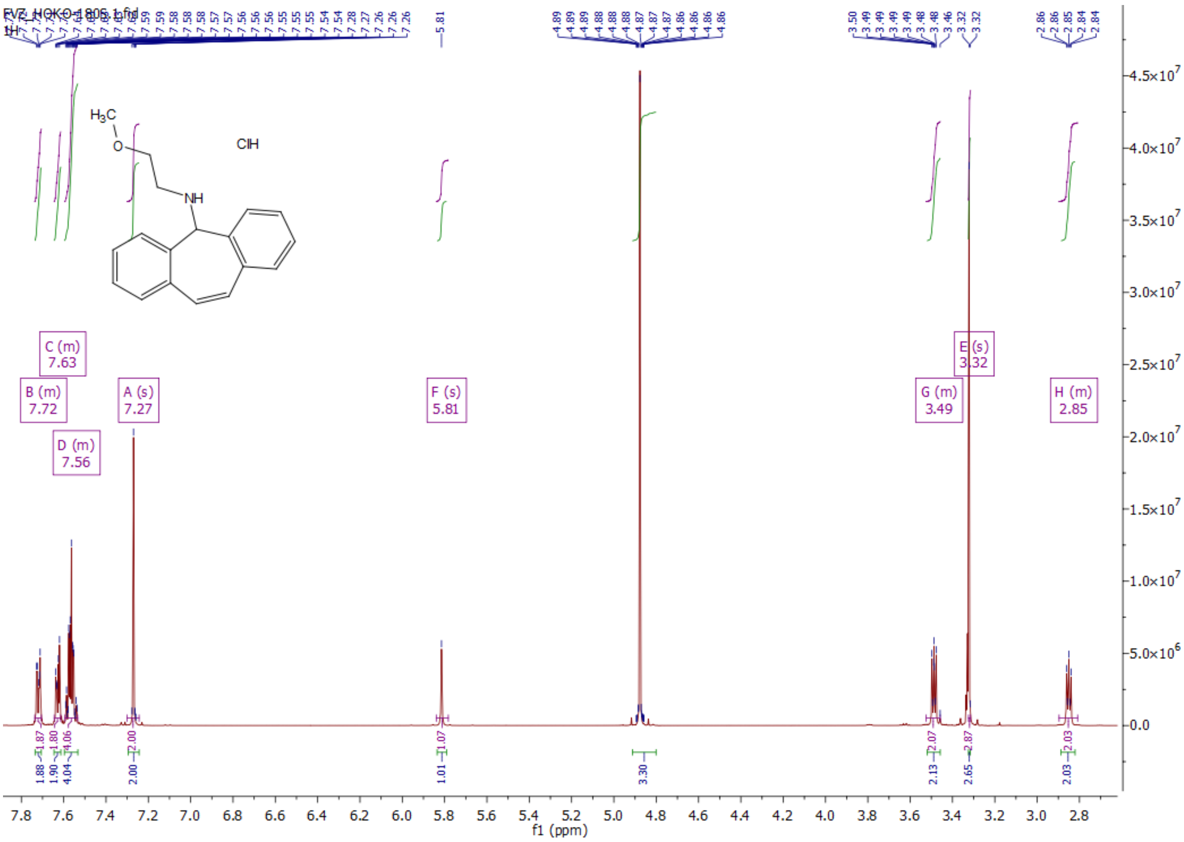


*N*-(2-methoxyethyl)-5*H*-dibenzo[*a,d*][7]annulen-5-amine hydrochloride (**6j**)

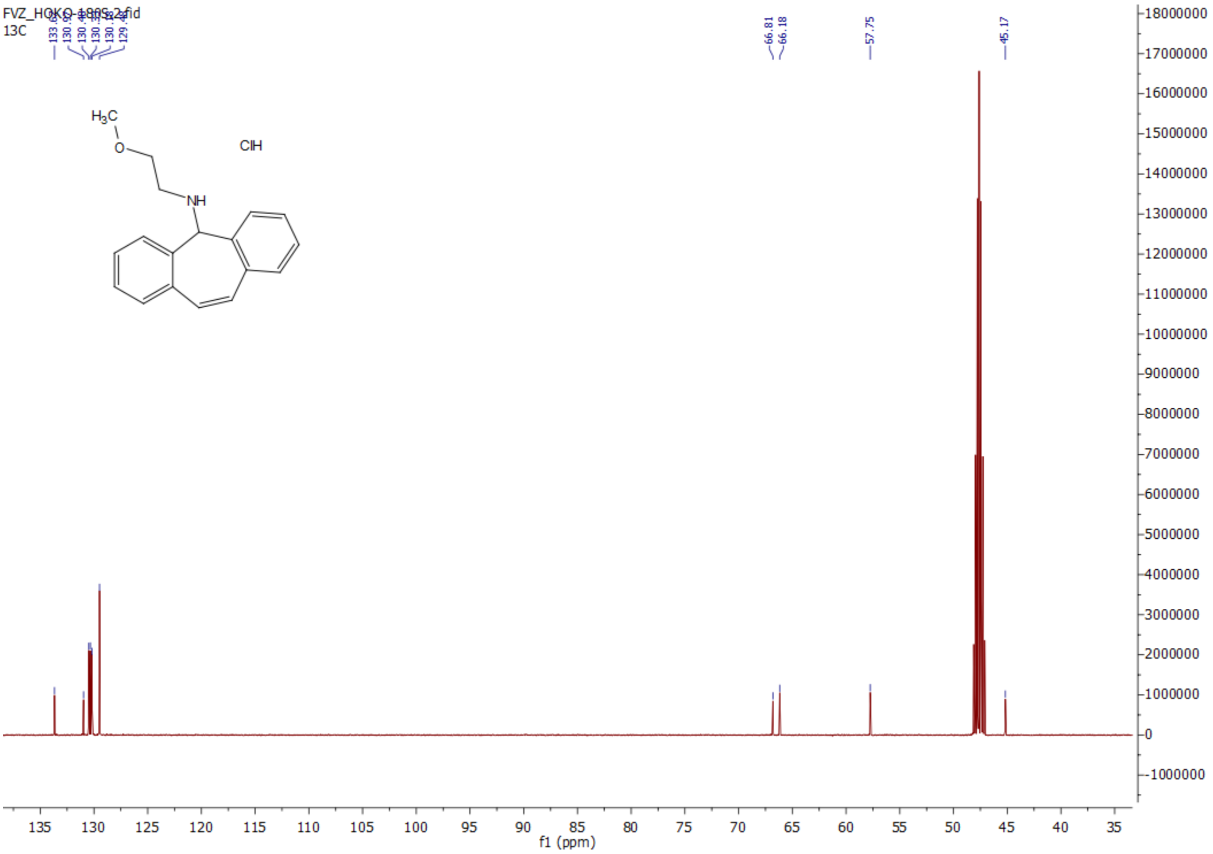

1-(5*H*-dibenzo[*a,d*][7]annulen-5-yl)pyrrolidine hydrochloride (**6k**)

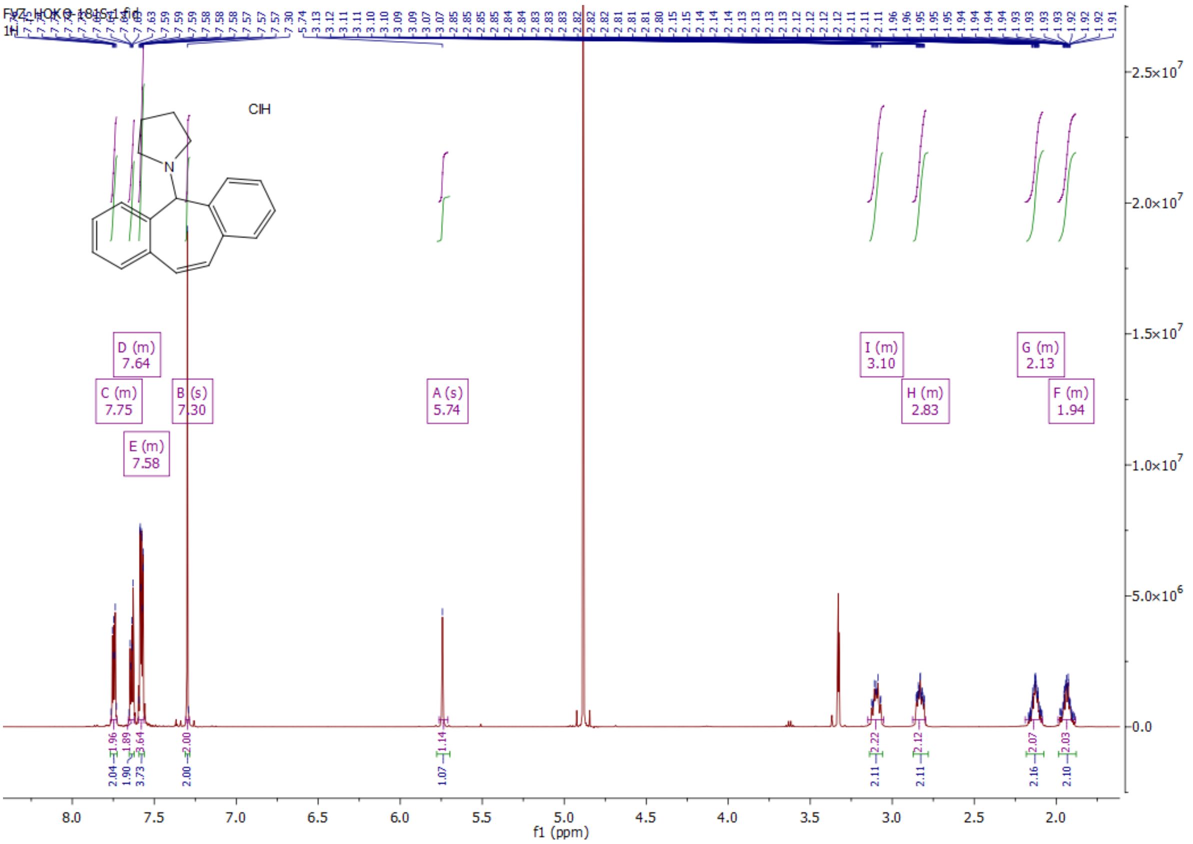

1-(5*H*-dibenzo[*a,d*][7]annulen-5-yl)pyrrolidine hydrochloride (**6k**)

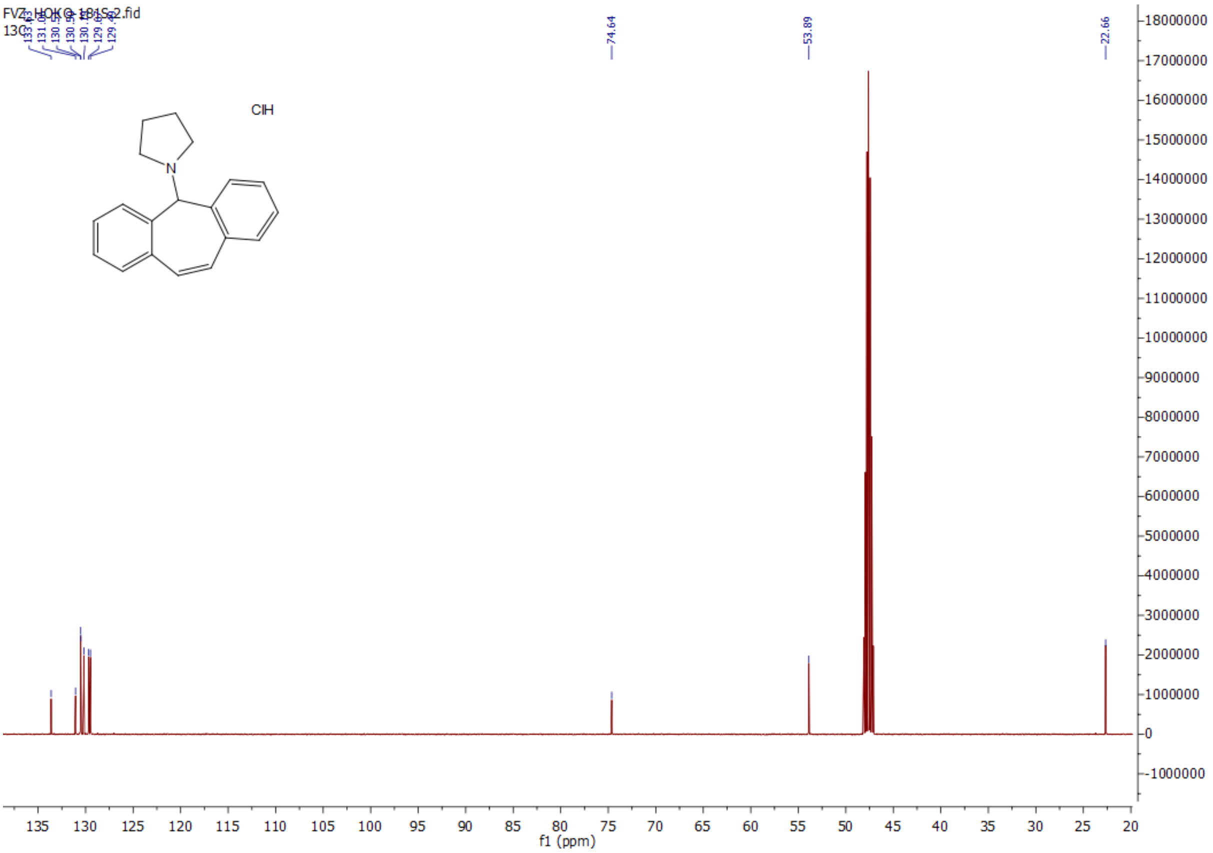

2-((5*H*-dibenzo[*a,d*][7]annulen-5-yl)amino)ethanol hydrochloride (**6l**)

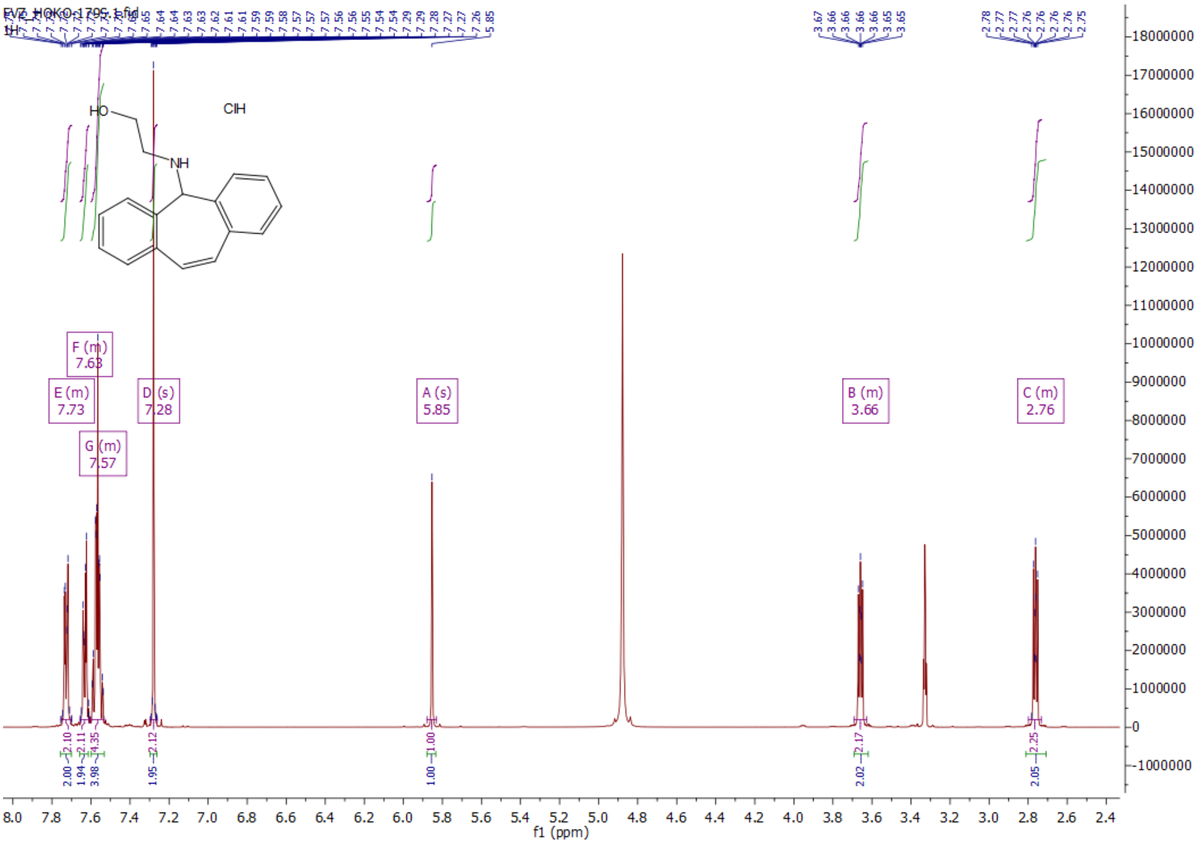

2-((5*H*-dibenzo[*a,d*][7]annulen-5-yl)amino)ethanol hydrochloride (**6l**)

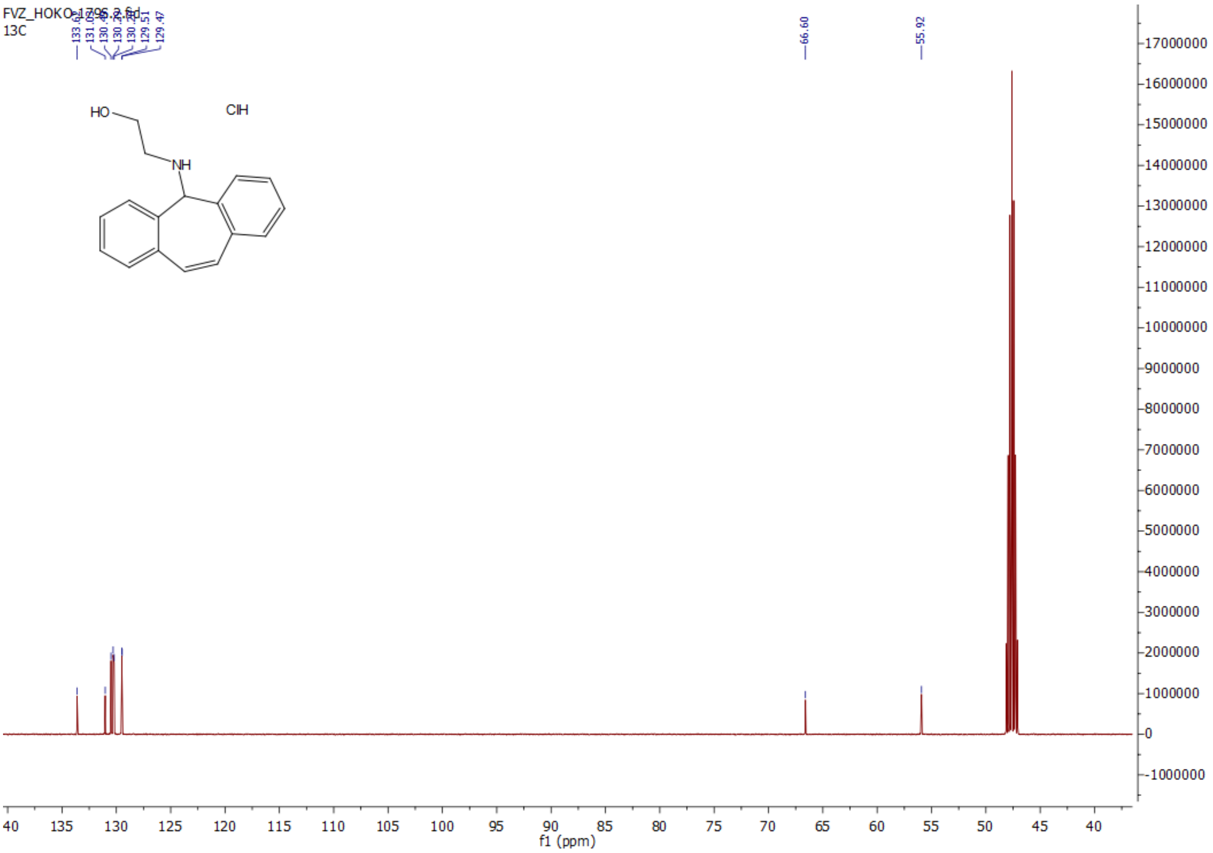
